# Assembling and annotating the Tainung 67 rice genome to trace its *japonica* and *indica* ancestries

**DOI:** 10.64898/2026.07.15.738793

**Authors:** Jerome P. Panibe, Long Wang, Tzi-Yuan Wang, Chang-Sheng Wang, Mei-Yeh Jade Lu, Wen-Hsiung Li

## Abstract

Here we reported the 409.0 Mb assembly of the Tainung 67 (TNG67) genome, an early hybrid of *japonica* and *indica* cultivars. We used different platforms to sequence the TNG67 genome with a total coverage of 209.8x, from a combination of Illumina paired-end reads, Illumina mate-pair reads, and Oxford Nanopore Technology long reads. The assembly has an N50 of 32.3 Mb with the longest scaffold = 45.2 Mb. We have annotated 38,938 genes where 28,376 (72.9%) finished with the Blast2GO annotation stage. There were 4,821 genes, (98.4%) of the total gene count that have complete BUSCOs. 48.33% (197.5Mb) of the TNG67 genome is composed of repeats. We predicted a total of 527 blast-resistant genes in TNG67. We also analyzed a total of 20 grain size genes and 38 photoperiod-related genes of TNG67, Nipponbare and TN1, and grouped them in terms of whether the TNG67 genes are more similar to Nipponbare, more similar to TN1, hybrid, same, or unique. We also determined whether the sequences of the TNG67 genome were derived from its *japonica* or *indica* ancestors. This TNG67 genome may help rice researchers improve yield, develop resistance against biotic and abiotic stress, and understand the evolution of a hybrid cultivar of *japonica* and *indica*.

## Background

Tainung 67 (TNG67) is an in-demand cultivar for rice research in Taiwan. This elite *japonica* rice was extensively planted from 1979 to 1998 and has been used as a parent to breed many rice cultivars in Taiwan mainly for two reasons. First, TNG67, despite being a *japonica* cultivar, exhibits photoperiod insensitivity, making it suitable to grow in Taiwan, which has long days in summer. Second, it is a high-yielding cultivar. TNG67 is a *japonica* because its direct parents are Tainung 61 and a descendant of the cross of Taichung Shih 138 and Tainung 61 (see Figure 1). All of these three cultivars are of *japonica* type. Its other indirect *japonica* parents are Taichung 65, Taichung 178, Taichung Jiaocai No. 15, plus some unknown cultivars (see Figure 1). However, it possesses *indica* genetic material from TN1 (Taichung Native 1) (Hour et al. 2007) (see Figure 1), enabling it to flower under long days (i.e., photoperiod insensitive). TNG67 is also described as lodging resistant, possibly due to the *sd1* (semi-dwarf) gene inherited from TN1, though no direct evidence exists. Moreover, TNG67 is highly susceptible to the blast pathogen (Wang et al. 2019), making it a model rice for blast disease studies.

**Figure 1.**
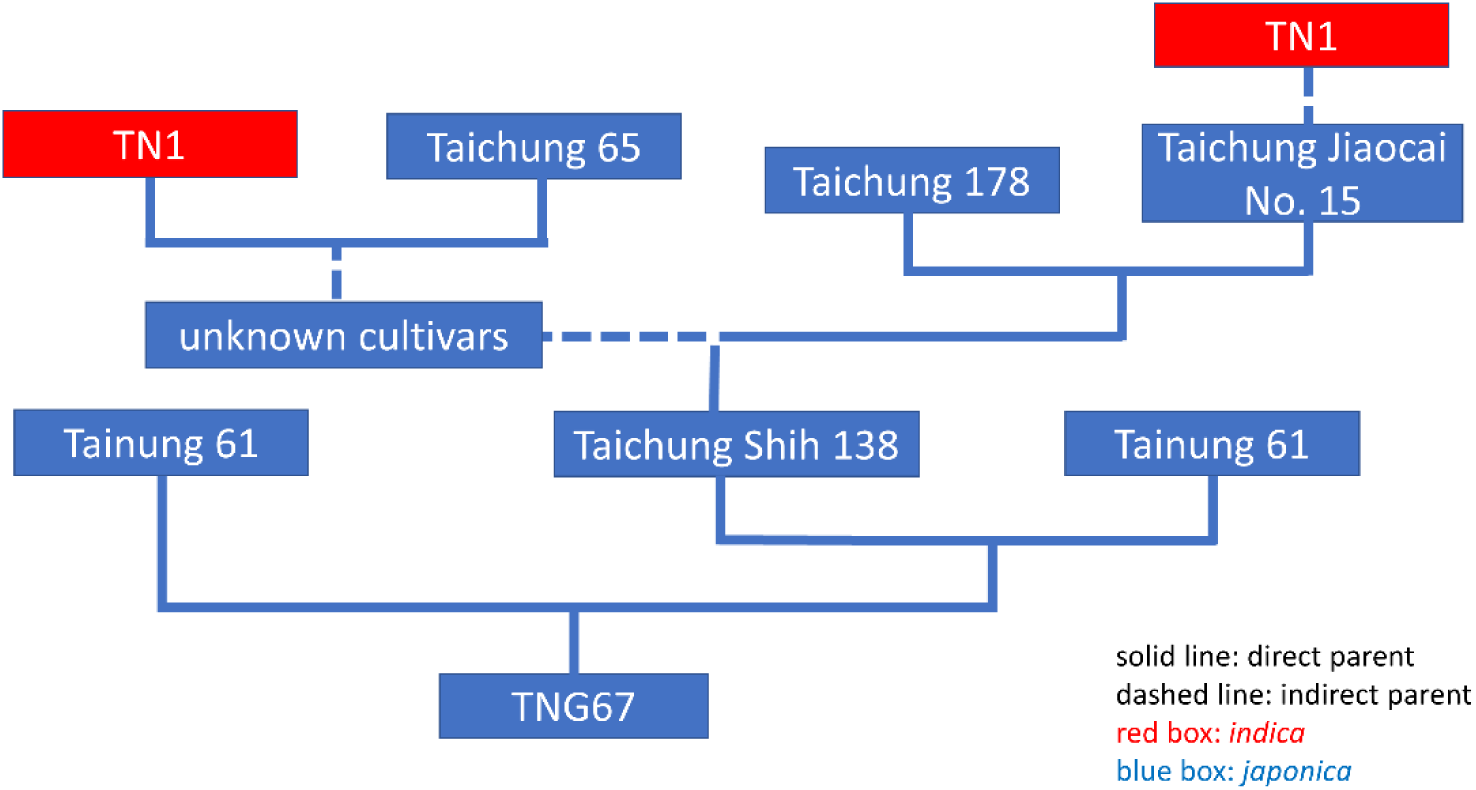
A simplified pedigree of TNG67. TNG67 was derived primarily from a cross between Tainung 61 and the offspring of the cross of Taichung Shih 138 and Tainung 61. The *indica* TN1 cultivar is an indirect parent of TNG67 and can be traced through the ancestry of Taichung Shih 138, a *japonica* cultivar. (Hour et al. 2007)

Many studies have been made on TNG67, including the following. Lin et al. (2014) identified an amino acid mutation in the OsGAPDHB (β subunit of glyceraldehyde-3-phosphate dehydrogenase) protein that affected the fragrance characteristic of the cultivar (Lin et al., 2014). The transcriptome of TNG67 was profiled to study its gene expression in response to low temperatures. (Yang et al., 2015) Resistance of TNG67 against the fungus *Fusarium fujikuroi* was also investigated, and compared against the susceptible *Zerawchanica karatals* (ZK) cultivar (Cheng et al, 2020). Response to the fungus was related to jasmonic acid (JA) signaling pathways (Cheng et al, 2020). A pool of 3000 TNG67 mutant strains was also developed using sodium azide with varying responses to the fungal pathogen *Magnaporthe oryzae* (Wang et al., 2019). The SNP rate between TNG67 and Nipponbare was also estimated at 0.3% to 0.4% (Hour et al., 2007). Two BAC libraries were made in 2005 to facilitate the analysis of genes as well as for structural and functional studies. The first BAC library has an average insert size of 138.4 kb and coverage of 15.1x, while the second one has an average insert size of 137.8 kb and a coverage of 3.2x (Lin et al., 2006).

Many rice genomes have assembled recently using long read sequencing ((Lu et al. (2022), Panibe et al. (2021), and Zhou et al. (2020)). One of these rice genomes is TN1 (Panibe et al., 2021), an indirect *indica* parent of TNG67. In view of the importance of TNG67 in rice genetics research, it is high time that its genome be assembled and characterized. It will facilitate studies not only on itself but also on other rice varieties, especially those that used TNG67 as a direct or indirect parent.

We therefore will pursue the following objectives: (1) to sequence and assemble the TNG67 genome using a combination of short and long-read sequencing technologies, (2) to determine whether specific segments of the TNG67 assembly predominantly originate from *japonica* or indica, (3) to identify blast resistance genes in TNG67 and to provide plausible explanations for its high susceptibility to blast disease, (4) to investigate the reasons behind TNG67’s photoperiod insensitivity (PI), despite being a ‘*japonica*’ cultivar, and (5) to study why TNG67’s grain size is similar to that of *japonica* cultivars. Completing the genome assembly and decoding the genes of TNG67 will provide an invaluable resource for rice researchers in Taiwan and globally.

## Results

### TNG67 genome assembly

The nanopore reads were assembled by Canu v2.2.1 (see Figure 2), producing a 404.3 Mb assembly composed of 2,708 contigs with an N50 of 474.6 kb (see Table S1). This Canu primary assembly was polished by Pilon for one run. By using various Illumina long paired-end and mate-pair reads, SSPACE turned the 2,708 contigs into 2,209 scaffolds with the longest one measuring 2.6 Mb. Gaps were formed after using SSPACE and were closed by PBJelly using the corrected the nanopore long reads, reducing the total gap length from 227,888 bp to 199,012 bp. The remaining gaps were reduced further by GapCloser and GapFiller (see Figure 2), using paired-end reads. After using GapFiller, the TNG67 assembly had a total length of ∼409.3 Mb, an N50 of 539.5 kb and a gap length 114,876 N’s. At this point, the TNG67 assembly still had 2,206 scaffolds. To form the chromosome-level scaffolds, we did a RagTag reference-guided assembly using the Nipponbare reference genome produced by IRGSP and polished the genome with one round of Pilon correction. After uploading to NCBI for submission, contaminant contigs were detected and any gaps that they sandwiched removed from the assembly. The contaminant contigs were unassigned sequences and removing them would improve the genome assembly. A total of 125,190 bp of sequences were removed.

**Figure 2.**
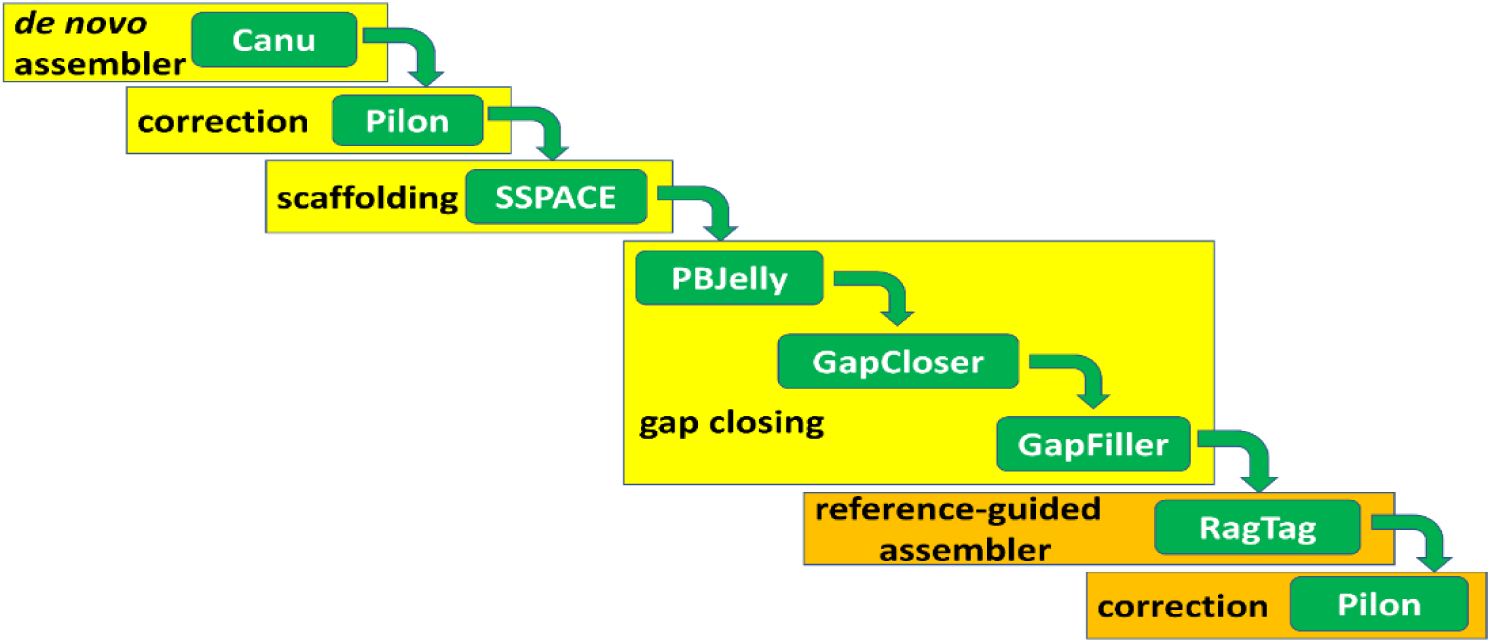
Workflow of the de novo assembly of TNG67. Yellow boxes denote de novo assembly part; orange boxes denote reference-guided assembly; and green boxes indicate software/tool used.

To further improve the genome assembly, we did another round of RagTag reference-guided assembly. This time we used the Nipponbare T2T genome assembly (Shang et al. 2020). We used the original sequences that were produced by GapFiller minus the contaminant sequences. The RagTag re-assembly inserted a gap of 100 N’s to link the adjacent scaffolds, increasing the total length of gaps to 258,012 bp. One round of Pilon correction reduced theTNG67 assembly to size of 408,946,873 bp with an N50 of 32,263,301 Mb. After uploading to NCBI submission, the assembly was flagged with contaminants again and we redo the process of RagTag assembly producing a final assembly with a size of 409.0 Mb with an N50 of 32.3 Mb.

### Comparison of the genome assemblies of TNG67, Nipponbare and TN1

The genome assembly statistics of TNG67, TN1 and Nipponbare T2T are compared in Table 1. TNG67’s genome size (409.0 Mb) is more similar to that of TN1 (409.5 Mb) than to that of Nipponbare T2T (385.7 Mb) (Shang et al., 2023), which is somewhat larger than the International Rice Genome Sequencing Project (IRGSP) assembly (Genbank accession #: GCA_001433935.1) (374.4 Mb). The assemblies of TNG67, TN1 and Nipponbare T2T have 15, 20 and 12 scaffolds of length > 100 kb, and all of them have 12 scaffolds > 1 Mb (Table 1), which are the 12 chromosomal length scaffolds (see Table 2). When their longest scaffolds (chromosome 1’s) are compared, TN1(45.5 Mb) > TNG67 (45.2 Mb) > Nipponbare T2T (43.9 Mb) (see Table 2). Their N50’s follow the same order: TN1 (33.1 Mb) > TNG67 (32.3 Mb) > Nipponbare T2T (31.1 Mb) (see Table 1). However, when their N90’s are compared, Nipponbare T2T (27.5 Mb) > TN1 (26.6 Mb); TNG67 (26.6 Mb). TNG67 has the largest number of N’s in its assembly, with 274,812 Ns distributed in 1,786 gaps, compared to TN1’s 128,560 Ns in 1,076 gaps and Nipponbare T2T’s gapless assembly. A total of 38,938 genes were predicted in the TNG67 genome, higher than TN1’s 37,526 genes and Nipponbare T2T’s 57,359 genes (Table 1). Although TNG67 has more predicted genes than do TN1 and Nipponbare, it has slightly fewer complete BUSCOs (4,821), at a rate of 98.4% compared to 98.7% in TN1 and 98.6% in Nipponbare T2T. The amounts of repeat sequences are TN1 (210.9 Mb) > TNG67 (197.5 bp Mb) > Nipponbare T2T (188.1 Mb). About 48.33% of TNG67’s genome is composed of repeats, compared to 51.52% of TN1 and 48.76 % of Nipponbare T2T.

**Table 1.** Comparison of the genome assemblies of TNG67, TN1 and Nipponbare. To have a fair comparison against the TNG67 and TN1 (Panibe, et al. 2021) assemblies, the Nipponbare T2T genome (Genbank accession #: GCF_034140825.1) used here contains its unplaced scaffolds.

|  | TNG67 | TN1 | Nipponbare<br>T2T |
| --- | --- | --- | --- |
| Total sequence<br>length | 408,959,857 | 409,523,013 | 386,337,859 |
| Number of<br>scaffolds (>100<br>kb) | 15 | 20 | 14 |
| Number of scaffolds (> 1 Mb) | 12 | 12 | 12 |
| Longest scaffold | 45,156,375 | 45,524,768 | 43,929,697 |
| L50 | 6 | 6 | 6 |
| L75 | 9 | 9 | 9 |
| N50 | 32,306,537 | 33,082,646 | 31,111,469 |
| N90 | 26,640,693 | 26,568,179 | 27,474,823 |
| # N's / # gaps | 274,812 / 1,786 | 128,560 / 1,076 | 1 |
| # N's per 100 kbp | 67.20 | 31.39 | 0 |
| GC% | 43.45 % | 43.82 | 43.72 |
| Number of genes | 38,938 | 37,526 | 37,483 |
| mean gene length | 4,165 | 4,057 | 3,496 |
| mean CDS length | 1,226 | 1,204 | 1,415 |
| complete BUSCO | 4,821 (98.4%) | 4,830 (98.7%) | 4,828 (98.6%) |
| Repeat sequence (bp) | 197,518,184 bp<br>(48.33 %) | 210,923,132<br>( 51.52 %) | 188,090,818<br>(48.76 %)* |

\*based on the 12 chromosomes of Nipponbare T2T

**Table 2.**
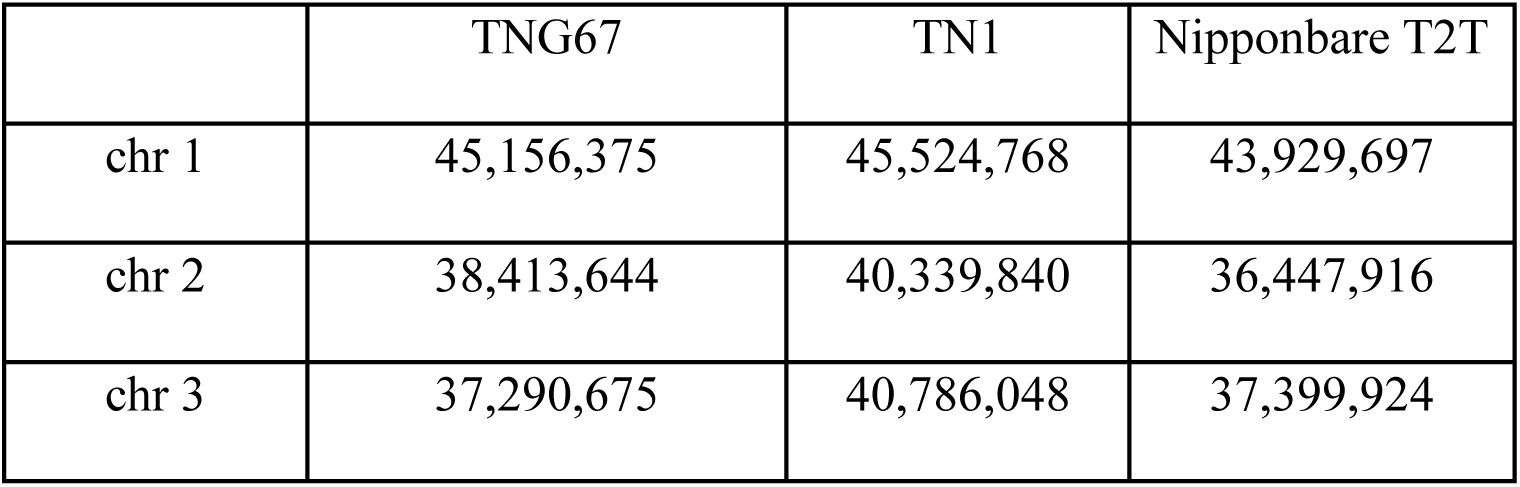
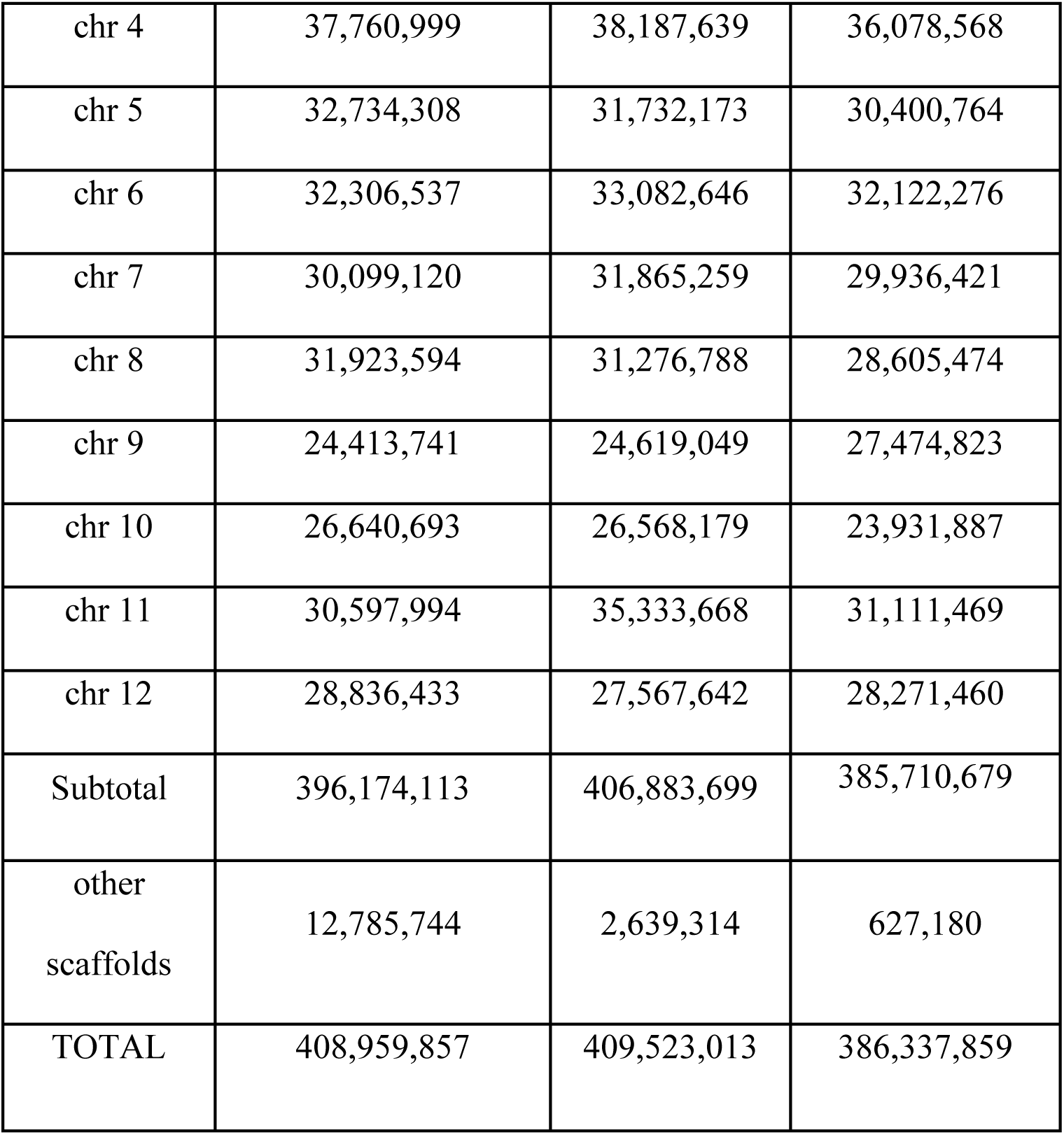
Chromosome lengths (bp) of the genomes of TNG67, TN1 and Nipponbare.

|  | TNG67 | TN1 | Nipponbare T2T |
| --- | --- | --- | --- |
| chr 1 | 45,156,375 | 45,524,768 | 43,929,697 |
| chr 2 | 38,413,644 | 40,339,840 | 36,447,916 |
| chr 3 | 37,290,675 | 40,786,048 | 37,399,924 |
| chr 4 | 37,760,999 | 38,187,639 | 36,078,568 |
| chr 5 | 32,734,308 | 31,732,173 | 30,400,764 |
| chr 6 | 32,306,537 | 33,082,646 | 32,122,276 |
| chr 7 | 30,099,120 | 31,865,259 | 29,936,421 |
| chr 8 | 31,923,594 | 31,276,788 | 28,605,474 |
| chr 9 | 24,413,741 | 24,619,049 | 27,474,823 |
| chr 10 | 26,640,693 | 26,568,179 | 23,931,887 |
| chr 11 | 30,597,994 | 35,333,668 | 31,111,469 |
| chr 12 | 28,836,433 | 27,567,642 | 28,271,460 |
| Subtotal | 396,174,113 | 406,883,699 | 385,710,679 |
| other<br>scaffolds | 12,785,744 | 2,639,314 | 627,180 |
| TOTAL | 408,959,857 | 409,523,013 | 386,337,859 |

### Predicting blast resistance genes in TNG67

R (resistance) genes encode proteins that confer plants a natural defense against the blast pathogen. There are several types of R genes and the most common are the NB-LRR (nucleotide-binding leucine-rich repeat) genes composed of the NB-ARC (nucleotide-binding adaptor shared by APAF-1, R proteins, and CED-4) domain and leucine-rich repeat (LRR). Sometimes an R gene is composed of only the NB-ARC domain and is said to be incomplete. Having both the NB-ARC and LRR domains makes an R gene complete.

The combined result of Pfam and NLR-Parser predicted a set of 537 R proteins in TNG67 (see Table S2). These 537 R proteins included 526 R genes (some R proteins are isoforms of the same R gene) and OsTNG11t000869, which has both an NB-ARC (OsTNG11t000869.1) and an NBS-LRR (aka NB-LRR) (OsTNG11t000869.2) domain for its isoforms. This makes a total of predicted 527 R genes in the TNG67 genome (see Figure S1). Furthermore, 145 of them only have the NB-ARC domain, while 379 genes also have the NBS-LRR domain, of which 378 are found in the chromosomal-level scaffolds. The remaining R genes are neither classified as NB-ARC nor NB-LRR. The distribution of the NBS-LRR R genes among the chromosomes of TNG67 is in Figure S2. The shortest Chromosome 11 having more R genes than the longest Chromosome 1.

Using the list of cloned R genes (Mahesh et al. 2016), we determined if a given R gene is present or absent in TNG67. The two R genes, *Pik-m-TS2* and *Pik-2,* are from *japonica* varieties. The *indica* R genes *Pi5-1*, and *Pi25* and the *japonica* R genes *Pi54(Pik-h)*, *Pid2*, and *Pid3* are missing in the TNG67 genome. Moreover, in the TNG67 genome, the following R genes *pi21*, *Pib*, *Pi-ta*, *Piz-t*, *Pi37*, *Pik-m-TS1*, *Pit*, *Pi5-2*, *Pb1*, *Pish*, *Pia(RGA4)*, *Pik-p-1*, *Pik-p-2*, *Pik-1*, *Pi64* were all originated from the *japonica* cultivars and some were found to harbor mutations. The following R genes, *Pi9*, *Pi36*, *Pi1-5*, and *Pi1-6*, were from *indica* and the mutated R gene *Pi54rh* originated from a wild rice species.

### Blast2GO annotation of TNG67 genes

We used the free version of the Blast2GO software for the functional annotation of the TNG67 genes. The fasta gene sequences were blasted via blastx to the nr database. The resulting blast results were saved as an xml file, which was used as the input in Blast2GO: 1) to perform mapping to tag the genes with GO terms, and then 2) to get the final set of functionally annotated TNG67 genes. The top 3 hit species were *Oryza sativa japonica* group (28,843 hits), *Oryza sativa* (8,730 hits), and *Oryza sativa indica* group (6,462 hits). With regards to the data distribution of the Blast2GO analysis of the TNG67 genes, 28,607 (73.47%) genes of the input 38,936 genes finished the annotation stage. A total of 1,562 genes (4.01%) did not have any blast hits, while 7,251 (18.62%) genes did not have any GO mapping. Of those genes that had inferred GOs, 1,516 (3.89%) genes only finished at the mapping stage and did not reach the annotation stage.

### Analysis of grain size genes of TNG67, Nipponbare and TN1

We checked the gene structures of grain-size genes. *Indica* varieties are characterized as having long grains, as opposed to shorter and round grains in *japonica* cultivars. By inspecting the different grain size genes as reported by Jiang et al. (2022), we could identify differences and similarities among the grain size genes of TNG67, Nipponbare, and TN1. From the list of genes in Table 3, we conducted an orthologue analysis of the proteomes of the three cultivars. Using the gene names, we searched the RAP-DB (Sakai et., 2013) to determine the specific Nipponbare proteins, and from the results of an OrthoFinder run, we identified their orthologues in TNG67 and TN1.

**Table 3.**
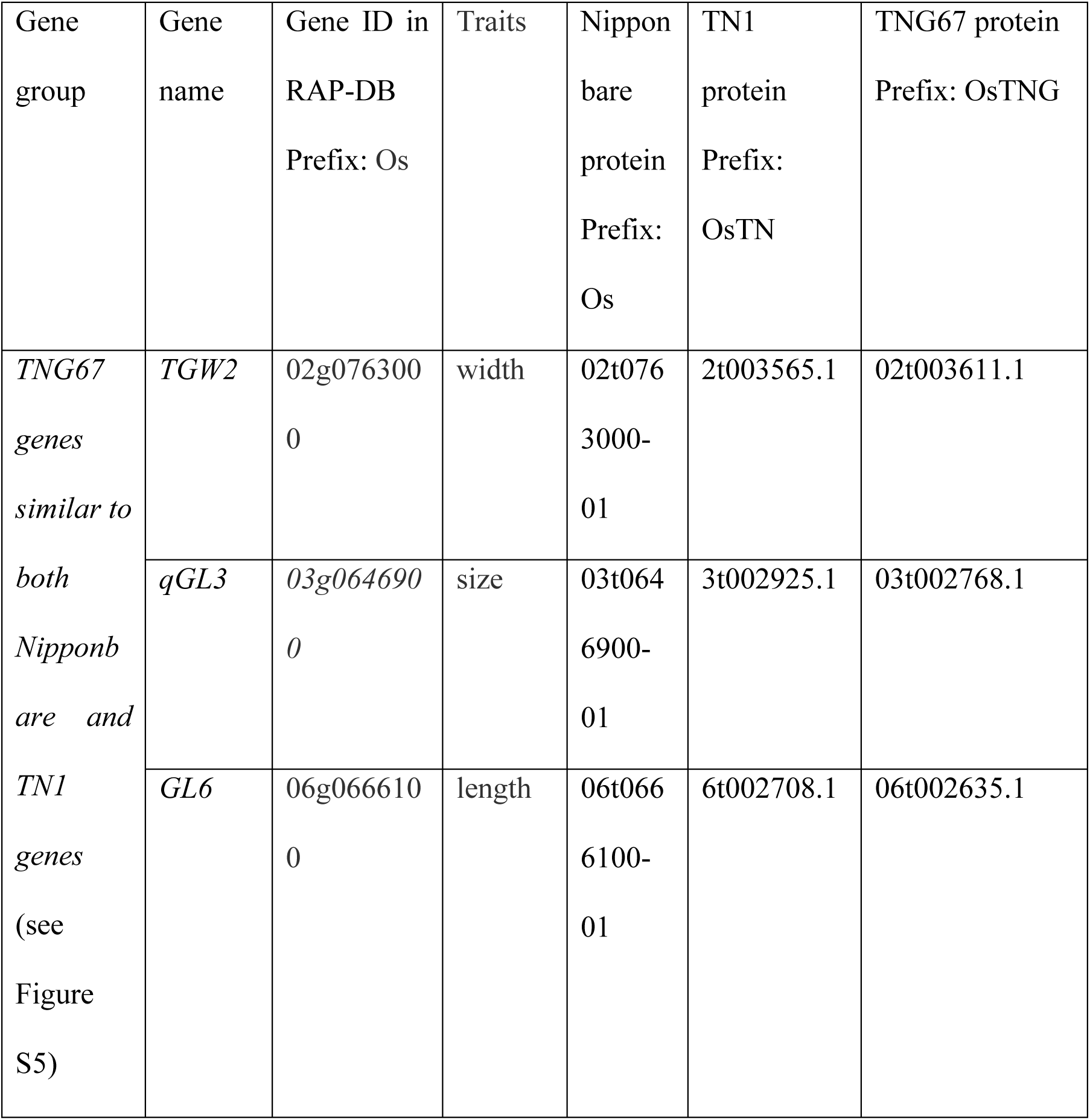
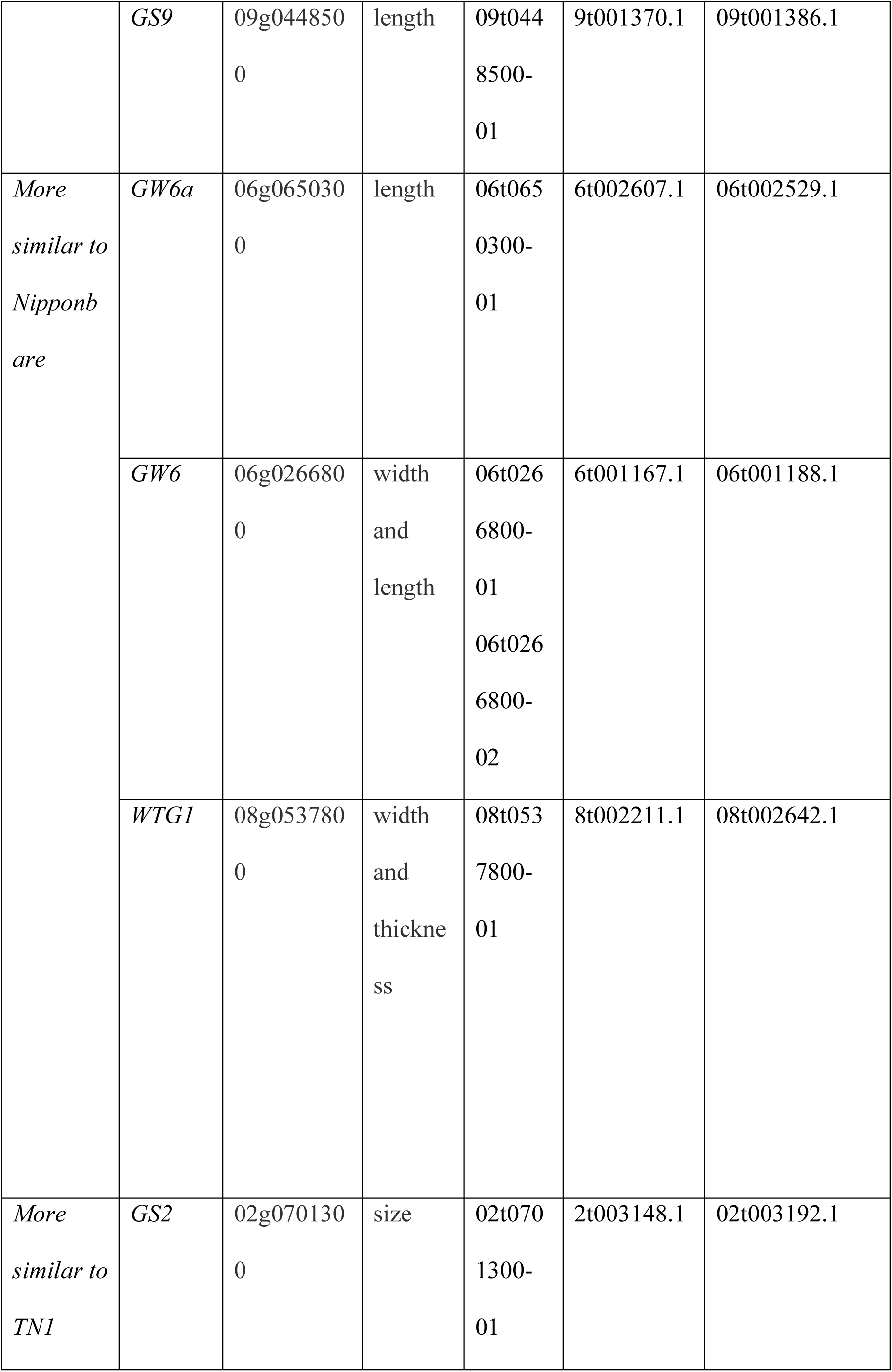
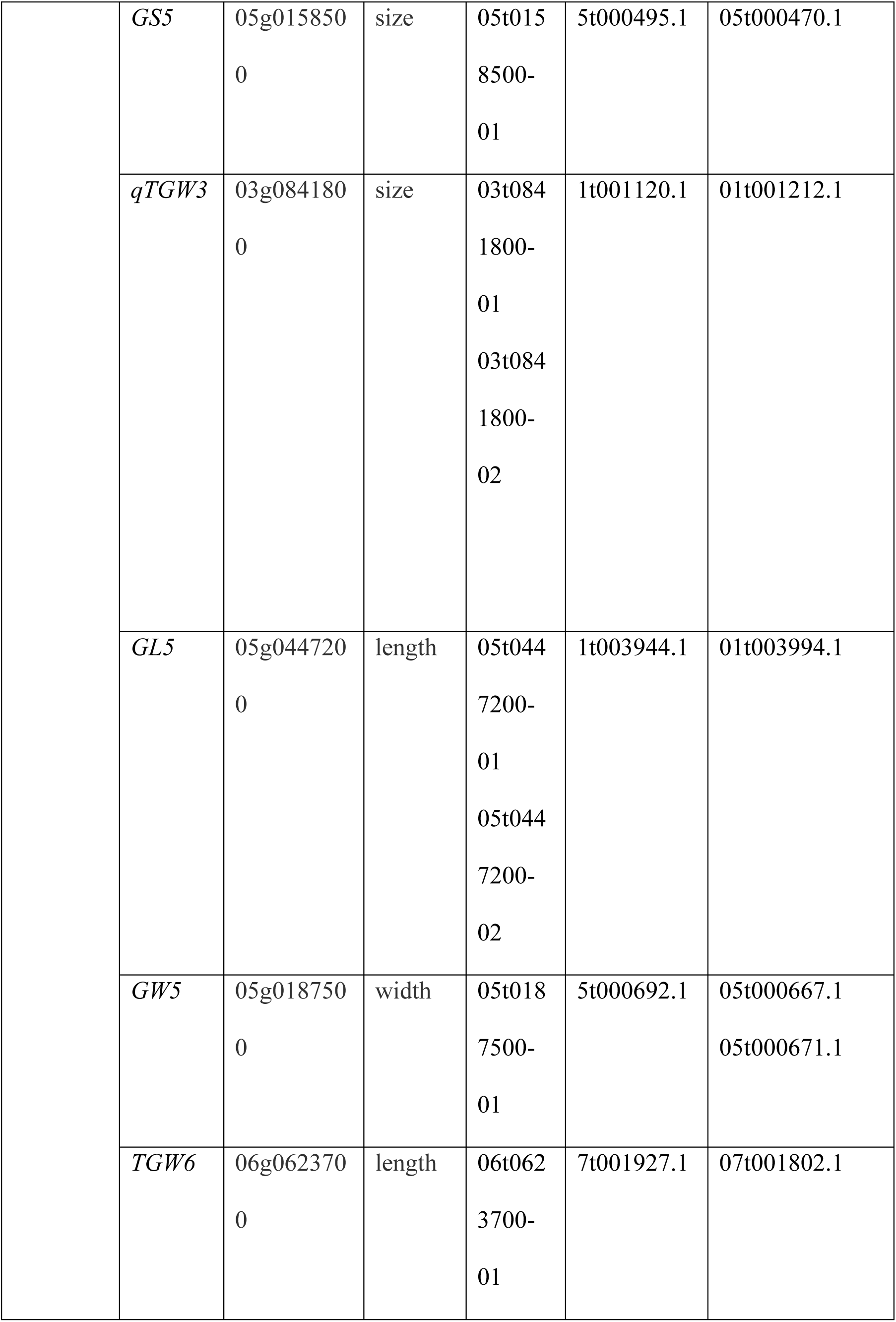
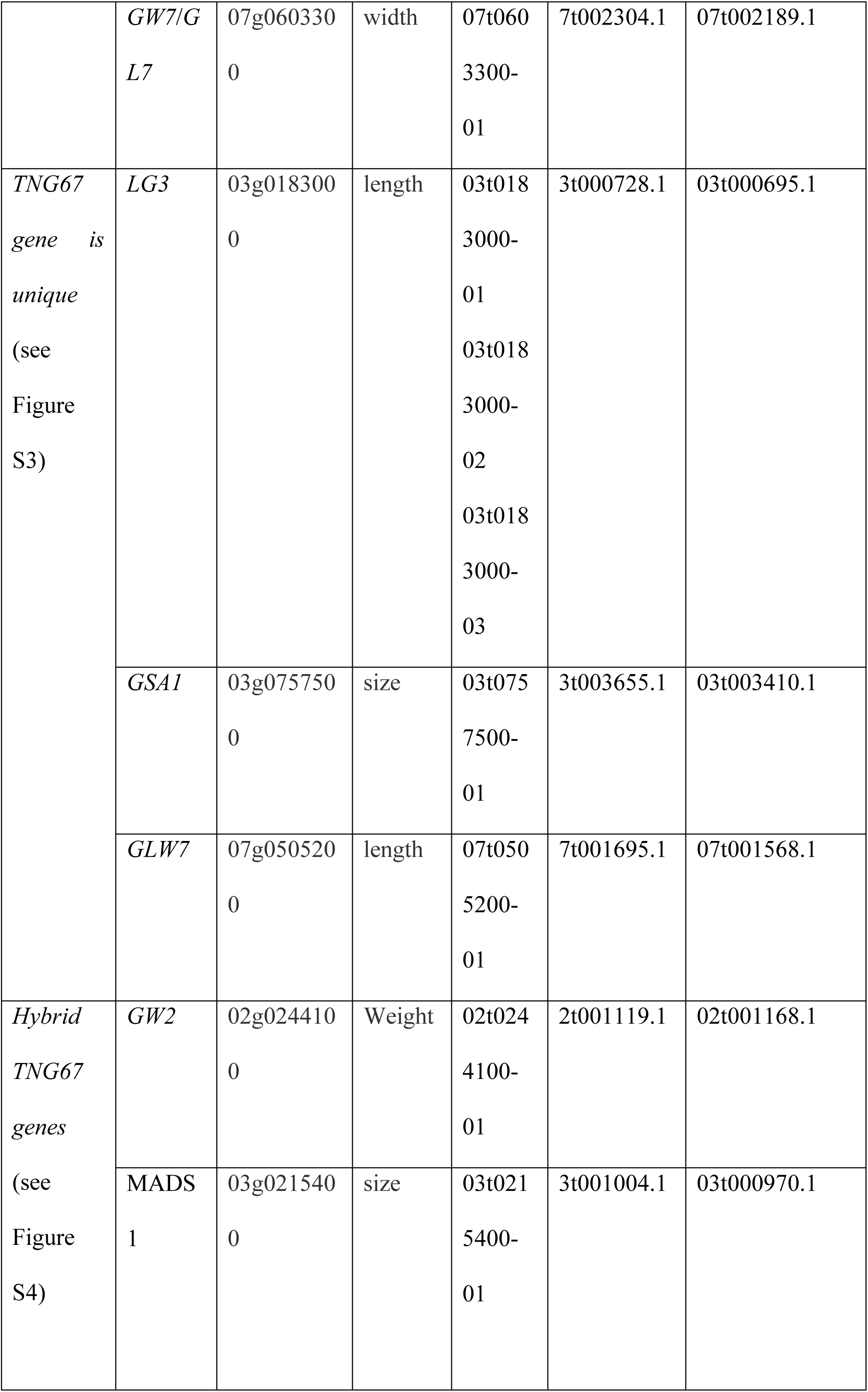
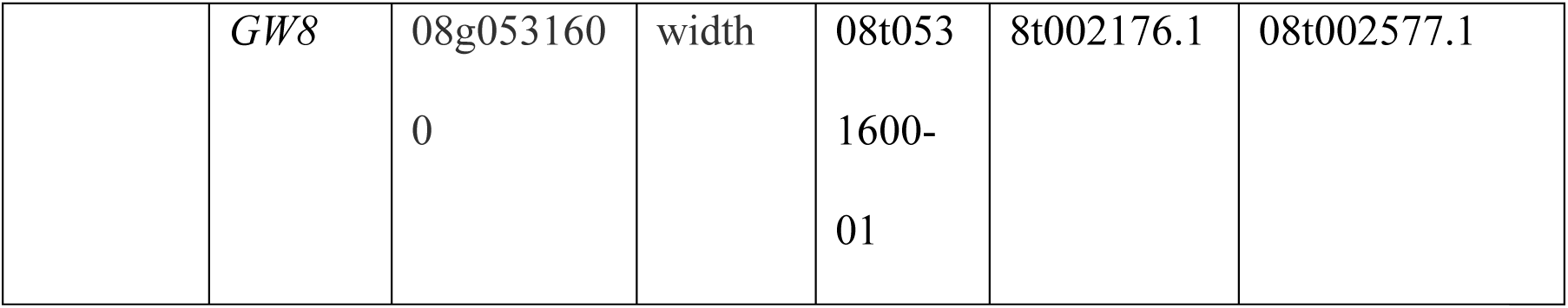
Characterization of common grain size genes in the three rice cultivars. For each gene, the gene ID was searched in RAP-DB to identify the Nipponbare protein. Gene names: *TGW2* (thousand grain weight 2); *qGL3* (grain length 3.1); *GL6* (short grain 6/ grain length 6); *GS9* (grain shape gene on chromosome 9); *GW6a* (grain weight on chromosome 6-a); *GW6* (grain width 6); *WTG1* (wide and thick grain 1); *GS2* (Grain size 2); *GS5* (grain size 5); *qTGW3*; (thousand grain weight 3); *GW5* (Grain width 5); *TGW6* (total grain weight 6 / thousand-grain weight 6); *GW7/GL7* (grain length on chromosome 7, grain width QTL on chromosome 7); *GSA1* (grain Size and Abiotic stress tolerance 1); *GLW7* (grain length and weight on chromosome 7); *GW2* (Grain weight 2); *MADS1* (MADS box gene1); *GW8* (grain-width 8).

To visualize the gene structures, we used the online tool GenePainter. Inputs were the Clustalw alignment of protein orthologues and a gff file containing information about the gene and its associated protein isoforms. From the GenePainter diagrams, we classified the TNG67 genes into five groups: whether a TNG67 gene is more similar to TN1 than to Nipponbare, is more similar to Nipponbare than to TN1, is unique (see Figure S3), or is a hybrid (similar to both Nipponbare and TN1 in certain parts) (see Figure S4); or the three cultivars have highly similar gene structures (see Figure S5). We say that the gene structure of TNG67 is unique if one or more parts of the TNG67 gene show no similarity to Nipponbare and TN1. Sometimes, a Nipponbare gene can have more than one protein isoform, and we only picked the best representation with respect to its completeness of exonic and intronic regions.

As TNG67 is described phenotypically as a *japonica* cultivar having short and round grains, we expect the grain size genes of TNG67 to be similar to those of Nipponbare. Surprisingly, the following seven TNG67 genes are more similar to TN1 than to Nipponbare (see Table 3): *GS2* (grain size of chromosome 2), *GS5* (grain size 5), *qTGW3* (thousand grain weight 3), *GL5*, *GW5* (grain width 5), *TGW6* (thousand-grain weight 6), and *GW7/GL7* (grain width QTL on chromosome 7/ grain Length on chromosome 7). The following three TNG67 genes are more similar to Nipponbare: *GW6a* (grain weight on chromosome 6-a), *GW6* (grain width 6), and *WTG1* (wide and thick grain 1). Moreover, the following four TNG67 genes are unique with respect to TN1 and Nipponbare: *LG3*, *GSA1* (grain size and abiotic stress tolerance 1), *GLW7* (grain length and weight on chromosome 7), and *GW8* (grain-width 8). There are two cases (*GW2* and *LGY3*) where the TNG67 gene is a hybrid. Finally, there are four TNG67 genes that are similar to both TN1 and Nipponbare genes: *TGW2* (thousand grain weight 2), *qGL3* (grain length 3.1), *GL6* (short grain 6/ grain length 6), and GS9 (grain shape gene on chromosome 9). For the *GS3* (grain size 3) gene, no orthologue was found in TNG67 or in TN1.

We investigated why there are TNG67 genes more similar to TN1 than to Nipponbare, given that TNG67has round and short grains just like any typical *japonica* cultivars. Furthermore, there are 7 genes in which TNG67 and TN1 have similar gene structures (see Table 3), while there are only 3 TNG67 genes that are more similar to Nipponbare (see Table 3). We suspected that some differences in exonic regions between TNG67 and TN1 have no effect on the resulting protein sequences, because the mutations were not in important protein regions. To test our hypothesis, we checked the domains of the TNG67-TN1 similar genes and the TNG67-Nipponbare similar genes. We used hmmscan and the Pfam v35.0 database. To search for possible mutations, we used the data from the RiceVarMap v2.0 database (ref), which contains data from the 3,000 Rice Genomes Project, which includes data from TNG67, Nipponbare and TN1 cultivars. We also did a sequence alignment of the proteins using ClustalW. This determined which portions of the alignment correspond to the alignment of the exonic regions (orange boxes) in the gene structures, and pinpointed the gaps in the alignment that correspond to the loss of exonic regions (yellow boxes) in the gene structure diagrams. We first inspected the alignments in Figure 3, where the TNG67 and Nipponbare genes are more similar. For GW6a, the protein sequences of the two cultivars are identical, while that of TN1 has an insertion of two amino acids (KT) creating an insertion in the first exon (Figure 3a) and an amino acid change from serine to glycine. The serine to glycine (S to G) change is radical because it is from a polar amino acid to the smallest non-polar amino acid. Moreover, the KT insertion is also radical because K (lysine) is positively charged. Thus, these two changes could have changed the property of the GW6a protein in TN1. For GW6, Nipponbare has two isoforms and we selected the second isoform (Os06t0266800-02) as the representative protein of Nipponbare GW6 because of the completeness of its exonic regions. Aligning this Nipponbare protein to its TNG67 and TN1 orthologues reveals amino acid changes in the first exon of the TN1 gene (Figure 3b). In addition, the second exon of the TN1 *GW6* gene showed an R (arginine) to H (histidine) change. This change could affect the protein structure of the TN1 GW6 protein because histidine has a ring structure, whereas arginine has a long side chain. Another difference in the TN1 GW6 protein is that it has an insertion of two serine amino acids (SS) in the second exon of the TN1 gene. Thus, the *GW6* gene structure of TNG67 is highly similar to that of Nipponbare but different from that of TN1 (see Figure 3b). For WTG1, the TN1 protein shows two insertions of amino acid sequences when compared to the WTG1 protein sequences of TNG67 and Nipponbare, which are highly similar (see Figure 3c)

**Figure 3.**
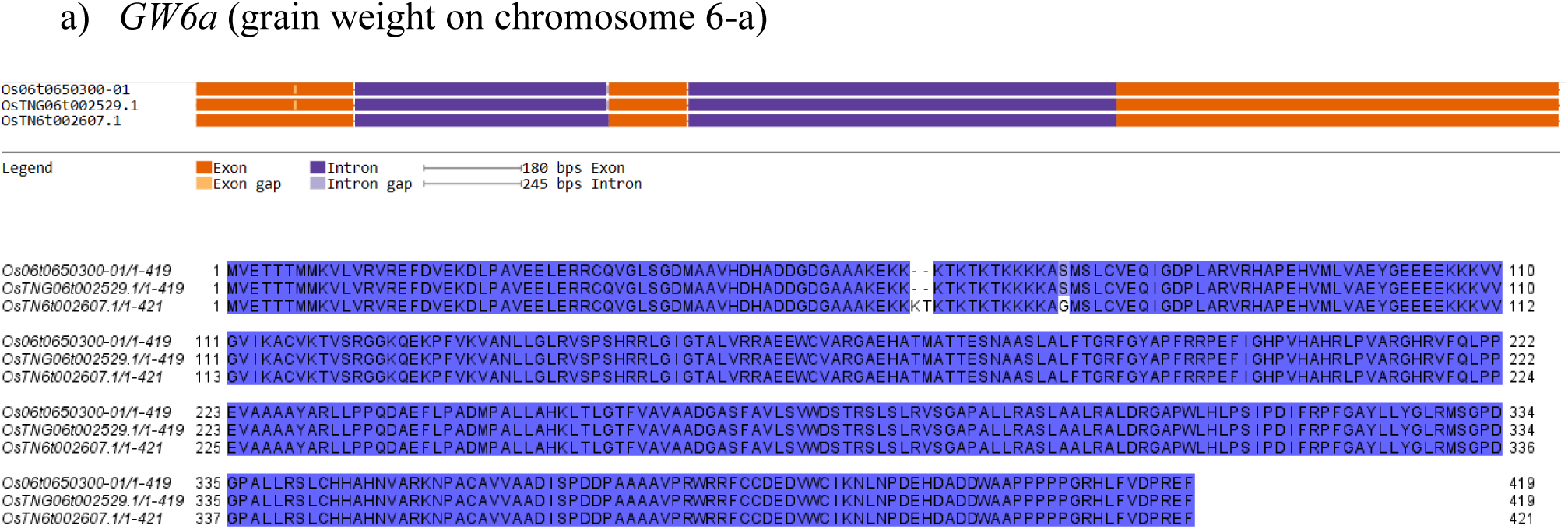
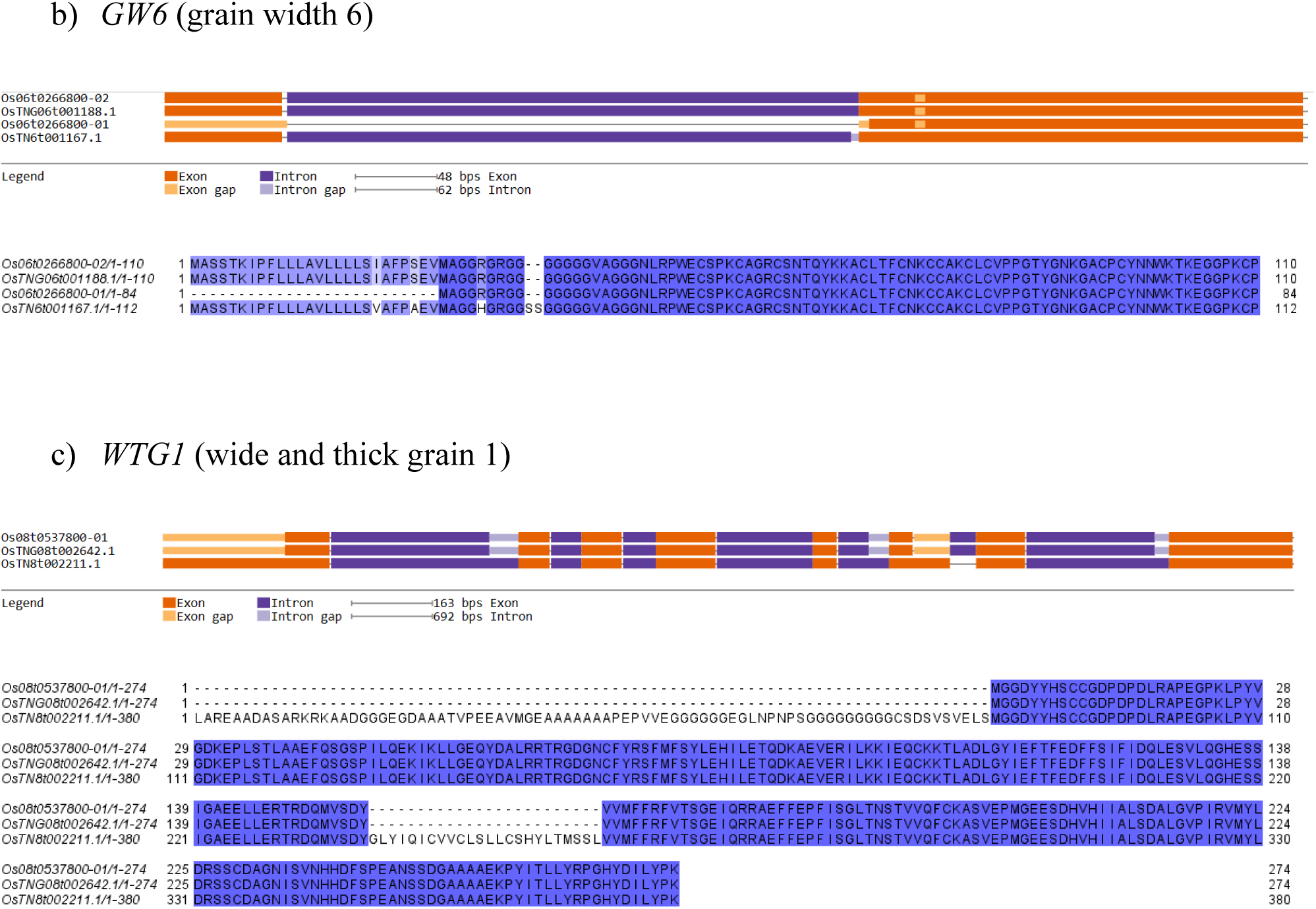
Structures and protein sequence alignments of grain size genes for the cases wherein TNG67 and Nipponbare are more similar to each other than to TN1. In the alignment, dark blue signifies 100% identity, and lighter blue indicates 50-100% identity. Dashes represent gaps. Losses of exonic regions in gene diagrams correspond to these gaps, while 100% sequence alignment aligns with the gene diagrams’ orange bars. Amino acid changes in protein sequences are evident only in the sequence alignment. a) GW6a: TNG67 and Nipponbare are similar, while the protein sequence of TN1 GW6a shows an insertion of KT amino acids and a change from S (serine) to G (glycine). b) GW6: TNG67 and the Nipponbare second isoform are similar to each other. The TN1 GW6 protein sequence has a change from isoleucine (I) to valine (V), a change from serine (S) to alanine (A), and a change from argininine(R) to histidine (H) and an insertion of two serines (SS). c) WTG1: TNG67 and Nipponbare are similar while TN1 shows two insertions of amino acid sequences.

Some genes show a structural resemblance between TNG67 and TN1. However, an inspection of their protein sequence alignments reveals differences between TNG67 and TN1. In the *GS2* gene (see Figure 4a), two amino acid changes (A to T and G to V) are found in TN1 with respect to the TNG67 protein OsTNG02t003192.1. In the former, the change is from a nonpolar amino acid (A) into a polar one (T), while in the latter the methyl group of valine occupies more space than the lone hydrogen R-group of glycine. For the *GS5* gene, TNG67’s OsTNG05t000470.1 and TN1’s OsTN5t000495.1 show similar gene structures, except for small losses of exonic regions, which are shown as gaps in their protein sequence alignment (Figure 4b). Compared to TNG67 and Nipponbare, TN1 shows M to V, V to A, E to D and NQ to TK changes. In addition, there is a K to Q difference between TNG67 and TN1 (see Figure 4b). These amino acid changes indicate that the TNG67 GS5 is different from the TN1 GS5. For the *qTGW3* gene, we choose Os03t0841800-01 as the representative isoform of Nipponbare (Figure 4c), and it shows the loss of a region in its first exon. However, TN1’s OsTN1t001120.1 has lost the entire 1^st^ exon. Hence, the *qTGW3* gene of TNG67 and that of TN1 are different. For the *GL5* gene, the complete first exon in TN1 makes it different from the partial first exons of Nipponbare and TNG67 (see Figure 4d). For *GW5* (see Figure 4e) and *TGW6* (see Figure 4f), the loss of a small exonic region in the TN1 gene makes it different from TNG67. These differences are seen as the gaps in the protein sequence alignment. For *GW7*/*GL7*, Nipponbare has a different gene structure compared to TNG67 and TN1 (Figure 4g),

**Figure 4.**
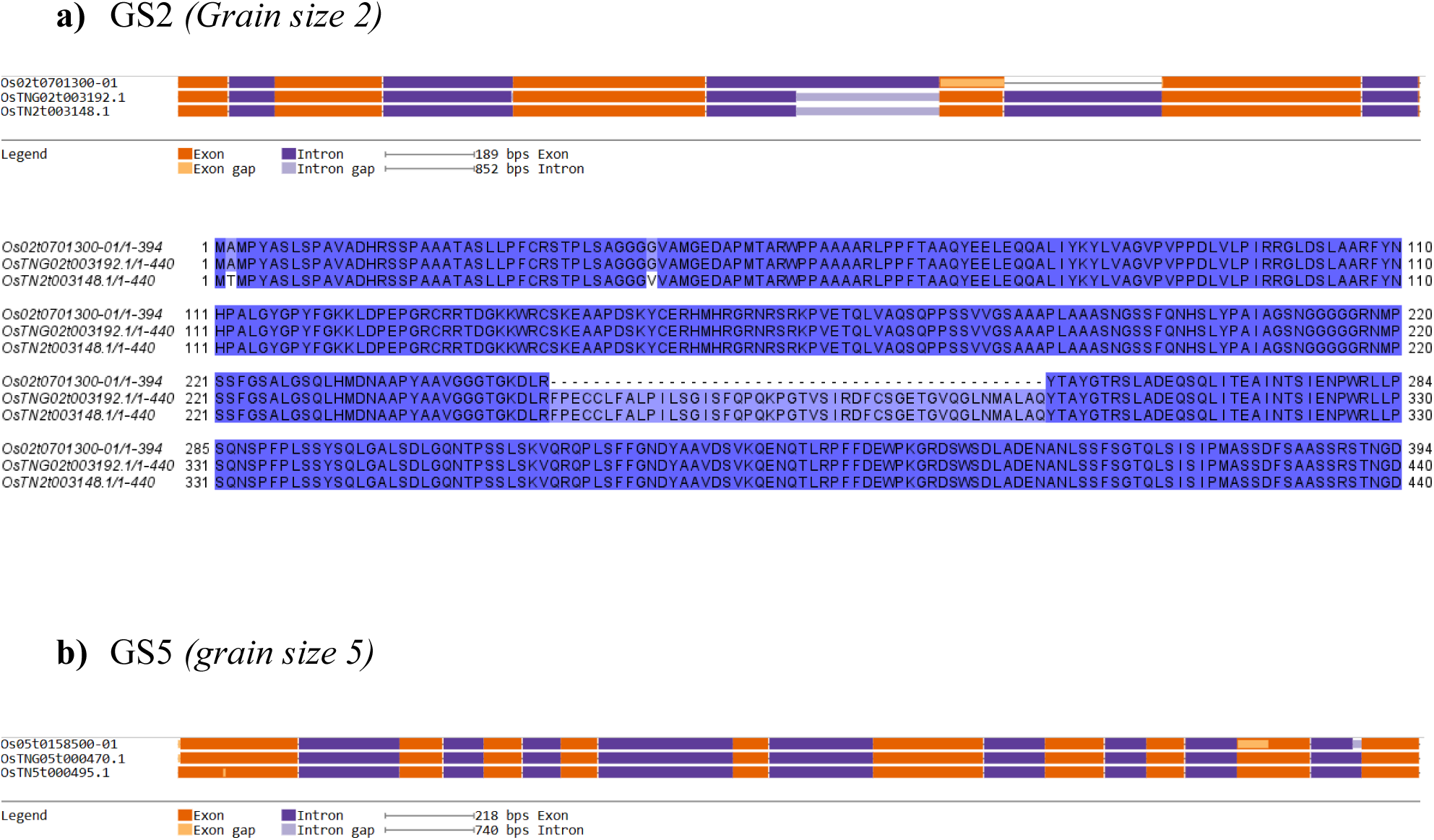
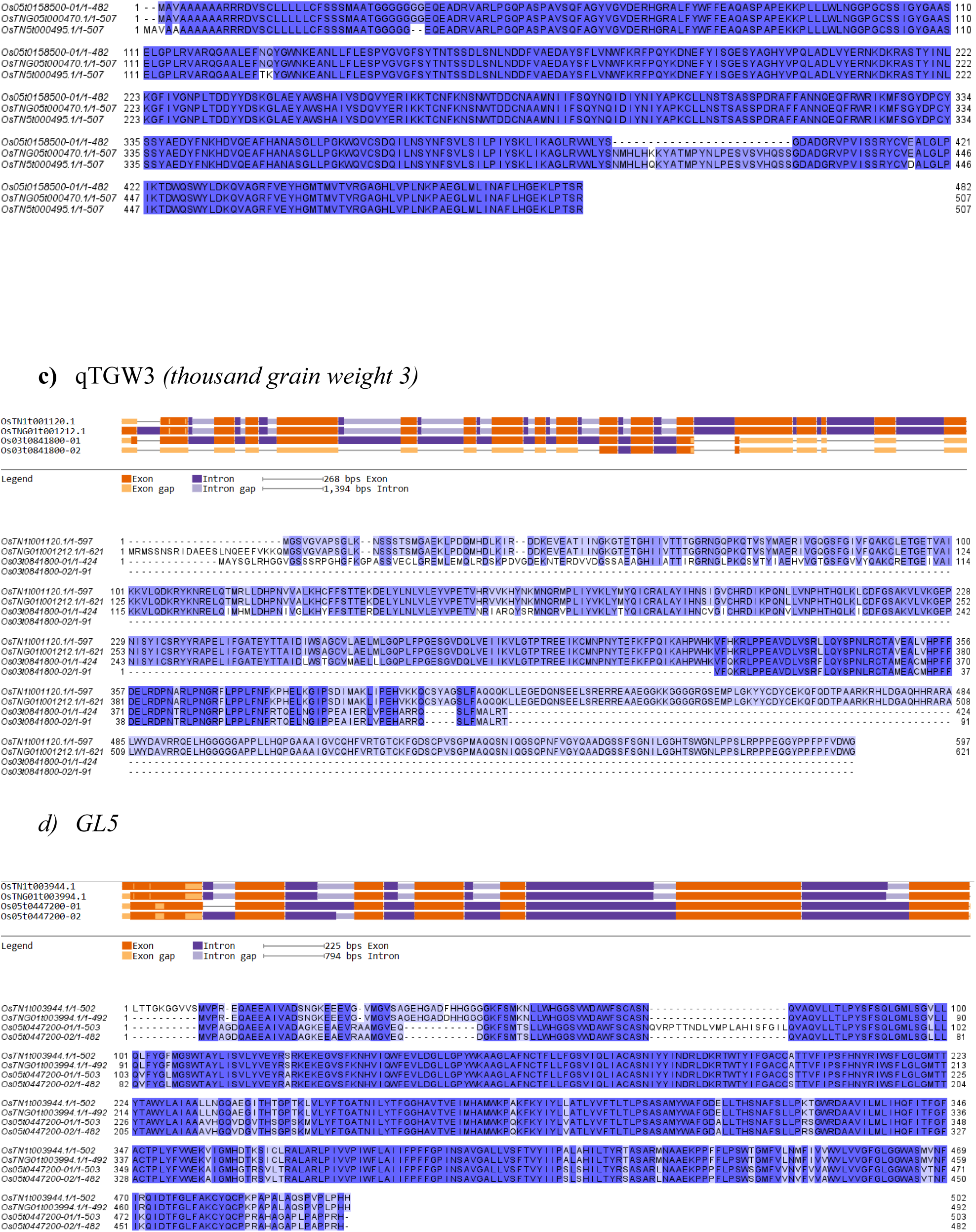
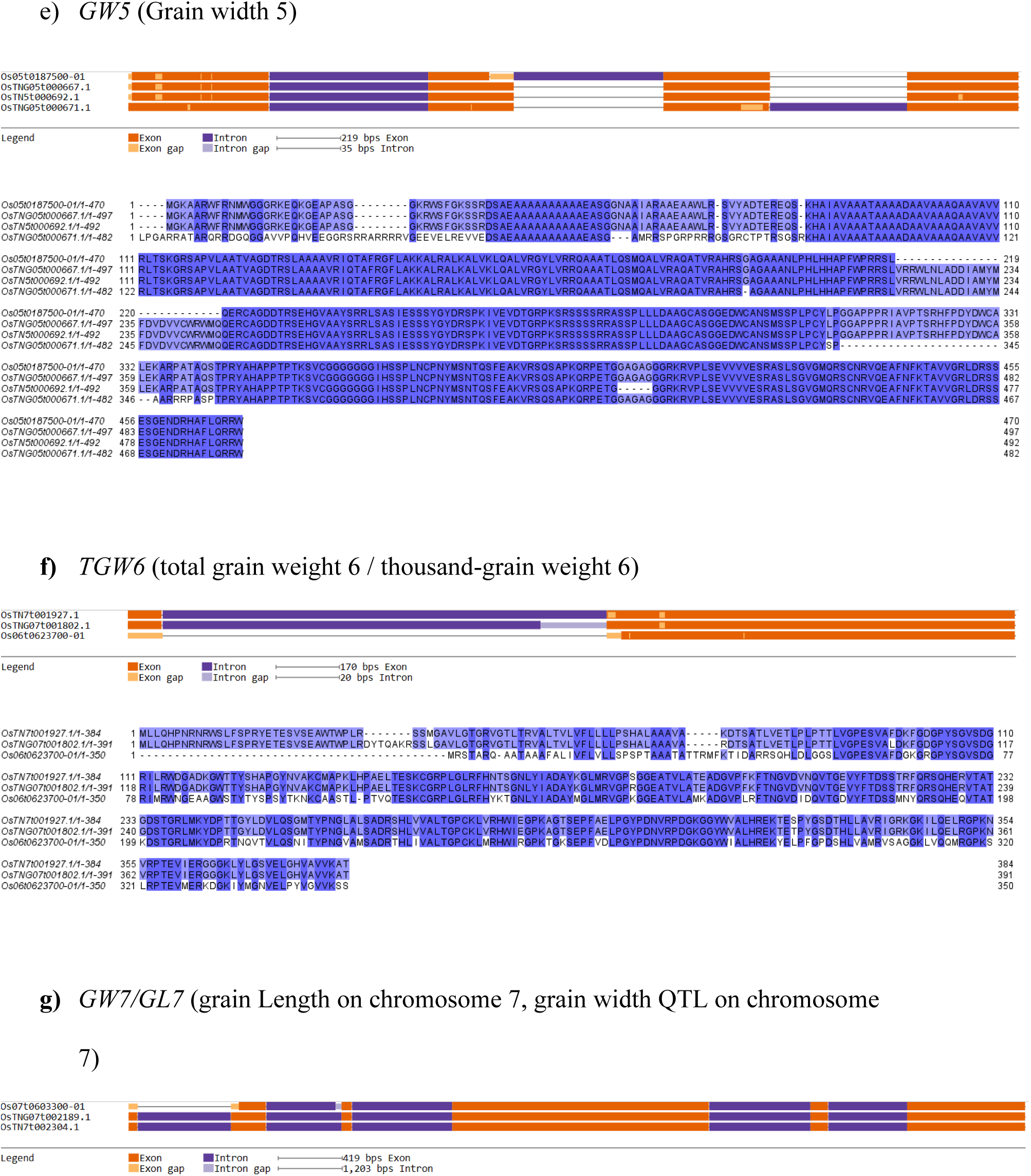
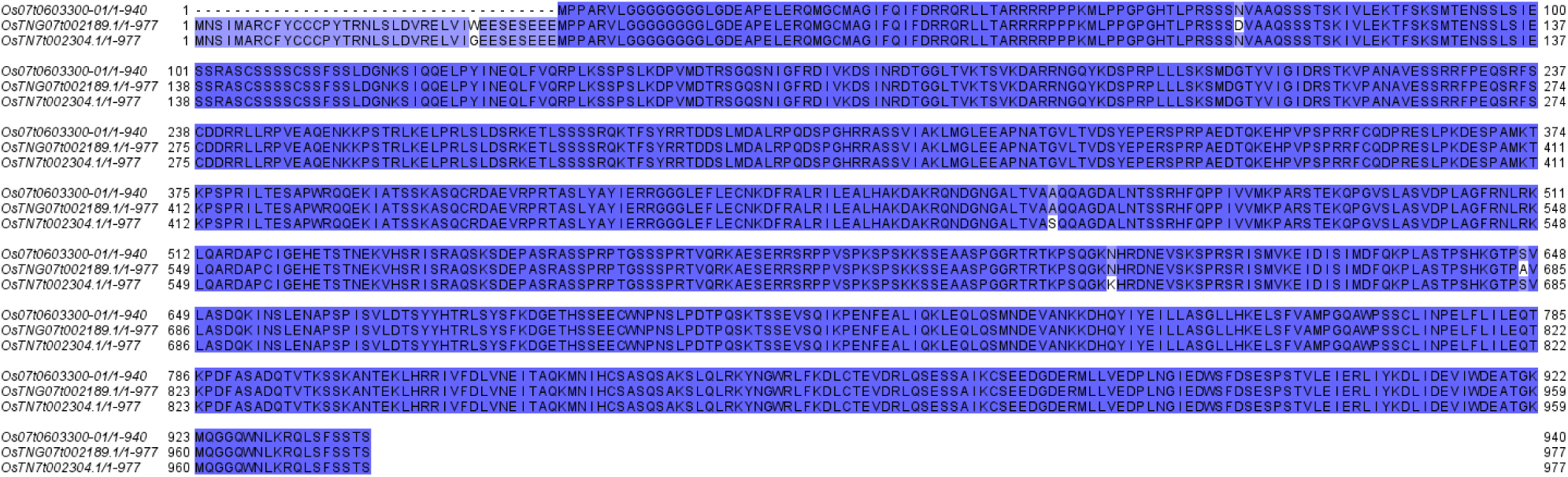
Gene structure diagrams of grain size genes wherein TNG67 and TN1 are more similar to each other than to Nipponbare. a) *GS2*: TNG67 and TN1 have similar gene structures. b) *GS5*: TN1 and TNG67 have similar gene structures. c) *qTGW3*: Except for the second isoform of Nipponbare, TN1 and TNG67 are similar to each other. d) *GL5*: TNG67 and TN1 are similar in the N-terminal part, which is different in Nipponbare. The remaining exonic regions are similar among the three cultivars. e) *GW5*: TNG67 and TN1 are similar, and the loss of the exonic region in the second exon of the Nipponbare GW5 makes it different from the same gene in TNG67 and TN1. f) *TGW6*: TN1 and TNG67 have some similarities, like the presence of exon 1 and the loss of an exonic region in the second exon upper Part. g) *GW7/GL7*: TNG67 and TN1 are the same

Our analysis of the TNG67-TN1 similar genes versus the TNG67-Nipponbare similar genes shows that the latter are more similar to each other in terms of their gene structure and protein sequence alignment. These comparisons reveal small exonic losses or gains, as well as amino acid changes that could not be seen on the gene structure diagrams. The amino acid changes may alter the protein structure, thereby making the protein product of the gene different between TNG67 and TN1. The results suggest that those TN1 grain size genes identified to be similar to their TNG67 counterparts are actually not really similar (in protein sequence). We conclude that the very high similarity of *GW6a*, GW6, and *WTG1* in terms of their gene structure and protein sequence changes between TNG67 and Nipponbare suggests that these three genes in TNG67 function like typical Nipponbare genes and may explain TNG67’s short and round grain shape, as in *japonica* cultivars.

### Analysis of photoperiod genes of TN67, Nipponbare and TN1

We compared the photoperiod genes of TNG67, Nipponbare, and TN1. From the list of common photoperiod genes by Mondal et al. (2019), we listed the gene names, the gene IDs, and each Nipponbare photoperiod protein and its TNG67 and TN1 orthologues (see Table 4). Then, we used GenePainter and ClustalW to create the gene models (diagrams) and protein sequence alignments. From the gene diagrams, patterns would appear whether a TNG67 gene is similar to both Nipponbare and TN1 (see Figure S6), is similar to Nipponbare (see Figure S7), to TN1, unique (see Figure S8), or hybrid (see Figure S9). TNG67 photoperiod genes would be grouped based on this criterion, as we did for the grain size genes.

**Table 4.**
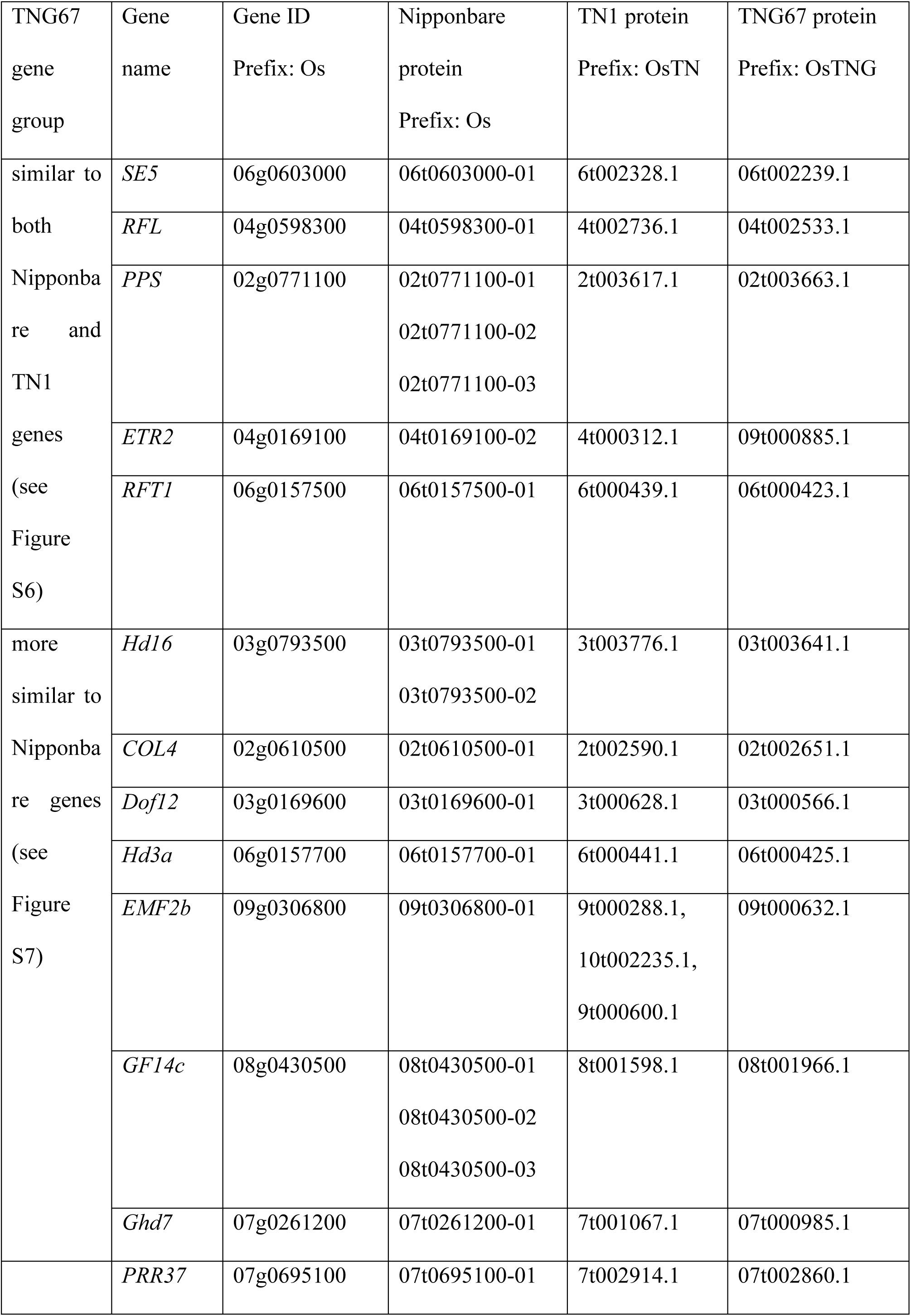
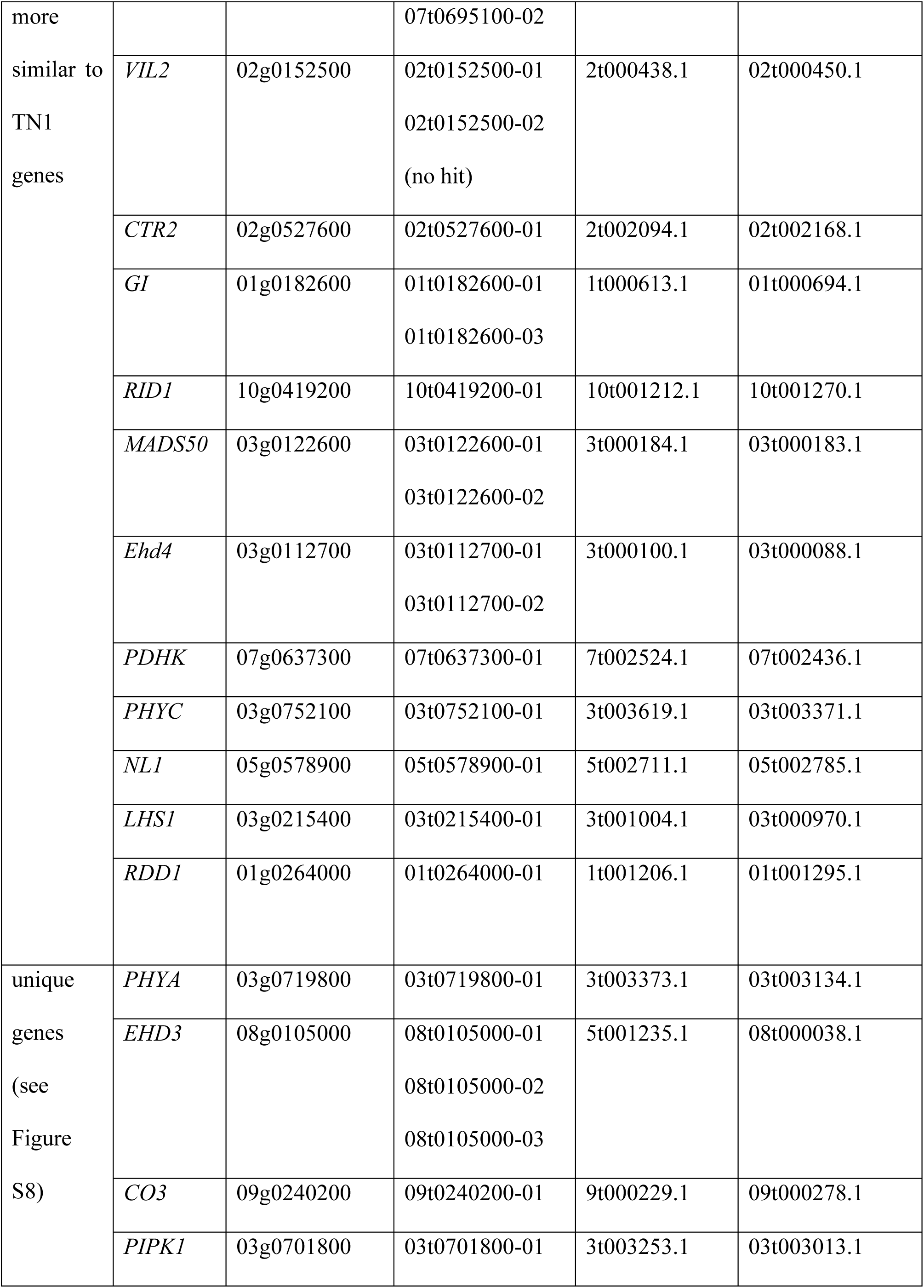
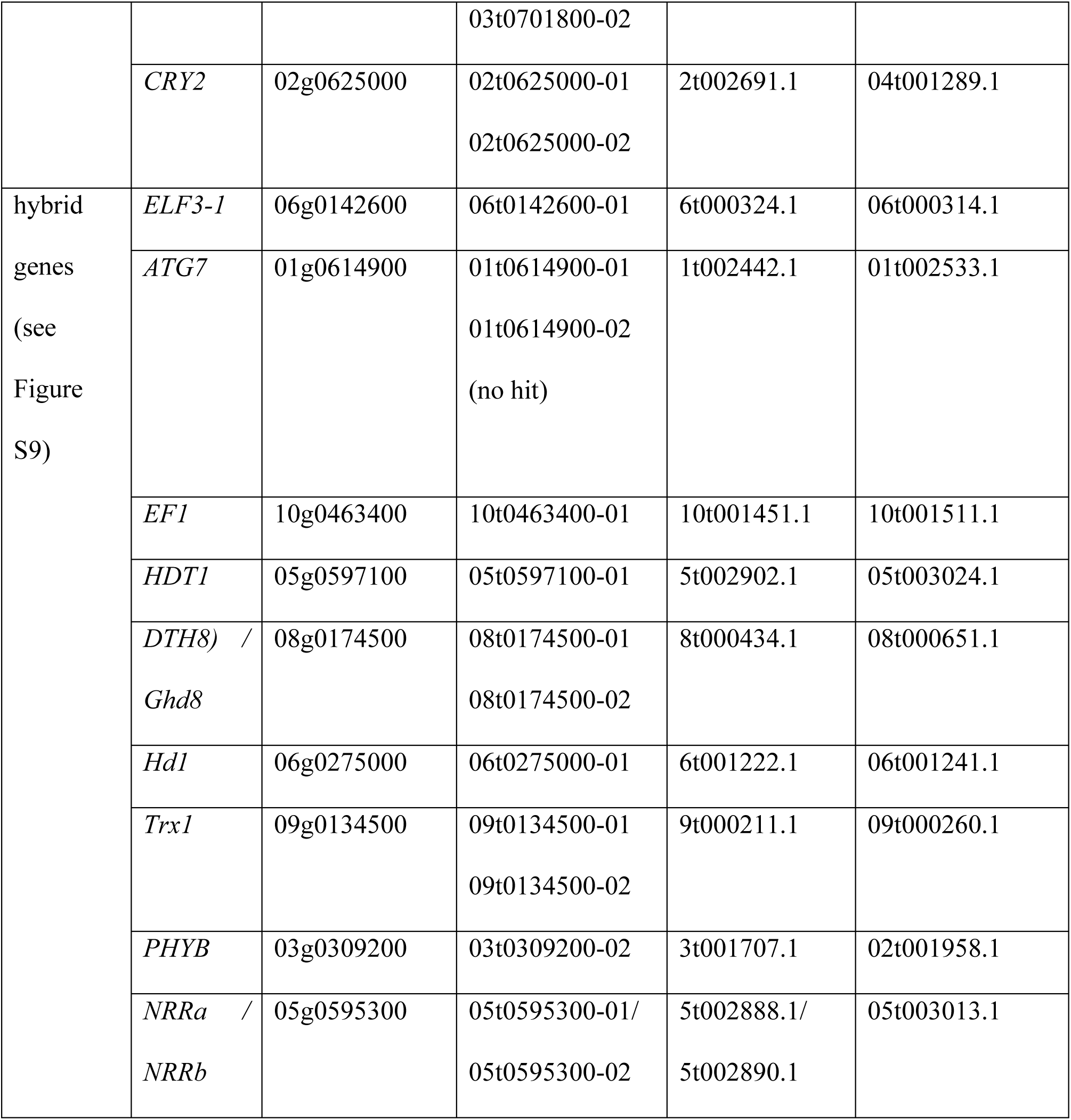
Photoperiod genes of TNG67, Nipponbare and TN1. The gene IDs were searched in the RAP-DB website using each gene name as the query. Gene names: *SE5* (Photosensitivity 5); *RFL* (Rice LFY-like gene, Probable transcription factor FL); PPS (peter pan syndrome); *ETR2* (ethylene response 2); *RFT1* (Rice Flowering-locus T 1); *Hd16* (heading date 16); *COL4* (CONSTANS-like gene 4); *Hd3a* (heading date 3A); *EMF2b* (embryonic flower 2b); *PRR37* (pseudo-response regulator 37); *CTR2* (constitutive triple-response2); *GI* (gigantea); *RID1* (Rice Indeterminate 1); *MADS50* (MADS-box transcription factor 50); Ehd4 (Early heading date 4); *PDHK* (pyruvate dehydrogenase kinase); *PHYC* (Phytochrome C); *NL1* (neck leaf 1); *LHS1* (leafy hull sterile 1); *RDD1* (rice dof daily fluctuations 1); *PHYA* (Phytochrome A); *EHD3* (Early heading date 3); *CO3* (CONSTANS 3); *PIPK1* (rice phosphatidylinositol monophosphate kinase 1); *CRY2* (cryptochrome 2); *ELF3-1* (early flowering 3-1); *ATG7* (autophagy-related 7); *EF1* (Earliness 1); *HDT1* (histone deacetylase 1); *DTH8* (days to heading on chromosome 8); *Ghd8* (Grain number, plant height and heading date 8); *Hd1* (Heading date 1); *Trx1* (*Oryza sativa* Trithorax1); *PHYB* (Phytochrome B); *NRRa* (nutrition response and root growth a); *NRRb* (nutrition response and root growth b).

Table 4 shows that there are only five TNG67 photoperiod genes that are more similar to Nipponbare genes than to TN1 genes whereas there are 12 TNG67 photoperiod genes that are more similar to TN1 genes than to Nipponbare genes. These 12 genes are *PRR37* (pseudo-response regulator 37), *VIL2* (vernalization insensitive 3-like 2), *CTR2* (constitutive triple-response 2), *GI* (gigantea), *RID1* (Rice Indeterminate 1),*MADS50* (MADS-box transcription factor 50), and *Ehd4* (early heading date 4), *PDHK* (pyruvate dehydrogenase kinase), *PHYC* (phytochrome C), *NL1* (neck leaf 1), *LHS1* (leafy hull sterile 1) and *RDD1* (rice dof daily fluctuations 1). These genes are discussed below. Note that the prefixes OsTN1, OsTNG and Os of gene IDs stand for TN1, TNG67 and Nipponbare, respectively.

For *PRR37*, the two isoforms of Nipponbare (TN7t002914.1 and OsTNG07t002860.1) are easily spotted because they share the loss of two exonic regions, both of which are present in the TN1 and TNG67 genes (see Figure 5a). The protein sequence alignment also clearly shows that TNG67 PRR37 is more similar to TN1 PRR37 than to Nipponbare PRR37 (see Figure 5a). *VIL2* is a perfect example where the gene structure models of TNG67 and TN1 are like mirror images of each other (see Figure 5b). Thus, TNG67 *VIL2* is more similar to TN1 *VIL2* than to Nipponbare *VIL2*. The same can be said about *CTR2* (see Figure 5c). For *GI*, TNG67’s OsTNG01t000694.1 and TN1’s OsTN1t000613.1 share the loss of an exonic region near the 3’ end and are more similar to each other than either of them is to Nipponbare’s two isoforms, which share the second to the last exon near the 3’ end (see Figure 5d).

**Figure 5.**
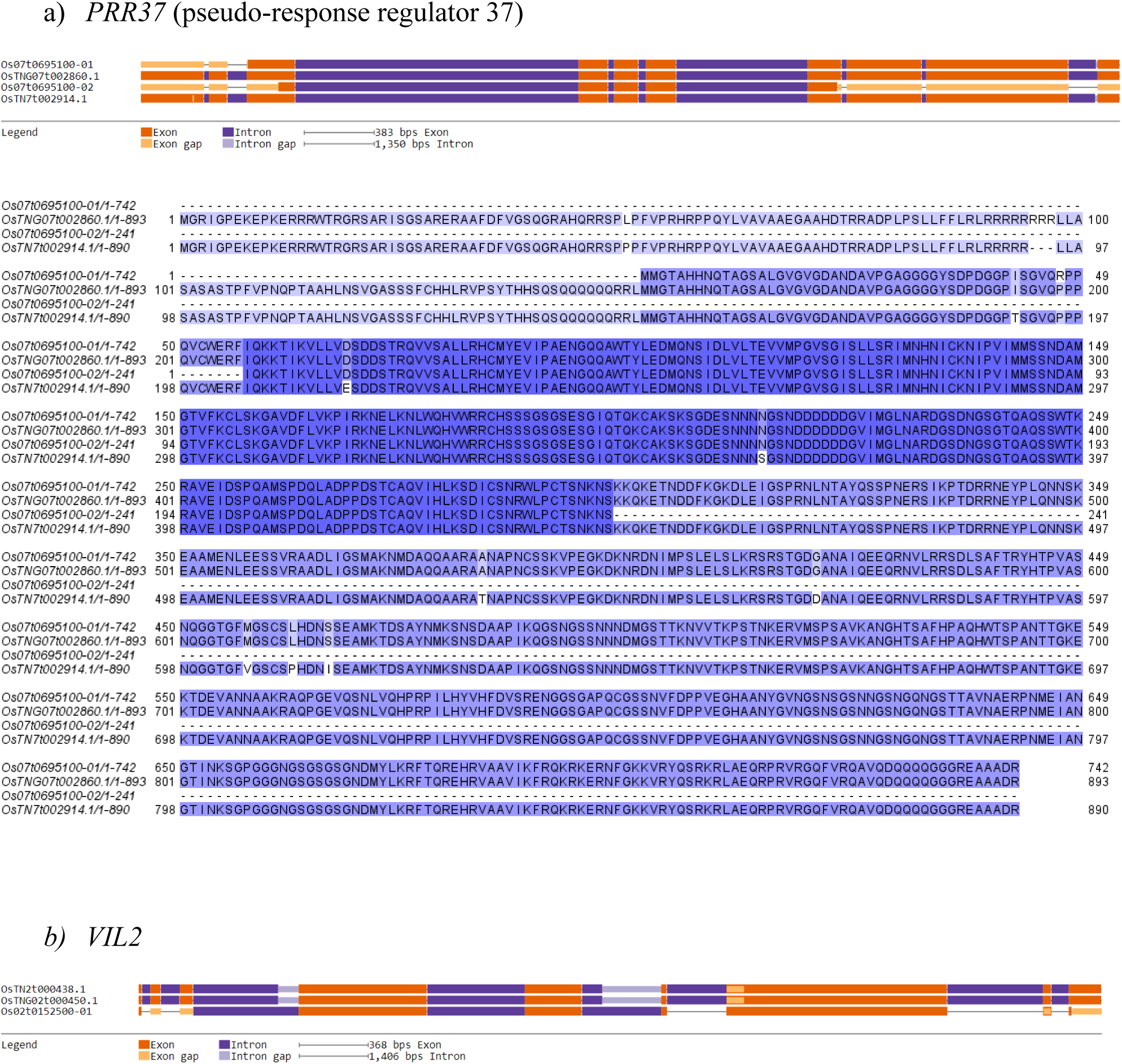
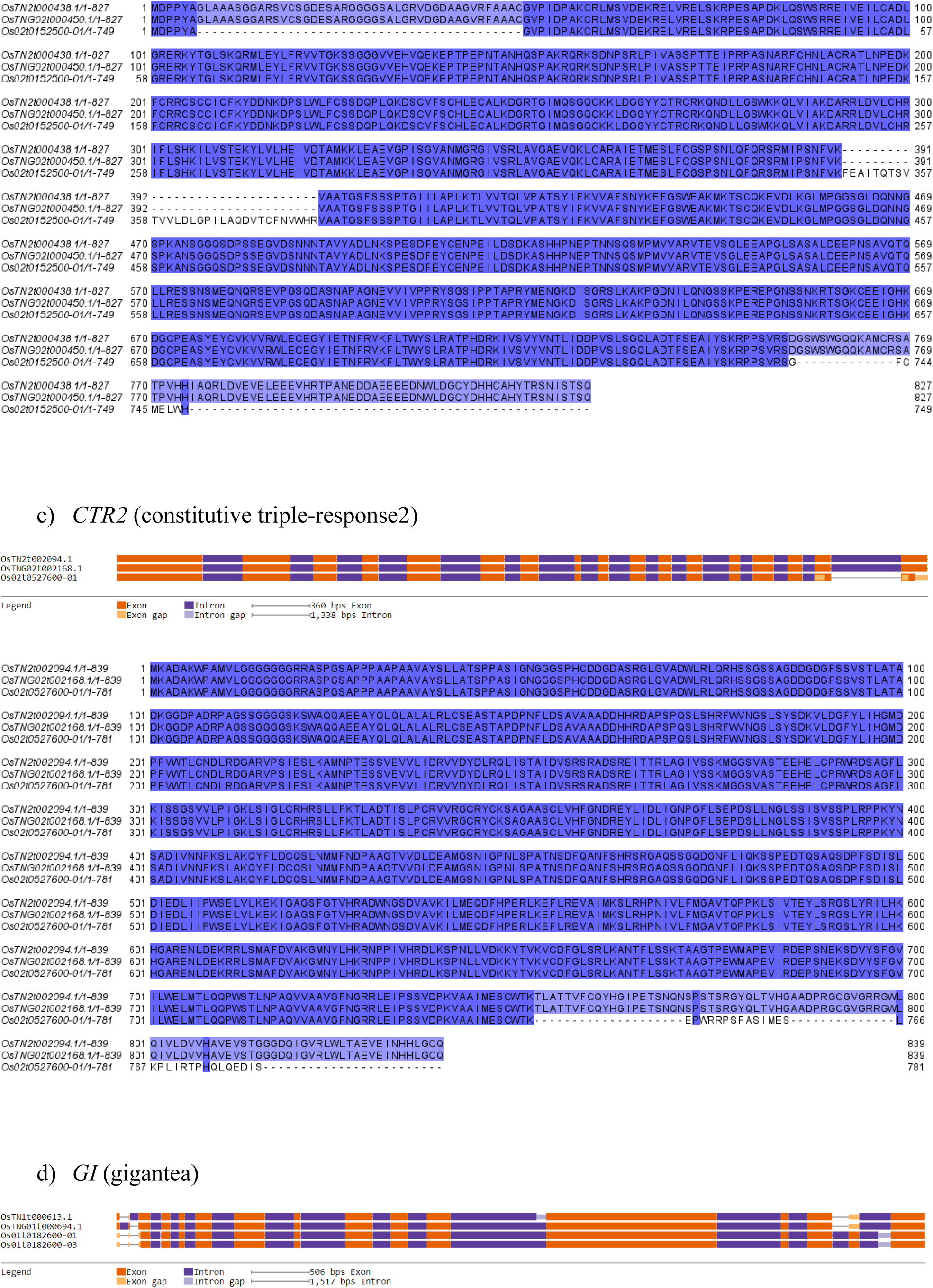
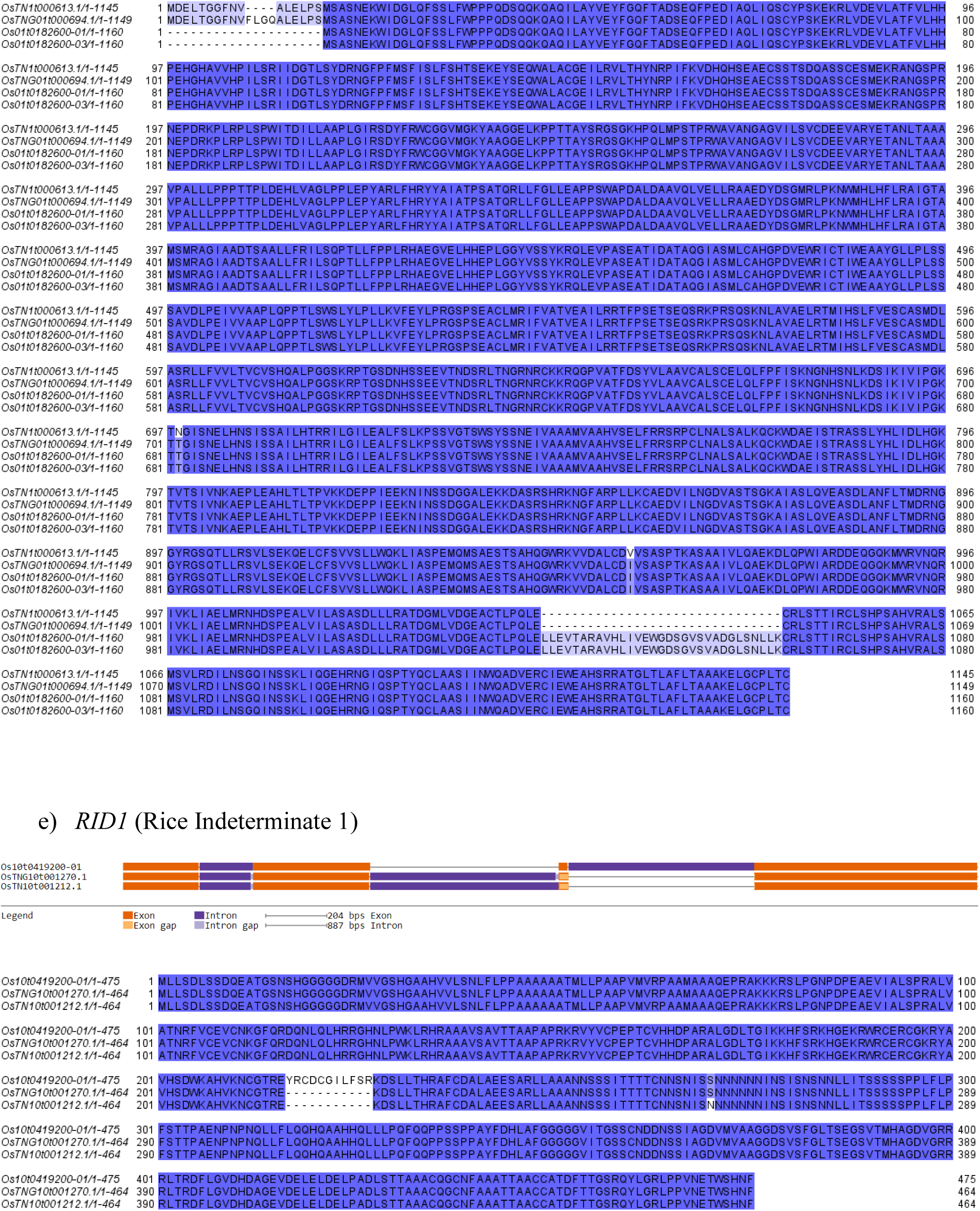
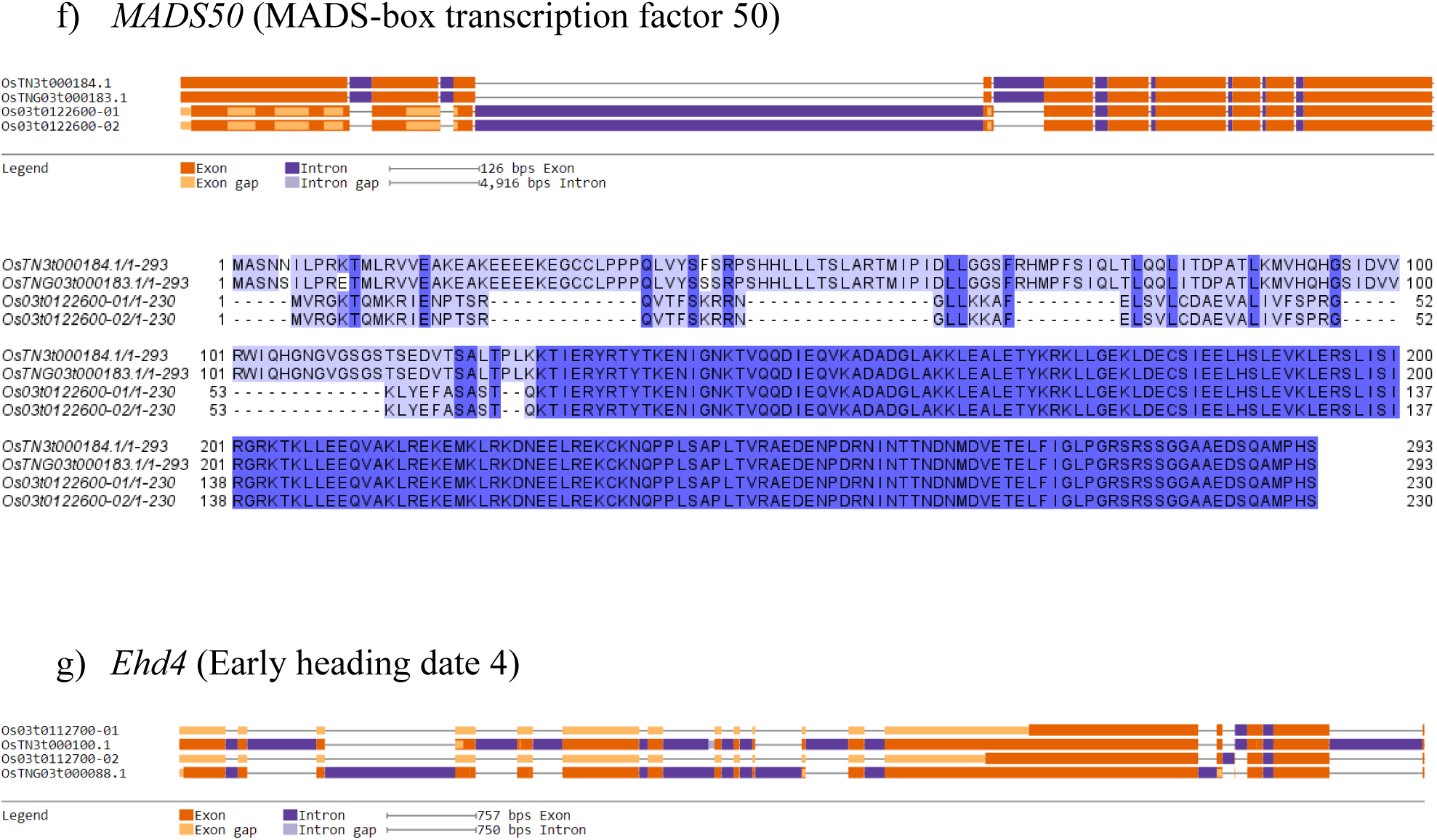
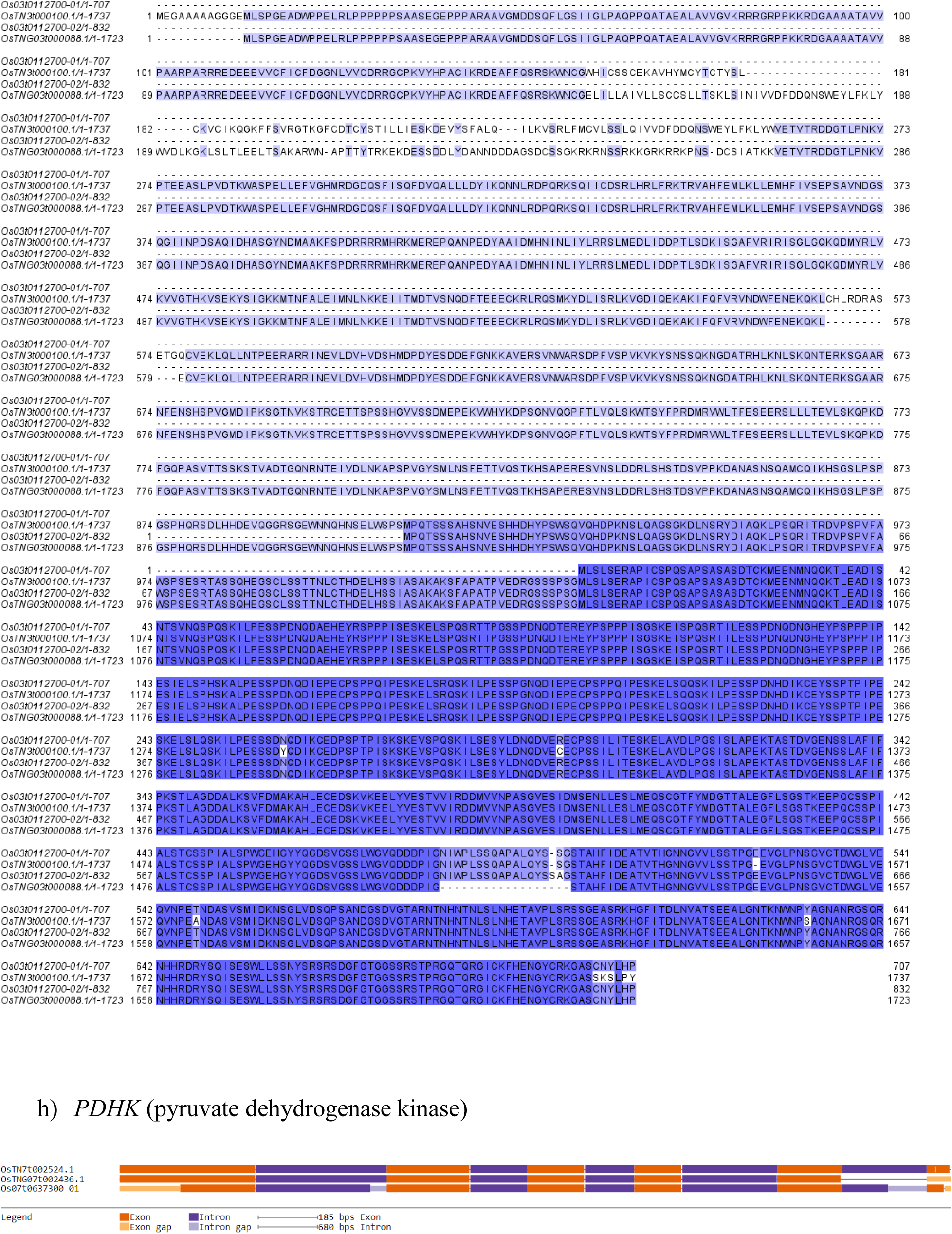
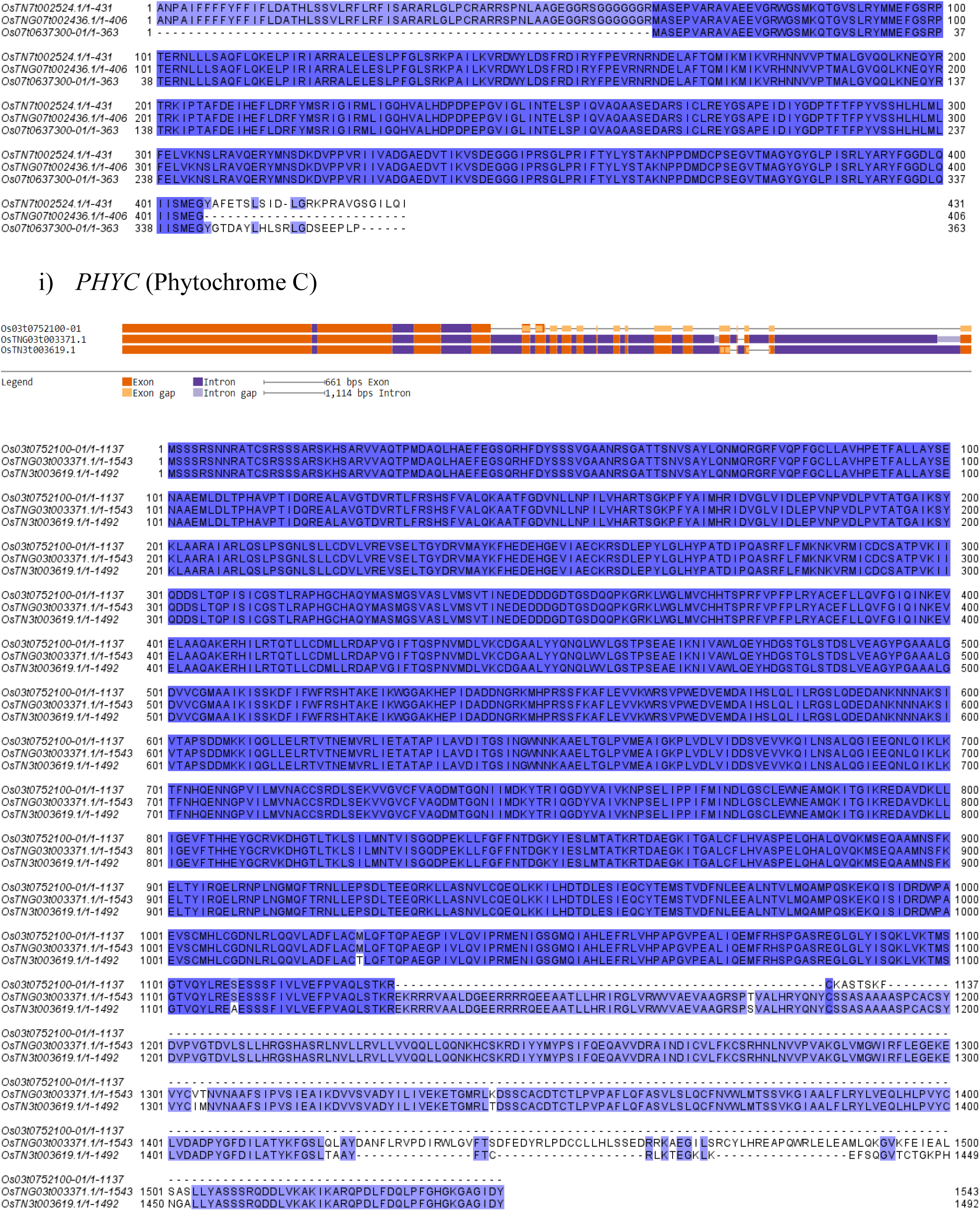
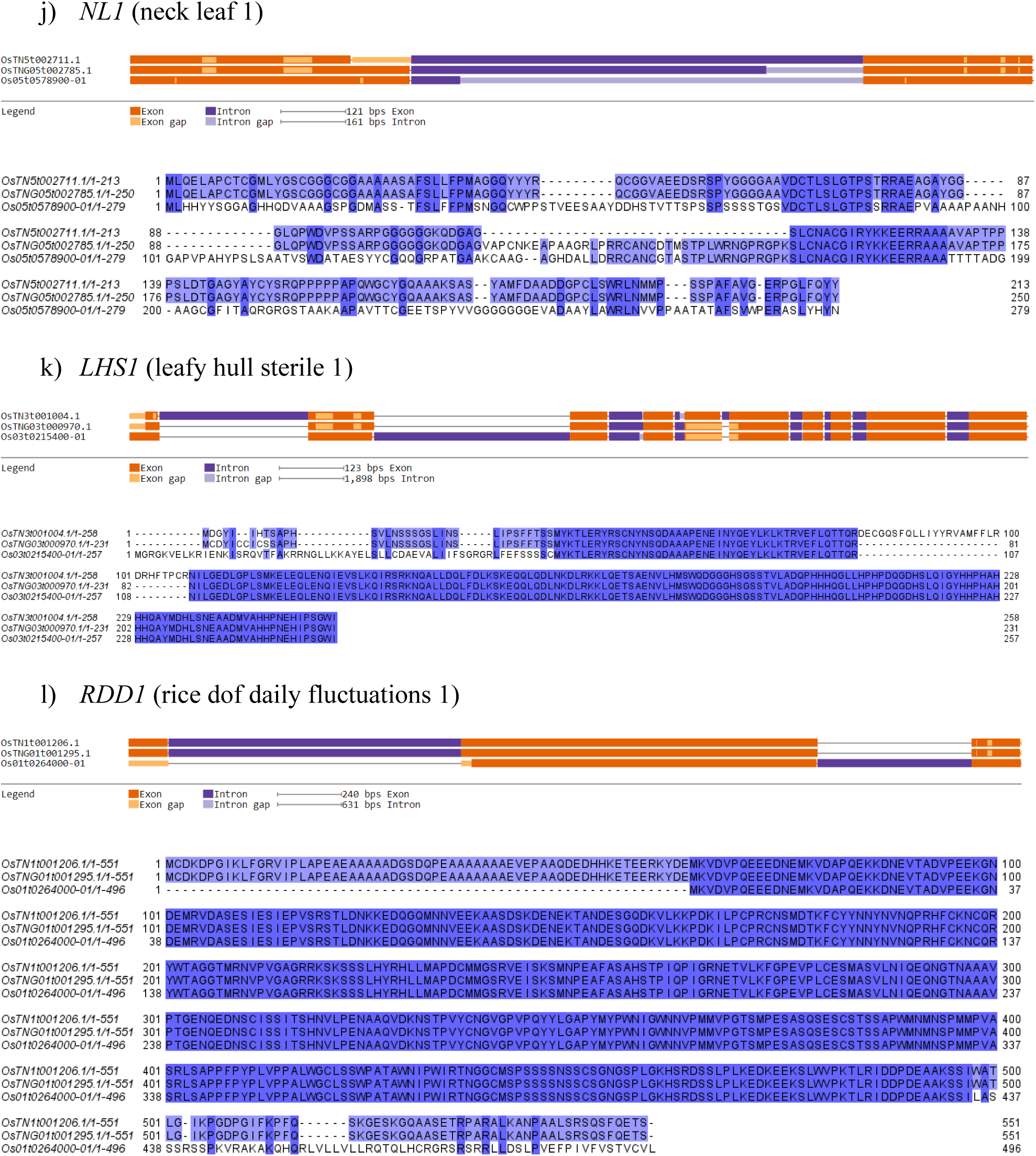
Photoperiod genes of TNG67 that are more similar to their TN1 orthologues than Nipponbare orthologues. a) *PRR37*, b) *VIL2*, c) *CTR2*, d) *OsGI*, e) *RID1*, f) *MADS50*, g) *Ehd4*, h) *PDHK*, i) *PHYC*, j) *NL1*, k) *LHS1*, and l) *RDD1*

For *RID1*, TNG67’s OsTNG10t001270.1 and TN1’s OsTN10t001212.1 share the loss of 11 codons in an exonic region in their gene structure (*see* Figure 5e). At the protein sequence level, TNG67 RID1 and TN1 RID1 differ by only one amino acid, i.e., serine vs. asparagine, both of which are polar and uncharged. Therefore, the TNG67 and TN1 RID1 proteins are very similar. For *MADS50* (Figure 5f), it is clear from the gene structure models and sequence alignment that TNG67’s OsTNG03t000183.1 and TN1’s OsTN3t000184.1 are more similar to each other than either of them is to Nipponbare’s Os03t0122600-01 and Os03t0122600-02, which show losses of exonic regions when compared to their orthologues in TNG67 and TN1 (see Figure 5f). The same can be said to *Ehd4* (Figure 5g), where the patterns of missing exons agree better between TNG67’s OsTNG03t000088.1 and TN1’s OsTN3t000100.1 than between either of them and Nipponbare’s Os03t0112700-01 or Os03t0112700-02, which are isoforms.

For *PDHK* (see Figure 5h), TNG67’s OsTNG07t002436.1 and TN1’s OsTN7t002524.1 have a better sequence alignment than the alignment between either of them and Nipponbare’s Os07t0637300-01. The first amino acid of Nipponbare’s PDHK is methionine, while it is alanine for both TNG67 and TN1 PDHK, although alanine might be an annotation error (see Figure 5h). For *PHYC* (Figure 5i), the gene structure diagrams show many deletions in Nipponbare’s gene, which lead to the losses of long amino acid segments in Nipponbare PHYC. Thus, TNG67 PHYC is more similar to TN1 PHYC than to Nipponbare PHYC (Figure 5i). For NL1, the protein sequence alignment shows that TNG67 and TN1 NL1 share many deletions when compared to Nipponbare NL1 (Figure 5j). Thus, TNG67 NL1 is more similar to TN1 NL1 than to Nipponbare NL1. For LHS1, there are similarities in the patterns of exonic losses between TNG67’s OsTNG03t000970.1 and TN1’s OsTN3t001004.1, and between TNG67’s OsTNG03t000970.1 and Nipponbare’s Os03t0215400-01. However, these lost exonic regions do not contain any coding sequences, and the protein sequence alignment (see Figure 5k) reveals that there are more matching sequence segments between TNG67 and TN1. Thus, TNG67 LHS1 is more similar to TN1 LHS1 than to Nipponbare LHS1. For *RDD1*, the gene structure models clearly show higher similarities between TNG67 and TN1 (see Figure 5l). For the protein sequence alignment, the violet-blue colored sequences correspond to matches between TNG67 RDD1 and TN1 RDD1 (Figure 5l), confirming that TNG67 RDD1 and TN1 RDD1 are more similar to each other than either of them is to Nipponbare RDD1.

### Determining the origin of TNG67 genomic sequences and genes

We tried to determine the origin of each annotated TNG67 gene, as to whether it was derived from its *japonica* direct and indirect parents or from TN1, which is an *indica* cultivar. We conducted a progressive alignment between the chromosomes of TNG67, Nipponbare and TN1 to get a backbone file for each set of chromosomes compared. The backbone file contained the coordinates of genomic segments that were aligned among the three genomes (see Table 6), between two genomes only (see Table 7), or non-backbone (non-aligned) segments (see Table 8). We converted the coordinates to a bed file format, and extracted the genes using bedtools. Of particular interest are backbone sequences shared by two genomes only (see Table 7), specifically TNG67 vs Nipponbare or the TNG67 vs TN1, which would indicate that the TNG67 sequences were derived from Nipponbare or from TN1. The illustration of the backbone is in Figure 6.

**Figure 6.**
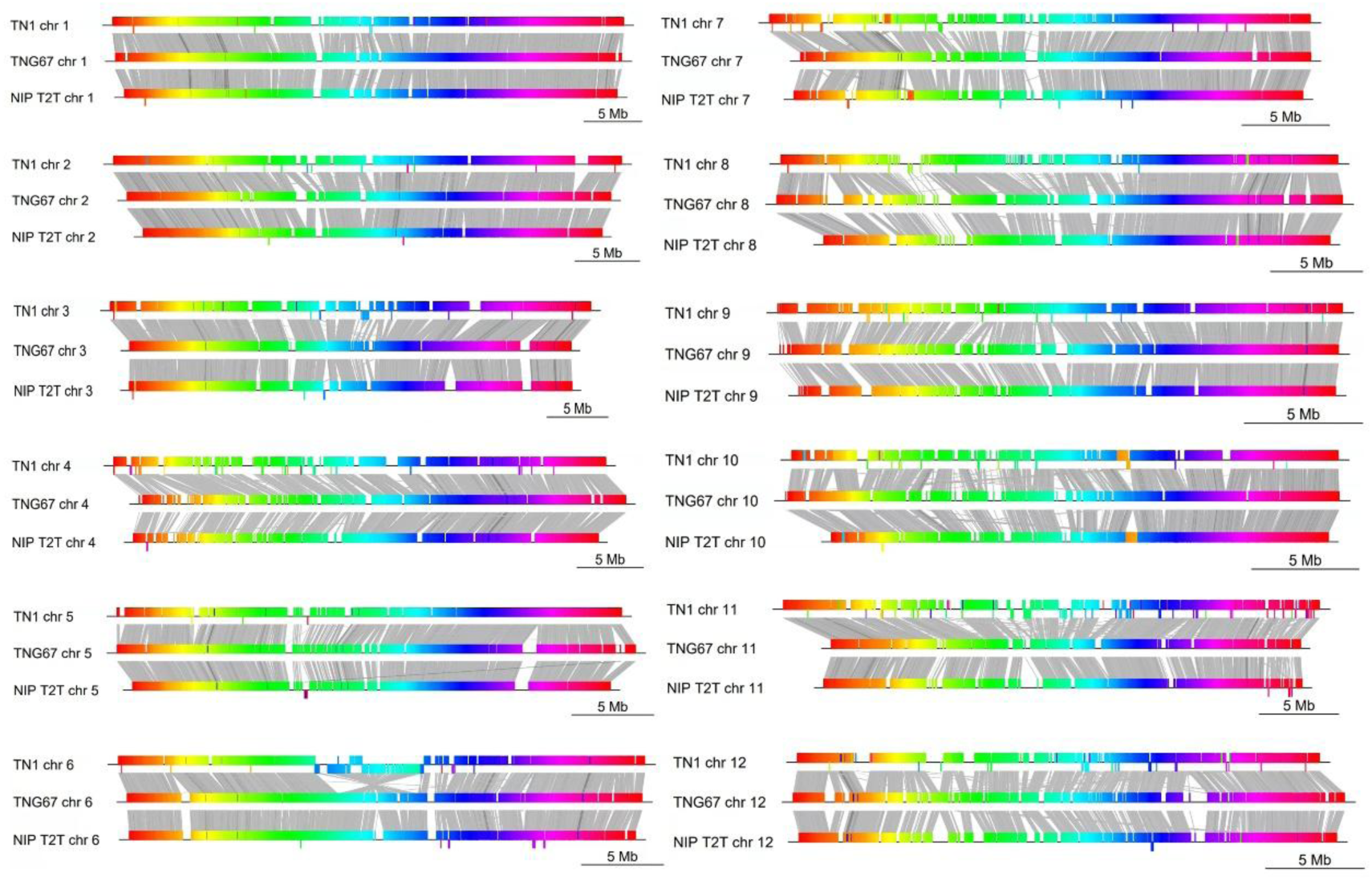
Progressive Mauve alignment of the chromosomes of TNG67 vs. TN1 and TNG67 vs. Nipponbare T2T. In a left to right direction, the chromosomes are in forward strand. Regions that are homologous have the same color and are connected by gray lines. Segments that have high conservation are represented as darker regions in the gray lines, and also dark regions in the colored blocks. Because two chromosomes are being compared at a time, the chromosome at the top or below are the references (TN1 or Nipponbare T2T). The arrangement of the colored blocks of the reference chromosome is fixed, and only the chromosome of TNG67 can indicate the rearrangements. An inverted region is seen as a protruding bar below, and a large inversion can be clearly seen on the TNG67-TN1 alignment for chromosome 6. In this figure, more inversions can be seen on TNG67-TN1 than on TNG67-Nipponbare. More conserved regions are also observed when TNG67 and Nipponbare are compared.

**Table 6.** Total length of backbone sequences in a genome that are shared among the three cultivars. The length is in terms of base pairs (bp).

|  | NIP-T2T | TNG67 | TN1 |
| --- | --- | --- | --- |
| chr 1 | 31,492,082 | 31,452,017 | 31,489,281 |
| chr 2 | 27,249,987 | 27,215,663 | 27,249,147 |
| chr 3 | 27,128,141 | 27,094,818 | 27,126,181 |
| chr 4 | 20,123,159 | 20,095,273 | 20,131,502 |
| chr 5 | 20,658,794 | 20,487,530 | 20,522,685 |
| chr 6 | 20,958,465 | 20,937,131 | 20,964,518 |
| chr 7 | 18,470,815 | 18,461,453 | 18,503,709 |
| chr 8 | 17,440,333 | 17,442,726 | 17,439,747 |
| chr 9 | 14,574,917 | 14,557,384 | 14,573,165 |
| chr 10 | 14,753,412 | 14,735,203 | 14,750,364 |
| chr 11 | 16,219,096 | 16,189,109 | 16,233,342 |
| chr 12 | 14,544,564 | 14,520,654 | 14,545,458 |
| Sum | 243,613,765 | 243,188,961 | 243,529,099 |

**Table 7.** Total length of backbone sequences that are shared between two genomes. The genomes of the three cultivars were first aligned by progressive Mauve. The resulting backbone file was inspected to determine the length of the shared sequences for each pairwise comparison. For each pair of genomes compared, the third genome that was not aligned was assumed to have not been aligned to the other two genomes. For example: under TNG67 vs NIP-T2T, the segments that were shared by TNG67 and IRGSP, but not by TN1, were counted. The length is in terms of base pairs (bp).

| Comparison | TNG67 vs NIP-T2T |  | TNG67 vs TN1 |  | NIP T2T vs TN1 |  |
| --- | --- | --- | --- | --- | --- | --- |
| Cultivar | TNG67 | NIP T2T | TNG67 | TN1 | NIP T2T | TN1 |
| chr 1 | 5,741,394 | 5,823,852 | 373,611 | 378,683 | 1,204,898 | 1,212,248 |
| chr 2 | 4,904,154 | 4,969,089 | 164,343 | 165,947 | 724,295 | 724,923 |
| chr 3 | 4,342,731 | 4,400,306 | 190,906 | 193,467 | 1,420,941 | 1,421,557 |
| chr 4 | 5,310,202 | 5,380,336 | 380,939 | 389,109 | 1,295,619 | 1,296,450 |
| chr 5 | 4,422,711 | 4,480,749 | 174,398 | 176,490 | 324,236 | 325,267 |
| chr 6 | 4,464,662 | 4,512,070 | 393,853 | 460,791 | 626,960 | 644,734 |
| chr 7 | 4,013,440 | 4,061,734 | 374,591 | 380,109 | 1,653,797 | 1,658,429 |
| chr 8 | 5,505,515 | 5,562,211 | 257,821 | 263,296 | 465,400 | 465,685 |
| chr 9 | 3,737,130 | 3,777,468 | 155,438 | 156,591 | 1,241,489 | 1,241,807 |
| chr 10 | 3,905,200 | 3,951,045 | 348,841 | 452,418 | 816,050 | 816,290 |
| chr 11 | 6,021,096 | 6,090,072 | 360,071 | 389,846 | 1,416,279 | 1,449,756 |
| chr 12 | 5,542,009 | 5,584,591 | 309,307 | 319,715 | 1,779,670 | 1,760,466 |
| Sum | 57,910,244 | 58,593,523 | 3,484,119 | 3,726,462 | 12,969,634 | 13,017,612 |

**Table 8.** Total length of non-backbone sequences in each cultivar. The counts refer to the length of sequences in each genome that did not align to the two other genomes. The length is in terms of base pairs (bp).

|  | NIP-T2T | TNG67 | TN1 |
| --- | --- | --- | --- |
| chr 1 | 5,397,347 | 7,549,389 | 12,421,625 |
| chr 2 | 3,471,909 | 6,101,272 | 12,165,809 |
| chr 3 | 4,427,541 | 5,642,688 | 11,847,464 |
| chr 4 | 9,268,820 | 11,962,339 | 16,345,515 |
| chr 5 | 4,629,559 | 7,599,258 | 10,677,422 |
| chr 6 | 5,807,942 | 6,457,167 | 10,984,765 |
| chr 7 | 5,739,400 | 7,159,562 | 11,215,453 |
| chr 8 | 5,048,918 | 8,615,116 | 12,925,217 |
| chr 9 | 7,779,369 | 5,843,827 | 8,581,317 |
| chr 10 | 4,361,434 | 7,591,840 | 10,349,051 |
| chr 11 | 7,337,814 | 7,986,111 | 16,693,069 |
| chr 12 | 6,335,316 | 8,454,584 | 10,867,802 |
| sum | 69,605,369 | 90,963,153 | 145,074,509 |

In Table 7, the numbers for TNG67 vs TN1 are smaller, but in terms of the non-backbone sequences (see Table 8) for TN1 is the biggest at 145 Mb, which means it has the most segments that are unique and will not align to either TNG67 or NIP T2T. Because 145 Mb is unique only to TN1, few will be left to align to TN1 and NIP-T2T. In terms of the backbone sequences common to all the three genomes (see Table 6), they are the same at 243 Mb. So the numbers for the TNG67 vs TN1 which is 3.5 Mb and 3.7 Mb, would refer to segments of TNG67 derived from TN1.

Using the information from the backbone sequences shared between two genomes, we extracted the genes that have overlaps with these backbone sequences. We classified these genes into (1) TNG67 genes derived from *japonica* ancestry, (2) TNG67 genes derived from *indica* ancestry, (3) TNG67 genes derived from *japonica* ancestry and not found in TN1 and (4) TNG67 genes derived from *indica* ancestry and not found in Nipponbare (Table 9).

**Table 9.** Numbers of TNG67 genes derived from *japonica* ancestry (JAP) and numbers of TNG67 genes derived from *indica* ancestry (IND)

|  | TNG67<br>genes from<br>*japonica*<br>ancestry | TNG67 genes<br>from *indica*<br>ancestry | TNG67 genes from<br>*japonica* ancestry<br>and not found in<br>TN1 | TNG67 genes derived<br>from *indica* ancestry<br>and not found in<br>Nipponbare |
| --- | --- | --- | --- | --- |
| chr 1 | 1332 | 169 | 180 | 15 |
| chr 2 | 1091 | 61 | 142 | 8 |
| chr 3 | 1077 | 75 | 174 | 10 |
| chr 4 | 890 | 83 | 189 | 33 |
| chr 5 | 874 | 49 | 209 | 13 |
| chr 6 | 853 | 137 | 169 | 17 |
| chr 7 | 823 | 71 | 105 | 24 |
| chr 8 | 921 | 47 | 275 | 14 |
| chr 9 | 696 | 29 | 81 | 5 |
| chr 10 | 735 | 79 | 118 | 21 |
| chr 11 | 867 | 129 | 262 | 21 |
| chr 12 | 734 | 76 | 199 | 15 |
| sum | 10,893 | 1,005 | 2,103 | 196 |

### funRiceGenes search

Using the funRiceGenes database (Huang et al., 2022) with keywords associated with each Nipponbare-derived genes of TNG67, we sought interesting features with terms related to blast resistance, grain size and yield, flowering time, and drought tolerance. Among those genes with blast resistance terms are OsTNG06g001327 (*Pi9|Piz-t|Pi2*), OsTNG08g000999 (*Pish*), OsTNG10g000462 (*acyl transferase 1*), OsTNG11g001701 (*Pish*), and OsTNG11g001702 (*Pish*). For TNG67 tagged with a blast keyword are OsTNG02g000113 (*γ-aminobutyrate transaminase*), OsTNG02g002216 and OsTNG02g002217 (*RAR1*), OsTNG03g003603 and OsTNG03g003604 (*calcium-dependent protein kinase 10*), OsTNG04g000993 (*terpene synthase 19*) and also genes that are associated with the blast resistance keywords, OsTNG06g001327, OsTNG10g000462, OsTNG08g000999, OsTNG11g001701, OsTNG11g001702. OsTNG03g003615 and OsTNG03g003616 are the only TNG67 genes that have an R protein keyword.

In our funRiceGenes analysis many genes have the keyword “resistance” (see Table S4). A recurring example is OsTNG06g001327, which codes for *Pi9*, *Piz-t*, or *Pi2* and is related to blast. *Pi9* is a paralog of *Piz-t* and *Pi2*, while are *Piz-t* and *Pi2* are alternative forms of the same gene. (Zhou et al., 2006) Another gene related to blast resistance is OsTNG04g000993 (*terpene synthase 19*). OsTNG07g000574 (*P-450 71Z2* | *Cytochrome P450 71Z2*) is associated with bacterial blight resistance. OsTNG05g001016 (*XIP*) is related to resistance against brown plant hopper. Others only have the “resistance” keyword in the funRiceGenes database. These genes are OsTNG01g001250 (*accelerator of internode elongation 1*), OsTNG04g001510 (*flavanone 3-hydroxylase* | *salicylic acid hydroxylase 1*), OsTNG11g001781 (*lipoxygenase 10*), and OsTNG11g002482 (*Phenylalanine ammonia-lyase 8*). An exception is OsTNG12g001689 (*pollen extensin 1*), which is not related to a pathogen but rather about lodging resistance.

Because TNG67 is a hybrid descendant of a *japonica* and *indica* cultivar, we sought for genes associated with flowering time, that are derived from Nipponbare, but not found in TN1. These genes are OsTNG02g001454 (*ubiquitin-like domain kinase γ4*), OsTNG03g003563 (*phytochrome-interacting factor-like protein 1*), and OsTNG11g001162 (*constans 3*), while the OsTNG05g001307 (*ZRT/IRT-like protein family 11*) gene has the flowering keyword. TNG67 genes with the keyword heading date include OsTNG03g003615 and OsTNG03g003616 (*plastochron 3*), OsTNG04g000168 and OsTNG04g000220 (*growth-regulating factor 1*), OsTNG05g002262 (*Heme Activator Protein like 1*), OsTNG06g001297 and OsTNG06g001298), and OsTNG08g000038 (*Early heading date 3*).

We searched keywords with a word grain that describes characteristics of grains. These are grain yield, grain size, grain filling, grain length, grain number and grain quality. For TNG67 genes associated with grain yield, we found OsTNG02g000469 (*leucine-rich repeat receptor kinase 1*), OsTNG03g003542 (*Oryza sativa photosystem 1-F subunit*), OsTNG05g002047 (*lactase 15*), OsTNG10g002261 (*chlorophyllide a oxygenase 1*), and OsTNG11g001304 (*vap-related suppressor of too many mouths 1*). Others have the grain size keyword: OsTNG06g000603 (*OsDA1*), OsTNG06g003029 (*FD Transcription Factor 2*), and OsTNG11g000417 (*Cyclin-T1;3*). Interestingly, for the grain length keyword, there are three TNG67 genes that have the same blastn best hit among the genes of Nipponbare. This is Os03g0171300, which encodes for PGL1 or *positive regulator of grain length 1*. When it comes to grain number, OsTNG02g000726 (*Cytochrome P450 71D8L*) and OsTNG11g001569 (*lax panicle 2*) are found. We also found one gene that has the keyword grain quality: OsTNG10g002261 (*chlorophyllide a oxygenase 1*). The same goes with OsTNG08g000885 (*Rb2-like*), which is tagged with the grain filling keyword.

Eight TNG67 genes have a “drought” term: OsTNG01g004606 (endo-1, 3-β-glucanase 1), OsTNG02g003739 (*ethylene-responsive element-binding protein 1*), OsTNG03g003551 (*actin depolymerizing factor 2*), OsTNG03g003603 and OsTNG03g003604 (*calcium-dependent protein kinase 10*), OsTNG03g003563 (*phytochrome interacting factor-like 13*), OsTNG04g001628 (*OsZEP*), OsTNG05g002230 (*b-ZIP transcription factor 42*), OsTNG06g000592 (*anther indehiscence-1*). Two of these 8 genes, OsTNG03g003603/OsTNG03g003604 and OsTNG05g002230, have the term for drought tolerance. The latter is also associated with drought stress. Two other genes, OsTNG03g003563 and OsTNG06g000592, also have the term for drought stress.

We also checked the funRiceGenes keywords that are associated with the TNG67 genes that were derived from TN1, but not found in Nipponbare (see Table S5). Chromosome 6 has OsTNG06g002278, which codes for “*SNF7 component of ESCRT-III complex*” and has the R protein keyword, while OsTNG07g000241 in chromosome 7 has “*Ethylene-insensitive protein 2*” and is associated with blast resistance. OsTNG09g000750 (Rb2-like) in chromosome 9 was tagged with the grain filling keyword, while OsTNG12g000004 (*pleiotropic drug resistance 2*) in chromosome 12 was found to be tagged with yield keyword in funRiceGenes.

## Discussion

We have achieved a chromosome-level assembly of TNG67. TNG67 is largely a *japonica* cultivar because its direct parents are Tainung 61 and a descendant of the cross of Taichung Shih 138 and Tainung 61 (see Figure 1), which are both of *japonica* type. The only source of *indica* genetic material for the TNG7 genome is TN1 (see Figure 1), the genome of which was assembled by Panibe et al. (2021). Except for the genome of TNG67, those of all other *japonica* cultivars in the pedigree (see Figure 1) have not been sequenced. It could have been better if at least one *japonica* parent of TNG67 had its genome assembled because it could serve as a reference for the assembly process. This problem was solved through the use of the high-quality Nipponbare T2T genome produced. The Nipponbare reference genome was used to order and orient the TNG67 scaffolds into a chromosome-level assembly after the Canu *de novo* assembly of the ONT long reads and its subsequent scaffolding (see Table 2). We mapped back the Illumina 150×2 bp short reads with a 45x coverage, and it had a mapping rate of 99.36% better than the Nipponbare T2T genome (99.14%), and TN1’s (98.30 %), supporting the hypothesis that TNG67 is more of a *japonica* variety. The TNG67 genome also has a complete BUSCO rate of 98.4%, comparable to Nipponbare T2T genome’s 98.6% and TN1’s 98.7%.

Using progressive Mauve, we determined genomic segments that are common among TNG67, Nipponbare and TN1 (see Table 6). We obtained information from the backbone output file of progressive Mauve with the three cultivars as input in one run of the software. We were able to obtain not only the genomic segments shared by the three cultivars, but also the segments shared by TNG67 and Nipponbare, by TNG67 and TN1 and by Nipponbare and TN1 (see Table 7). A total of 10,706 TNG67 genes overlap with the backbone segments of the TNG67-Nipponbare pair and were considered to be genes that from the *japonica* ancestry. Also, 1,082 TNG67 genes that overlap with the backbone segment of the TNG67-TN1 pair were deemed to be from the *indica* ancestry (see Table 9). By checking genes that exclusively have *japonica* or *indica* ancestry, we were able to infer that 2,055 of the 10,706 TNG67 genes were from the *japonica* ancestry but were not found in TN1 and 262 of the 1,082 TNG67 genes were from the *indica* ancestry but were not found in Nipponbare.

To explain why TNG67 is highly susceptible to the blast disease, we used the set of cloned R genes of Mahesh (2016) and compared them to their counterparts in TNG67. By aligning a cloned R gene and its orthologue in TNG67, we were able to decide whether the R gene is present or absent in the TNG67 genome and were also able to detect mutations. TNG67 is highly susceptible to the blast pathogen probably because some R genes are lost in TNG67 or have mutated. The lost R genes in TNG67 that originated from *japonica* cultivars are *Pi5-1* and *Pi25,* while the mutated R genes are *Pi21*, *Pib*, *Pi-ta*, *Piz-t*, *Pi37*, *Pik-m-TS1*, *Pit*, *Pi5-2*, *Pb1*, *Pish*, *Pia (RGA4)*, *Pik-p-1*, *Pik-p-2*, *Pik-1*, and *Pi64*.

The loss of an R gene that can pyramid with other R genes may cause a serious reduction in the blast resistance of TNG67. One such R gene is *Piz-t* (Xiao N et al. 2017). For *Pi-ta*, it has proven additive resistance with *Pi46* (Xiao W et al., 2016).

Debnath et al. (2018) tallied a list of R genes that pyramided with other R genes. The list of R genes that were mutated or lost in TNG67 included *Pi-ta*, *Pib*, *Pish*, *Pi21*, and *Piz-*t. *Pi-ta* had the largest number of pyramided genes: *Pi1*, *Piz-5*, *Pi2*, *Pi9*, and *Pi54*. *Pib* pyramided with *Pid1*, *Pita2*, and *Pish*. *Pi21* pyramided with *Pi34*, *QBR4-2*, and *QBR12-1*. *Piz-t* pyramided with *Pi9* and *Pi54*. *Pish* pyramided with *Pib* only. We suspect that the loss of *Pi-ta* incurs the greatest reduction in blast resistance because Pi-ta can pyramid with five R genes, some of which can also have an additive effect with other R genes. For example, *Pi54* can pyramid with either *Pi9* or *Piz-t* (Xiao N et al. 2017), thus increasing the number of R genes that can combat blast.

Rice is a short-day (SD) plant, which means it flowers only when the photoperiod or day length is shorter than 12 hours. Such a plant is said to be photoperiod sensitive (PS). If a rice cultivar flowers even if the photoperiod is longer than 12 hours (long-day or LD), it is called photoperiod insensitive (PI). TNG67 is a hybrid of *japonica* and *indica* cultivars, but its direct parents are *japonica*. So, TNG67 should behave like a *japonica*, which is photoperiod-sensitive. However, thanks to its *indica* ancestor TN1, which is PI, TNG67 has inherited genes that enable it to flower under LD conditions. To understand the reason, we compared the photoperiod genes of TNG67 to their equivalents in Nipponbare and TN1.

Our analysis classified TNG67 photoperiod genes into five groups: 1) similar to both Nipponbare and TN1; 2) more similar to TN1 than to Nipponbare; 3) more similar to Nipponbare than to TN1; 4) unique genes (different from both Nipponbare and TN1 genes); and 5) hybrid genes (hybrids of Nipponbare and TN1 genes) (Table 4). The second and the fifth groups are most interesting to us.

For the 12 phototoperiod genes of TNG67 in Group 2 (Table 4), we searched their orthologs in the regulatory network for LD (Vicentini et al., 2023). Looking for genes that are involved in the control of flowering under LD conditions, we found PRR37, VIL2, GI, Ehd2 (RID1), MADS50, and Ehd4, which are major inhibitors of the flowering regulatory network in rice. Four genes [Hd1 (Heading date 1), Ghd7 (Grain number, plant height, and heading date 7), Ghd8, PRR37 (pseudo-response regulator 37)] are essential for LD, and we expected them to be active in *indica* cultivars, which grow typically in warm tropical regions, i.e., under LD. We got a hint from the review paper of Vicentini et al. (2023) where a gene regulatory network was illustrated to show the interactions between genes under both LD and SD conditions. Central to the network is Ehd1, which promotes the RFT1 and Hd3a florigens under both SD and LD. Inhibit Ehd1 and flowering is repressed. Promote it and flowers are produced.

PRR37 is a major inhibitor of Ehd1 under long-day conditions and causes a delay in flowering. We hypothesize that TN1 (*indica*) PRR37 is an ineffective inhibitor of Ehd1. Another inhibitor of Ehd1 is Ghd8 (aka Hd5 or DTH8), which was classified as a hybrid photoperiod gene in TNG67 (Table 4). Ghd8 interacts with LFL1, which is inhibited by two genes under LD: MADS50 and VIL2. These two genes in TNG67 are more similar to TN1 than to Nipponbare.

GI is a major activator of Ehd1 under SD. It also promotes Hd1 under LD, which inhibits Ehd1. Because we found GI to be similar to its orthologue in TN1, its activity in TNG67 is likely for LD rather than for SD, likely resulting in flowering under LD. Ehd2 (aka RID1) and Ehd4 are two major activators of Ehd1 under both LD and SD, so they may help induce flowering in TNG67 under LD.

It is also important to take note that in our analysis, Hd1 and Ghd8 are categorized as hybrid photoperiod genes of TNG67. Hd1 is promoted by GI and PRR37, both of which are more similar to TN1 than to Nipponbare. We suspect that TN1 and TNG67 GI and PRR37 cannot activate Hd1, allowing the non-inhibition of Ehd1. Another possible reason of TNG67’s early flowering during LD is the hybrid nature of TNG67 Hd1 wherein it has features similar to TN1 and Nipponbare. Its similarity to the Hd1 of TN1, an *indica* cultivar, suggests that TNG67 Hd1 has mutations that hinder its function as an inhibitor of Ehd1. The same can be said about Ghd8. PHYB is another hybrid photoperiod gene, which means it has features similar to Nipponbare. Indeed, PHYB reacts to red light, which activates the Ehd1 inhibitors Ghd7 and PRR37. However, the TN1 features of TNG67 PHYB may reduce the function of Ghd7 and PRR37 in TNG67 during LD.

We also want to mention the similarity between TNG67’s and Nipponbare’s Ghd7 protein sequences. The amino acid changes in TN1 indicate that the TNG67 Ghd7 is more similar to its counterpart in Nipponbare, suggesting that the TNG67 Ghd7 protein does not behave like a typical *indica* Ghd7 protein. Amino acid changes in TN1 from glutamic acid to glycine and from aspartic acid to valine suggest loss of possible ionic interactions. Likewise, the change from proline to alanine could affect the secondary structure of the Ghd7 of TN1. These observations support our hypothesis that the Ghd7 of TNG67 is similar to that of Nipponbare.

From the above discussions, we judge that PRR37, VIL2, GI, MADS50, Hd1, Ghd8, and PHYB are good candidates for the LD in TNG67. Ehd2 and Ehd4 are major activators of Ehd1, so we do not expect them to be involved in the flowering of TNG67 under LD. In addition, the Ghd7 of TNG67 is closer to Nipponbare Ghd7 than to TN1 Ghd7, suggesting that the Ghd7 of TNG67 functions like that of a *japonica* cultivar, which flowers early under SD.

Among the above list of 7 candidate genes, to find the best ones that may explain TNG67’s insensitivity to photoperiod, we examined the domains of their protein sequences and checked whether there are missing domains or domain mutations. We used the hmmer of HMMER version 3.1b2 to detect the domains. We then used the ali coords of the output file of hmmscan to obtain the coordinates where the domain is located in the Clustal alignment.

We found the CCT domain of PRR37 to be identical among the three cultivars (see Table S6). The same is true for VIL2, where the plant homeodomain finger protein motif is 100% conserved among the three cultivars. For GI, no domain was detected for the three cultivars, but there is an exon loss located near the tail-end of the gene in TNG67 and TN1, which might render GI non-functional in TN1 and TNG67. For MADS50, the K-box domain is identical among the three cultivars, but the serum response factor SRF-type transcription factor (DNA-binding and dimerization domain) is missing in both TNG67 and TN1.

Some hybrid photoperiod genes exhibit mutations. For Hd1, the zf-box misses 12 amino acids in Nipponbare, while TNG67 and TN1 have only an amino acid difference (histidine vs. tyrosine) at position 106. On the other hand, the CCT domain in Hd1 is incomplete in TN1, but is the same in TNG67 and Nipponbare, which might imply that Hd1 does not contribute to TNG67’s insensitivity to photoperiod. For Ghd8, the histone-like transcription factor (CBF/NF-Y) and archaeal histone domain is similar between Nipponbare and TNG67, but in TN1 it lacks a few amino acids and shows amino acid changes in the said domain. For PHYB, TNG67 and TN1 are similar in the PAS fold, GAF domain, phytochrome region, PAS domain, and His kinase A (phospho-acceptor) domain, although there are a few amino acid mutations in TN1. The domains that have point mutations in TN1 are the phytochrome region and the GAF domain. In contrast, TNG67 and Nipponbare have similar amino acids at the residue sites where TN1 PHYB has mutated. However, a domain that has the full name of histidine kinase-, DNA gyrase B-, and HSP90-like ATPase (see Table S6) is present in both TNG67 and TN1, but shorter in Nipponbare. Thus, TNG67’s PHYB is likely more similar in function to TN1’s PHYB than to Nipponbare’s PHYB.

The above observations suggest that MADS50 and PHYB are the best candidates for further study on why TNG67 is photoperiod insensitive and that GI may also be a good candidate. The reasons are: 1) Both TNG67 and TN1 MADS50s lack a domain that is present in Nipponbare; 2) TNG67 and TN1 GIs both lack an exon that is present in Nipponbare; and 3) the PRR37 and VIL2 domains are identical among Nipponbare, TNG67 and TN1, so they are not good candidates.

We also studied the grain size genes of TNG67 and compared them to their counterparts in Nipponbare and TN1. Based on the groupings of the genes (Table 3), we suspect the TNG67 genes similar to their counterparts in Nipponbare to be the reason why TNG67 has round and short grains as in *japonica* cultivars. These genes are GW6a (grain weight on chromosome 6-a), GW6 (grain width 6) and WTG1 (wide and thick grain 1). For those grain size genes of TNG67 that are more similar to TN1, we do not expect them to contribute to TNG67’s round and short grains, because TN1, which is an *indica* cultivar, has long and round grains. Those TNG67 genes that are similar between TN1 and Nipponbare would have similar effects on the grain size. Just like the photoperiod genes, hybrid genes may carry some mutations that alter the functions of the genes. These are *GW2* (grain weight 2), *MADS1* (MADS box gene 1), and *GW8* (grain-width 8). We checked the domains of these hybrid genes. In *GW2* no domain was detected for the three cultivars. For MADS1, TNG67 and Nipponbare have the same K-box region, but in TN1 this domain is truncated at the N-terminal, with an inserted sequence of amino acids before the domain. In GW8, the SBP domain is intact among the three cultivars. Thus, TNG67 GW2 and MADS1 are likely responsible to the short grain in TNG67.

## Conclusion

In this study, we sequenced, assembled and annotated the TNG67 genome. The assembly includes 12 chromosomes and some unplaced contigs, leading to a total assembly size of ∼409.1 Mb. We predicted 38,938 genes in the TNG67 genome, larger than the number (37.748) for Nipponbare. Further analysis of the genome revealed 527 R genes, which potentially confer resistance to the rice blast disease. Most of the popular *japonica*-derived R genes in TNG67 were mutated, which may explain why the TNG67 genome is susceptible to the blast fungal pathogen. To know why TNG67 has round and short grains like those from *japonica* cultivars, we investigated 20 grain size genes and identified several candidate genes, which are more similar to their counterparts in Nipponbare than in TN1. We also studied 38 photoperiod genes to identify candidate genes potentially responsible for the photoperiod insensitivity of TNG67 and selected three best candidate genes for functional validation. In conclusion, this study provides much insight into the genome of TNG67, a historically very important rice cultivar of Taiwan. It increases our understanding of why TNG67 has grain size and shape similar to Nipponbare but photoperiod insensitivity as in TN1, an *indica* cultivar

## Materials and Methods

### Illumina sequencing

We sequenced the TNG67genome with a total coverage of 209.49x (Table 10), using Illumina HiSeq, Illumina MiSeq and the Oxford Nanopore Technologies (ONT). The Illumina paired-end (PE) reads can be subdivided into two: (1) short PE reads with 300×2 bp and 150×2 bp; they have a coverage of 18.25x and 44.89x, respectively. (2) PE reads with long inserts; they have a read length of 150×2 bp with a coverage subtotal of 36.14x. Another type of read was the mate-pair reads with 150×2 bp and insert sizes of 2 to 4 kb, 4 to 6 kb, and 6 to 10 kb, providing a subtotal coverage of 22.35x.

**Table 10.** Sequencing Data. . The Nipponbare reference genome size used in computing the coverage of the TNG67 reads is 373,245,519 bp.

| Sequencing technology | Type of read | ID | Read length (bp) | Total base pairs | coverage |
| --- | --- | --- | --- | --- | --- |
| Oxford Nanopore Technology | raw long reads | - | 3,048 | 32,895,748,276 | 88.13 |
| Illumina MiSeq | short PE reads | LGC-IS01_604bp | 300x2 | 6,812,370,823 | 18.25 |
|  |  | SG_DE51_351bp | 150x2 | 16,756,572,583 | 44.89 |
| Illumina<br>HiSeq | long<br>PE<br>reads | SG_DE01_C_1062bp | 150x2 | 8,850,813,397 | 23.71 |
| Illumina<br>MiSeq |  | SG-DE01-C-1062bp | 150x2 | 589,024,053 | 1.58 |
| Illumina<br>HiSeq |  | SG_DE01_D_1700bp | 150x2 | 3,521,485,850 | 9.43 |
| Illumina<br>MiSeq |  | SG-DE01-D-1700bp | 150x2 | 529,529,208 | 1.42 |
| Illumina<br>HiSeq | Mate-pair<br>reads | SG_DE01_MP2_4kb | 150x2 | 2,118,852,129 | 5.68 |
|  |  | SG_DE01_MP4_6kb | 150x2 | 3,126,238,373 | 8.38 |
|  |  | SG_DE01_MP6_10kb | 150x2 | 3,093,735,761 | 8.29 |

TOTAL 209.76x

### Nanopore sequencing

The genomic DNA of embryonic leaf was extracted via standard phenol-chloroform extraction protocol. The extracted genomic DNA was checked in 0.8-1% agarose gel and concentrated by the size selection of KAPA Pure Beads (Cat# KR1245, Kapa Biosystems, Wilmington, MA, USA). Library preparation was carried out using the ligation sequencing kits (Oxford Nanopore Technologies) SQK-LSK109 for sequencing on R.9.4.1 flowcells. The concentrated DNA (1∼3 μg) was end-repaired and ligated with AMX adaptors (SQK-LSK109) via a KAPA Hyper Prep Kit (Cat#KR0961, Kapa Biosystems, Wilmington, MA), following the manufacturer’s instructions. The constructed library was premixed with the LB and SQB buffer (SQK-LSK109), loaded in four flowcells (R9.4.1; FLO-MIN106), and sequenced by MinION devices for 48-72 h in laboratory. the fast5 raw reads were basecalled by Guppy 4.4.0 with default fast mode. The ONT long reads have an average read length of 3,048 bp with a coverage of 88.13x. The passed fastq reads were further assembled using Canu v2.2.1 (Koren et al. 2017).

### Quality trimming of sequencing reads

Trimmomatic version 0.39 was used to trim the Illumina 300 bp x2 PE reads and 150 bp x 2 PE reads with a minimum length of 300 bp and 75 bp, respectively. The last base of the reads was also removed to make them 300 bp and 150 bp, respectively. All adapter sequences were merged before running the software to save time. The output file is an interleaved fastq and were separated into read 1 and read 2 using reformat.sh script of bbmap via default options. The separated reads were processed by Trimmomatic version 0.39 using the same adapter fasta file used in trimming the paired-end reads (parameters: ILLUMINACLIP:adapters.fa:2:30:10 LEADING:20 TRAILING:20 SLIDINGWINDOW:4:20 MINLEN:30 CROP:150). The paired-end, long paired-end and mate-pair reads were further processed by sga for base correction (parameters: sga preprocess --pe-mode 1 (default); sga index -a ropebwt --no-reverse (default settings used for long PE and mate-pair reads; ropebwt option not used for the short PE reads); sga preqc (default); sga correct --learn (default)).

### TNG67 genome assembly

The assembly procedure is shown in Figure 2. The nanopore reads were assembled by Canu version 2.1.1into contigs, which were corrected by Pilon version 1.23 in one iteration. The corrected contigs were then scaffolded by SSPACE Standard version 3.0 using various long paired-end and mate-pair reads. Gaps formed when using SSPACE were largely closed by PBJelly (PBSuite version15.8.24). The remaining gaps were reduced further by GapCloser version 1.12 and GapFiller version 1.10 using paired-end reads. At this point, the TNG67 assembly still has 2,206 scaffolds, so the Nipponbare reference genome by IRGSP was utilized to infer chromosome-level scaffolds. The RagTag version 2.1.0 reference-guided assembler ordered and oriented the TNG67 scaffolds into a set of 12 chromosomes against the chromosomes of Nipponbare, with minimap2 version 2.22 as the aligner used. One round of Pilon was used to correct the assembly to produce the final TNG67 genome sequence. We saved this first version of the TNG67 assembly as TNG67_rice_v1.0.fasta. After submitting to NCBI, there were contigs that were detected to be contaminants such as mitochondrion sequences and eukaryotic virus sequence. We removed the contaminant sequences and any gaps flanked by them. Next, we further improved the TNG7 assembly using the Nipponbare T2T genome as reference (Genbank accession #: GCA_034140825.1). To do the re-assembly, we used the component scaffolds and contigs of TNG67_rice_v1.0.fasta, prior to its improvement via RagTag. We used these contaminant-free scaffolds and contigs of TNG67 as input to the reference-guided re-assembly with the Nipponbare T2T as reference. We used RagTag software again using default options, and after that we polished the genome with one run of Pilon using Illlumina short paired-end reads. After uploading to NCBI for submission, contaminants were still detected, and we redo the RagTag assembly and saved it as TNG67_rice_v1.2.fasta

### TNG67 genome annotation

The TNG67 genome was annotated using the two-pass MAKER approach as in Panibe et. al (2021). CEGMA version v2-2.5 and SNAP version 2013-11-29 were first executed before doing the first run of MAKER. After that, further training was done using Augustus version 3.2.3 and SNAP again. Transcripts (CDS + UTRs, version 2021-11-11) and proteins (version 2021-11-11) of the Nipponbare genome were downloaded from the RAPDB website. For the TN1 proteins and transcript sequences, they were extracted using gffread, version 0.12.7 on default settings. To be used as in input in the MAKER run, the TN1 and Nipponbare transcript sequences were merged. The same was done for the protein sequences. After MAKER finished running, the utility scripts gff3_merge was executed to get the gff file and fasta file, respectively. New gene names were generated using the maker_map_ids script. After two rounds of re-assemblies, the final annotation gff was produced by using Liftoff v1.6.3 with options -exclude_partial -overlap 1 (Shumate & Salzberg, 2020).

### BUSCO score and Blast2GO annotation

We checked the genome assembly completeness using BUSCO v5.2.2 using the poales_odb10 lineage set (Manni et al., 2021). Functional annotation of the TNG67 genes were done in three steps: Blasting, Mapping and Annotation. The TNG67 genes were first blasted via blastx to the nr database. To make the blast process faster, the fasta file was divided into five parts producing separate xml outputs. The xml files were used as input in Blast2GO v5.2.5 free version as blast xml legacy files. After all the blast xml results were loaded, GO Mapping in Blast2GO was executed using the Basic Mode. Default settings were also used for the Annotation step.

### R gene prediction

To predict the R genes of TNG67, hmmscan (Eddy 1998) was executed to detect the presence of NB-ARC domain with an LRR repeat, through the Pfam database (Mistry et al. 2021). The input was the TNG67 proteins derived from gffread (Pertea and Pertea 2020), version 0.11.8 (default options). If only an NB-ARC but no LRR was present, the tool NLR-Parser v1.0 (Steuernagel et al. 2015) was used to find LRRs. Pfam domains were predicted using hmmscan of HMMER 3.1b2 via default options. The database used was the Pfam-A.hmm file of Pfam v30.0 (El-Gebali et al. 2018). Outputs were saved using the --tblout, --domtblout and –pfamtblout parameters. A Perl script was written to process the pfam table output file. For the prediction of R genes by NLR-Parser, mast (Bailey and Gribskov 1998) was executed to get an xml file. This xml file served as input in the NLR-Parser command. After running NLR-Parser, the output was converted into a csv file through a script. Finally, the results of the Pfam domain prediction and NLR-Parser were integrated. The locations of the NB-LRR R genes in the 12 chromosomes of TNG67 were illustrated using CViT (Cannon et al., 2015) (Figure S2). In drawing the figure, the chromosomes were scaled in kilo base pairs (kbp). Other settings were inputted in the cvit_kb.ini parameter file.

To identify the equivalent of the TNG67 R genes with respect to the Nipponbare reference genome, we inputted their proteomes into OrthoFinder (Emms and Kelly 2019). The Nipponbare IRGSP-1.0 (GenBank accession #: GCA_001433935.1) proteins version 2022-03-11 were downloaded from the RAPDB website, while the TNG67 proteins used were the same as those used in the R gene prediction.

### GenePainter

Protein sequences of the target genes were concatenated and were aligned using Clustalw v2.1. Parameters: -type=protein -output=FASTA -align. To create the gff files of each protein, lines on the gff file, containing the gene name as well the corresponding protein isoforms were extracted from the gff file, and saved as a single .gff file. So, a gff file containing the gene name was first made, and then another gff file containing the protein name was created. These two files were concatenated and saved using protein isoform as the filename.

To create the gene structure diagrams, the Clustal alignment in fasta format was first uploaded in the GenePainter website. Then, the gff files of the proteins that were used in the alignment were uploaded. The Submit button was clicked and the output svg file was downloaded.

### genoPlotR

To create the Nipponbare-T2T-TNG67-TN1 alignments, the chromosome of each genome were aligned via progressiveMauve v2015-02-13 (output parameters: --output --output-guide-tree --backbone-output). A total of 12 Nipponbare-T2T-TNG67-TN1 chromosome alignments were done. The backbone output files of progressiveMauve were processed by a set of genePlotR commands to create the images of the alignment. Available at Figshare (https://doi.org/10.6084/m9.figshare.25241731).

## List of abbreviations

IRGSP: International Rice Genome Sequencing Project
LD: long-day
LRR: Leucine-Rich Repeat
NB-ARC: Nucleotide-Binding domain, Apaf-1, R proteins, and CED-4
NBS: Nucleotide-Binding Site
NB-LRR: Nucleotide-Binding site, Leucine-Rich Repeat
NBS-LRR: Nucleotide-Binding Site, Leucine-Rich Repeat
PI: photoperiod sensitive
PS: photoperiod insensitive
RAP-DB: Rice Annotation Project Database
R gene: resistance gene
SD: short-day
T2T: Telomere-to-Telomere
TN1: Taichung Native 1
TNG67: Tainung 67

## Declarations

## Ethics approval and consent to participate

Not applicable

## Consent for publication

Not applicable

## Availability of data and material

This Whole Genome Shotgun project has been deposited at DDBJ/ENA/GenBank under the accession JAVAIK000000000. The version described in this paper is version JAVAIK010000000. The sequencing reads are deposited under NCBI BioProject PRJNA1002363 and BioSample SAMN36841526. SRA accession numbers for ONT long reds: SRR25757643; Illumina paired-end reads: SRR25757644, SRR25757645, SRR25757649, SRR25757650, SRR25757651, SRR25757652; Illumina mate-pair reads: SRR25757646, SRR25757647, SRR25757648. The GFF (General Feature Format) annotation file, gene sequence, protein sequence, and coding sequence are available at Figshare (https://doi.org/10.6084/m9.figshare.32999885).

## Authors’ contributions

Jerome P. Panibe: genome assembly, analysis of grain size and photoperiod genes, determination of origin of TNG67 sequences whether they were derived from *japonica* or *indica,* funRiceGenes search

Wang Long: MAKER annotation and R gene prediction

Tzi-Yuan Wang: conducted ONT library preparation and sequencing.

Mei-Yeh Jade Lu: designed and led Illumina sequencing experiments

Chang-Sheng Wang: planted the TNG67 rice/advised the study.

Wen-Hsiung Li: designed, advised and supervised the study.

## Acknowledgements

We thanked the High Throughput Genomics Core Facility, Biodiversity Research Center, Academia Sinica, Taiwan for sequencing the TNG67 genome.

