## Supplementary material for "Assembling and annotating the Tainung 67 rice genome to trace its *japonica* and *indica* ancestries": Figure S1-S9

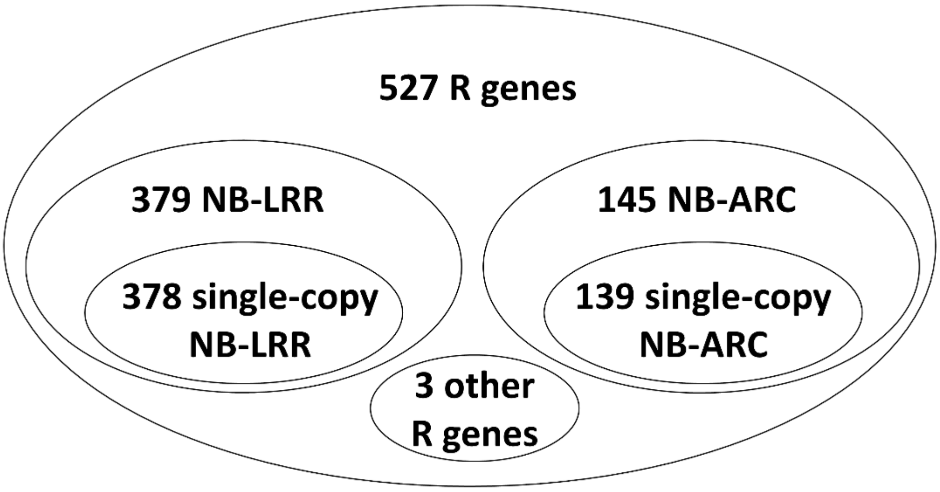


**Figure S1. Summary of predicted R genes.** A total of 527 R genes were predicted in the TNG67 genome.

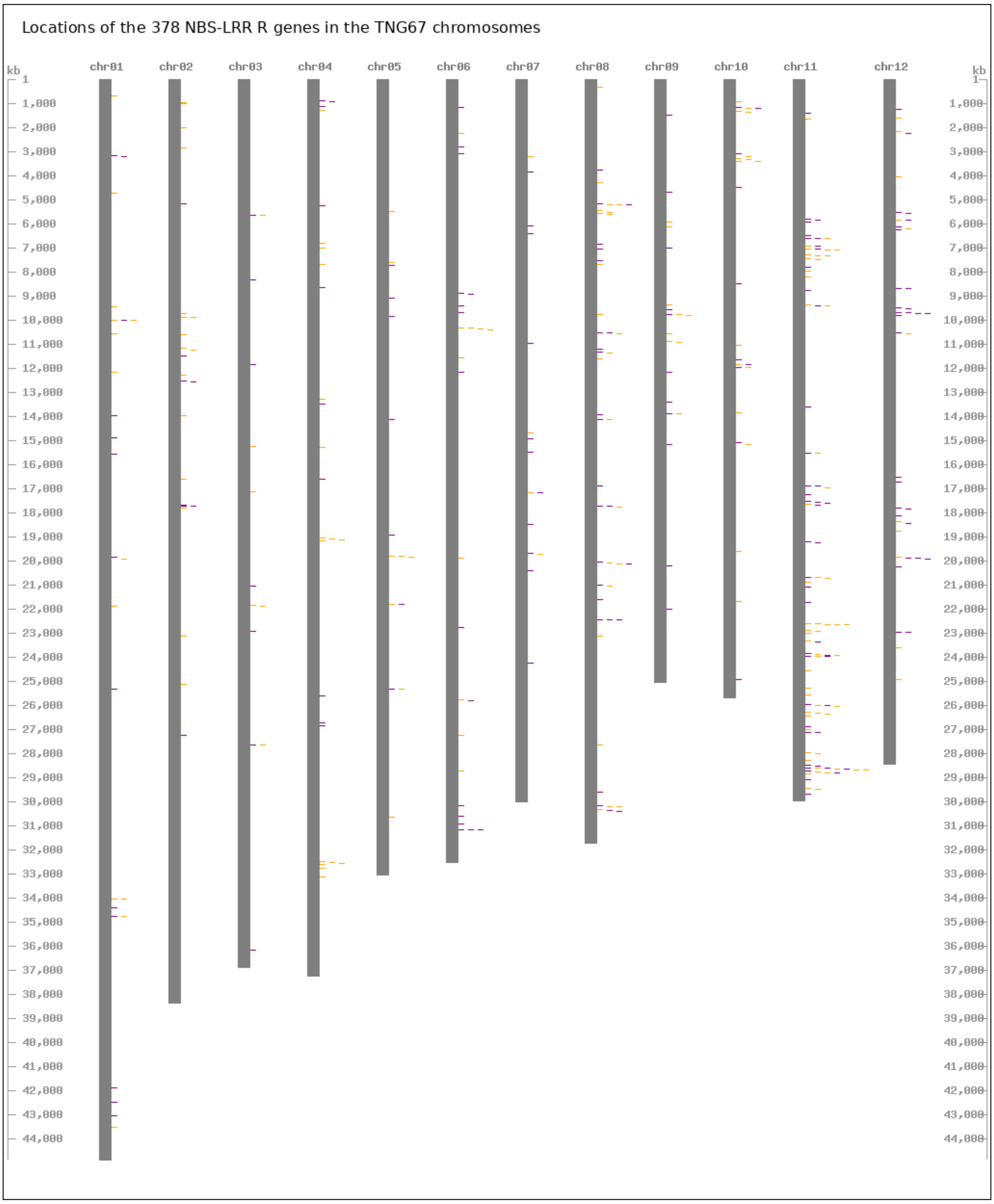


**Figure S2. Locations of the 378 NBS-LRR R genes of TNG67.**

Each orange bar means the gene is on the positive strand, while violet means it is on the negative strand. A scale in the form of a ruler (left and right) was set to have a tick size per 1 kb. For example, chromosome 1 has a length of 44 Mb. Refer to Table S2 to identify the gene(s) in a specific location.

### Analysis of grain size genes of TNG67, Nipponbare and TN1

#### *Grain size genes of TNG67 that are unique*

We next discuss the unique grain size genes of TNG67 (see Figure S3). With the exception of Os03t0183000-03 (because we chose Os03t0183000-02 as the representative of Nipponbare OsLG3), the first and second exons of *OsLG3* in the three cultivars are almost the same (Figure S3a), except with a lone change from glycine to tryptophan for TN1's OsTN3t000728.1, which are both nonpolar. TNG67's OsTNG03t000695.1 is the only gene that has complete 3$^{rd}$, 4$^{th}$ and 5$^{th}$ exons, which makes it a unique gene compared to the *OsLG3* genes in Nipponbare and TN1. For *GSA1*, it can be clearly seen from the gene structure models and the protein sequence alignment that TNG67's OsTNG03t003410.1 is different from its orthologues in Nipponbare and TN1. Therefore, the TNG67 GSA1 gene is unique.

For *GLW7*, the first exon of TNG67's OsTNG07t001568.1 and that of TN1's OsTN7t001695.1 agree but amino acid changes are found on its protein sequence alignment, which are located at the tail-end of exon 1 of the TNG67 gene. On the second exon of TNG67's OsTNG07t001568.1, it is almost a complete loss of its exonic region, making the *GLW7* gene of TNG67 unique.

*a) OsLG3*

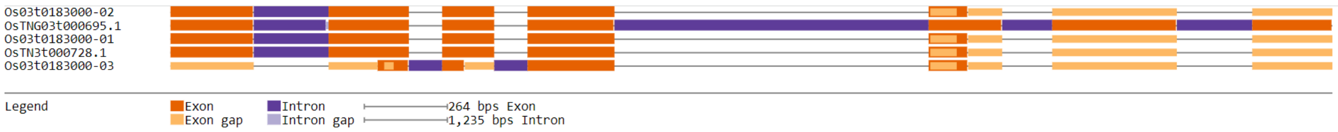

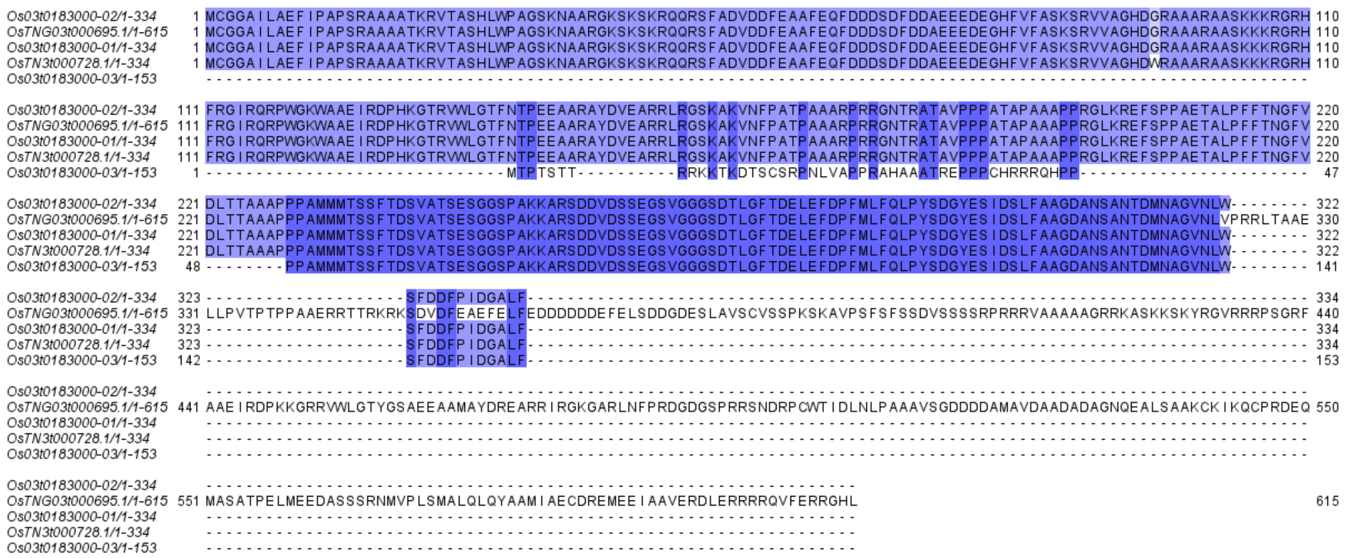
Os03t0183000-02/1-334
OsTNG03t000695.1/1-615
Os03t0183000-01/1-334
OsTN3t000728.1/1-334
Os03t0183000-03/1-153

*b) GSA1*

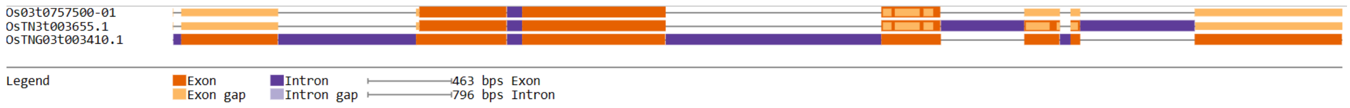
Os03t0757500-01
OsTN3t003655.1
OsTNG03t003410.1
Legend
Exon
Exon gap
Intron
Intron gap
463 bps Exon
796 bps Intron

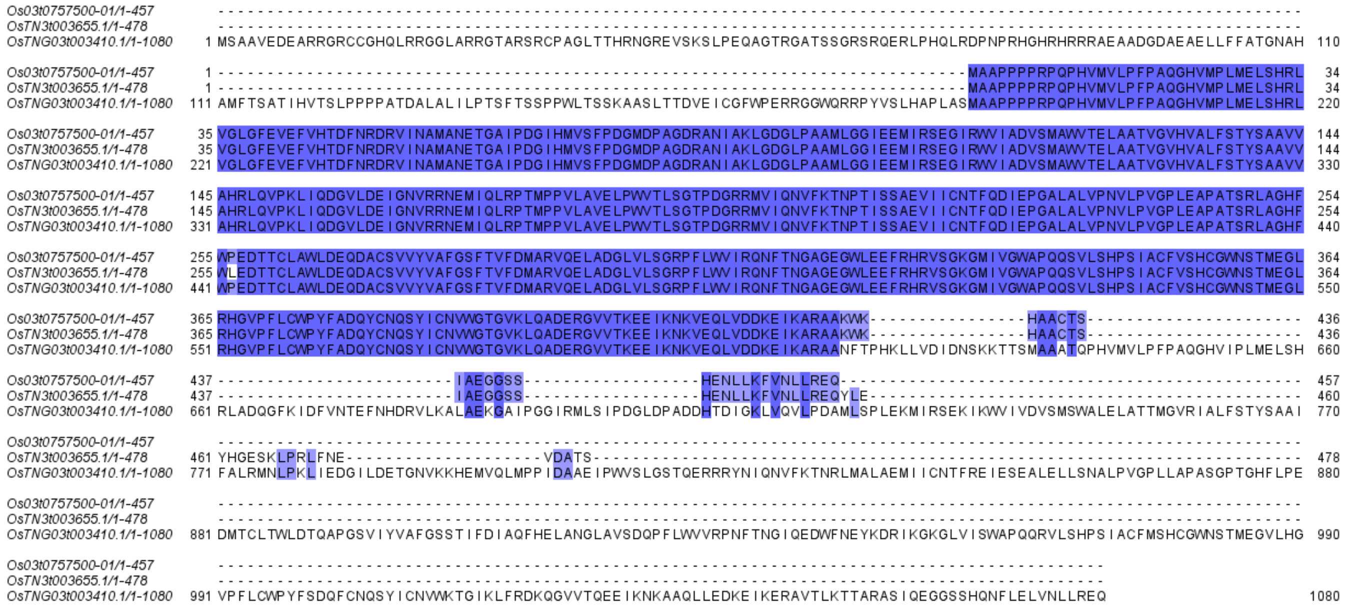
Os03t0757500-01/1-457
OsTN3t003655.1/1-478
OsTNG03t003410.1/1-1080

*c) GLW7*

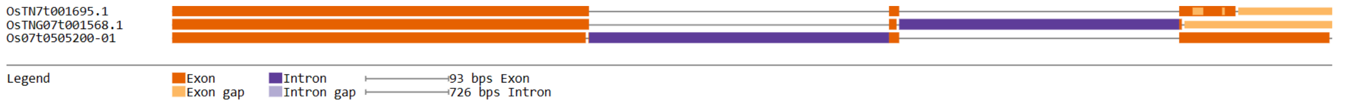
OsTN7t001695.1
OsTNG07t001568.1
Os07t0505200-01
Legend
Exon
Exon gap
Intron
Intron gap
93 bps Exon
726 bps Intron

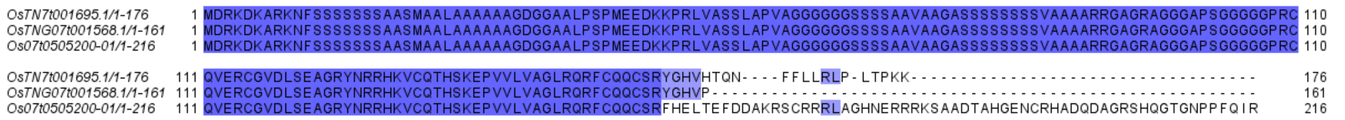
OsTN7t001695.1/1-176
OsTNG07t001568.1/1-161
Os07t0505200-01/1-216

**Figure S3. Gene structure diagrams and protein sequence alignment of grain size genes that are unique in TNG67.** a) *OsLG3*: the TNG67 gene is unique for having exons on the second half of the gene as opposed to the loss of exonic regions in Nippnbare and TN1. b) *GSA1*: the TNG67 gene is unique as compared to Nipponbare and TN1, which are similar to each other. c) *GLW7*: the TNG67 gene has lost most of its last exon as compared to Nipponbare, which has an intact last exon, and TN1, which has some missing parts.

***Grain size genes of TNG67 that are hybrid***

There are also hybrid genes of TNG67 according to our definition above. These TNG67 genes have parts in their gene structure, specifically their exonic regions that are similar to Nipponbare in one part, but similar to TN1 in the other part of gene. A TNG67 gene can also be a hybrid, if there is an exonic region not found in Nipponbare or TN1, but in the other part of the gene, there is a match to either or both of the two other cultivars. *GW2* and *LGY3* are the two hybrid grain size genes of TNG67. For *GW2*, the first exon of TNG67 OsTNG02t001168.1 is present, but missing in its TN1 and Nipponbre orthologues (see Figure S4a). However, in the fifth exon of the TNG67 gene, a portion of its exonic region is missing and this is also the same with TN1's OsTN2t001119.1. For *LGY3* of TNG67, the first half of its exon 1 is missing. It is almost the same with the TN1 orthologue, except for the loss of a tiny exonic region (Figure S4b), which corresponds to two cysteine amino acids in TNG67 OsTNG03t000970.1. Moving along the gene there are two more losses of exonic regions, which are the same between TN1 and TNG67. However, on the fifth and sixth exon, there is another loss of an exonic region, but this time, the loss is found in both TNG67 and Nipponbare. At the start of the GW8 protein sequence alignment (see Figure S4c), TN1 and TNG67 match but they do not start with a methionine, which could be an error in the gene annotation. Meanwhile, TNG67's OsTNG08t002577.1 has a noticeable sequence of amino acids that do not match Nipponbare's Os08t0531600-01 and TN1's OsTN8t002176.1, which are highly similar (see Figure S4c). In contrast to these TN1-Nipponbare matches, there are some amino acids that match between TNG67 and Nipponbare only. These suggest that TNG67 *GW8* gene is a hybrid between Nipponbare and TN1.

a) GW2

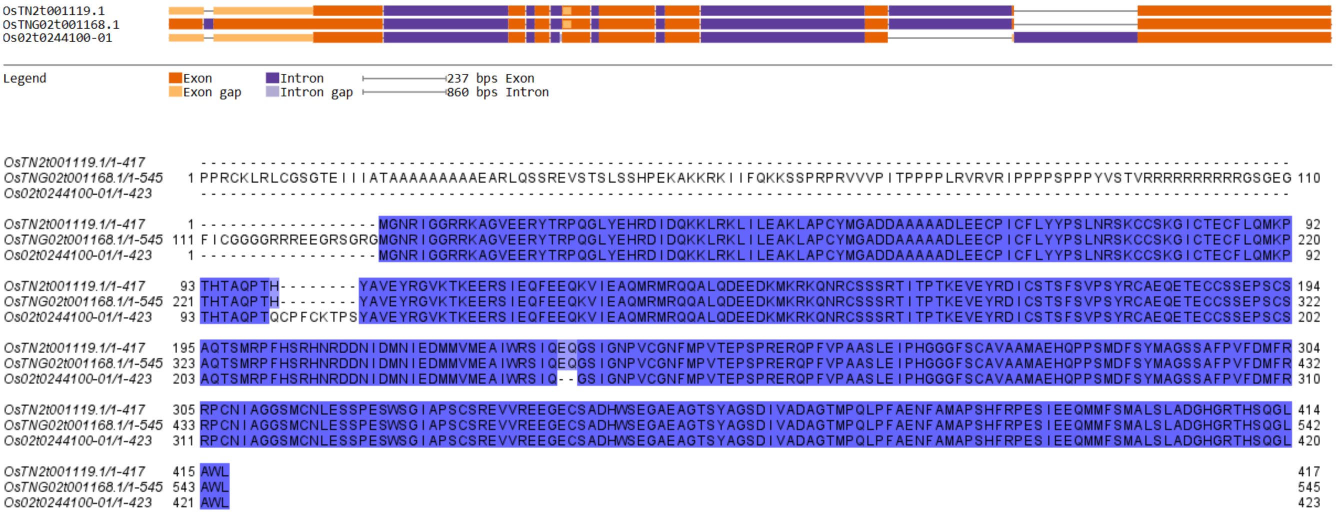


b) LGY3

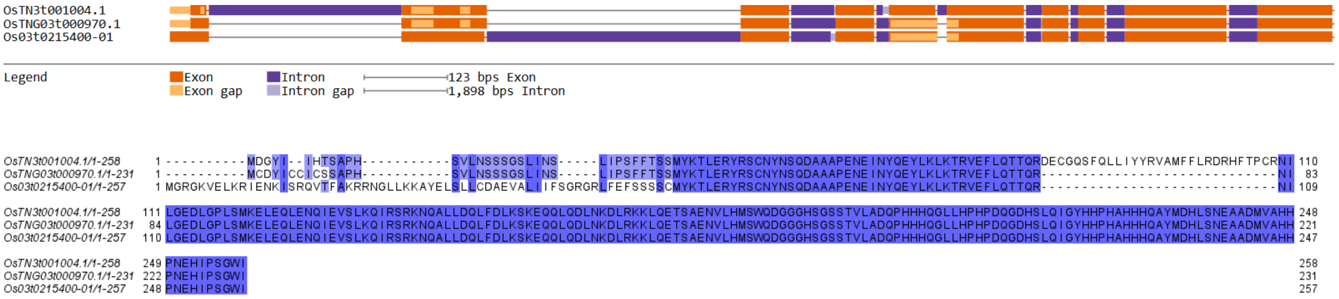


c) GW8

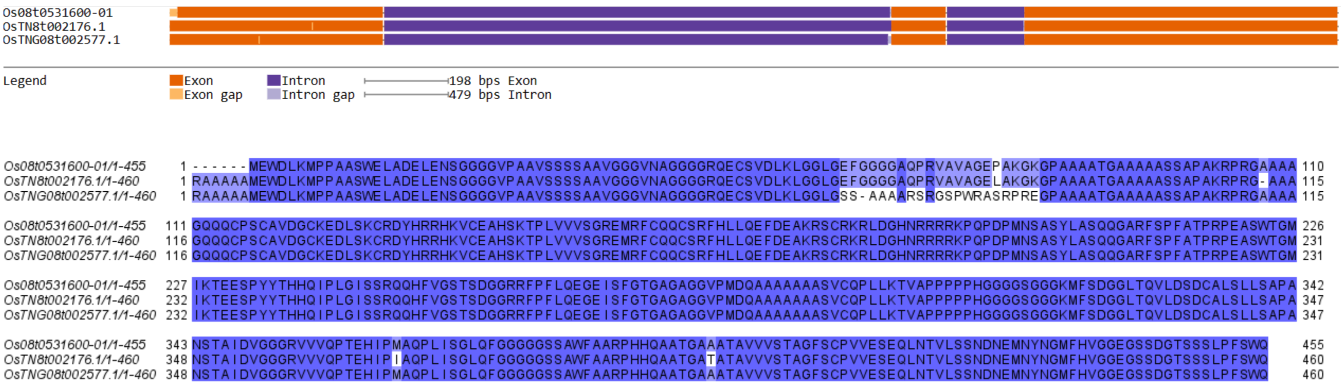


**Figure S4. Gene structure diagrams and protein sequence alignment of grain size genes of TNG67 that are hybrid.** a) *GW2*: TN1 and Nipponbare proteins are similar at the N-terminal part, but from the middle part to the end, TNG67 and TN1 are more similar. b) *LGY3*: TN1 and TNG67 proteins are similar at the N-terminal part, but for the mid-part Nipponbare and TNG67 are more similar, including losses of exonic regions. c) *GW8*: The last two exons are the same, but the first exon shows a loss of an exonic region in Nipponbare and many amino acid changes in TN1.

***Grain size genes of TNG67 that is similar to both Nipponbare and TN1***

Lastly, we consider the protein sequence alignments of the grain size genes that are similar among TNG67, Nipponbare and TN1. The gene structures and the protein sequence alignments of TGW2 (**Figure S5a**) and GL6 (**Figure S5c**) for the three cultivars are like mirror images of each other. Because the genes are very similar, they likely have the same effect on grain size across the three cultivars. For *qGL3*, TN1's OsTN3t002925.1 has an amino acid replacement that changed lysine to histidine. Because both are basic amino acids, the change may not have a significant effect on the TN1 protein. However, for *GS9*, TN1's OsTN9t001370.1 has amino acid mutations that could disrupt its secondary structure --- specifically, the leucine to proline and the proline to leucine change. Proline is a bulky amino acid and could make a dent on the amino acid chain. Hence, this mutation could have a drastic effect on the overall structure of the protein. Likewise, the change from arginine (a charged amino acid) to cysteine (**Figure S5d**) could also affect the 3D structure of OsTN9t001370.1. This could mean a loss of an ionic bond. Furthermore, the cysteine could form a disulfide bond with another cysteine in the protein, thereby affecting the structure of the protein. Thus, the TNG67 and TN1 GS9 genes are more similar to each other than to the TN1 GS9 gene.

a) TGW2

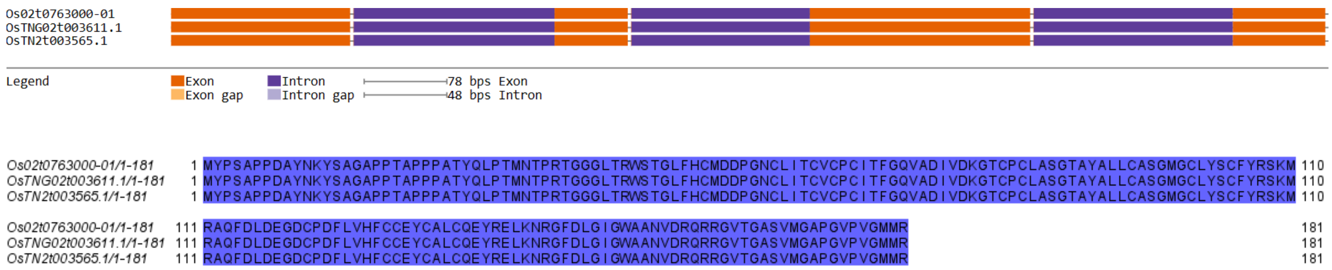


b) qGL3

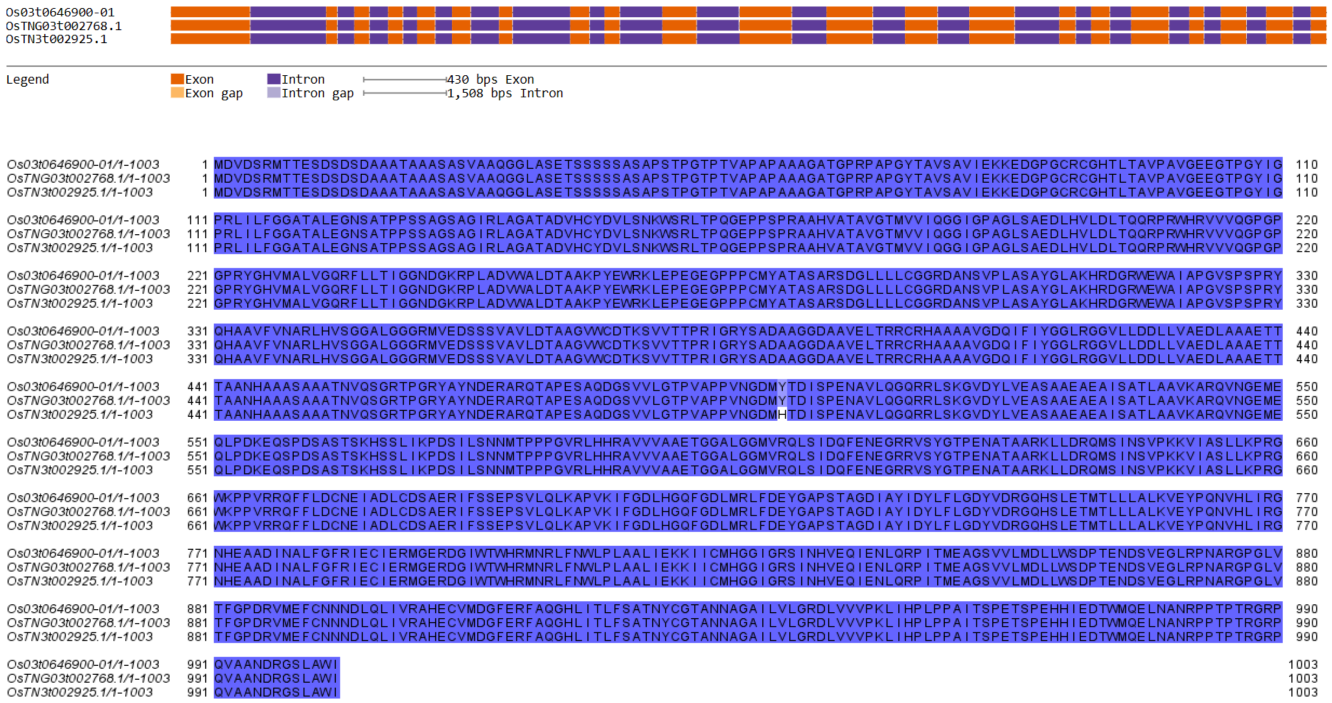
Os03t0646900-01
OsTNG03t002768.1
OsTN3t002925.1
Legend
Exon
Exon gap
Intron
Intron gap
430 bps Exon
1,508 bps Intron
Os03t0646900-01/1-1003
OsTNG03t002768.1/1-1003
OsTN3t002925.1/1-1003

c) GL6

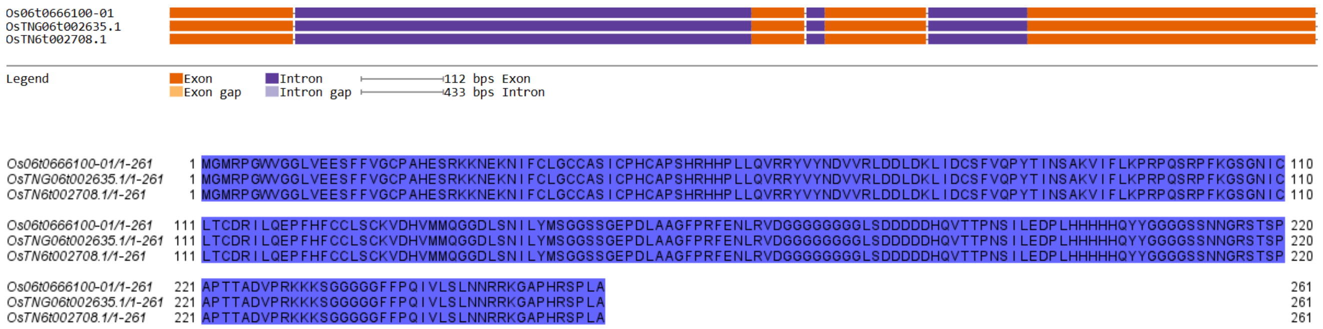
Os06t0666100-01
OsTNG06t002635.1
OsTN6t002708.1
Legend
Exon
Exon gap
Intron
Intron gap
112 bps Exon
433 bps Intron
Os06t0666100-01/1-261
OsTNG06t002635.1/1-261
OsTN6t002708.1/1-261

d) GS9

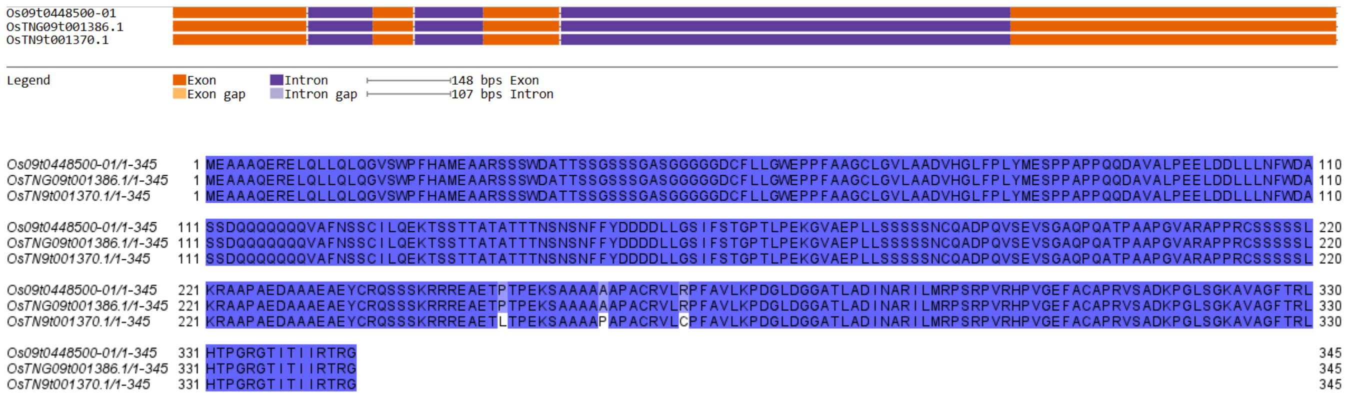
Os09t0448500-01
OsTNG09t001386.1
OsTN9t001370.1
Legend
Exon
Exon gap
Intron
Intron gap
148 bps Exon
107 bps Intron
Os09t0448500-01/1-345
OsTNG09t001386.1/1-345
OsTN9t001370.1/1-345

**Figure S5. Gene structure diagrams and protein sequence alignment of grain size genes of TNG67 that is both similar for Nipponbare and TN1.** TGW2, qGL3 , GL6,GS9. a) TGW2: TNG67, Nipponbare and TN1 are similar with each other. b) qGL3: similar gene structure c) GL6: same for all the cultivars; d) GS9: same for Nipponbare, TN1 and TNG67

**Analysis of photoperiod genes of TN67, Nipponbare and TN1**

***TNG67 is similar to both Nipponbare and TN1***

We now discussed the photoperiod gene of TNG67 that are similar for both Nipponbare and TN1. The first gene is *SE5* (photosensitivity 5). In Figure S6a, the gene structures for the three cultivars are similar. Their protein sequence alignment also shows 100% similarity. Thus, we expect the effect of SE5 in TNG67 to be neutral. For *RFL* (flo-fly homolog of rice), (see Figure S6b), the gene structure models of the three cultivars show a complete set of exons that agree with each other. But an inspection with its protein sequence alignment would reveal an amino acid change from an Aspartic Acid to a Glutamic Acid. The RFL proteins of TNG67 and Nipponbare have the former amino acid, while the TN1 RFL protein has the latter. But because both amino acids are polar and acidic, the effect of the amino acid change is neutral. So we expect the three RFL proteins to have the same function.

For *ETR2*, the gene structure models depict a very small exonic loss at the tail-end of TNG67 and TN1. These exonic loss corresponds to a gap at the end of the alignment (see Figure S6d), in which TNG67's OsTNG09t000885.1 and TN'1 OsTN4t000312.1 agree well. Disregarding the overhanging VLQNN subsequence of Nipponbare's Os04t0169100-02, and looking at the lone amino acid change from Valine to Leucine in TN'1 OsTN4t000312.1 (see Figure S6d), we can say that the ETR2 of the three cultivars are the same. This is because Valine and Leucine are nonpolar and neutral amino acids so we expect the protein sequences among the three to be the same. In *RFT1*, the gene structure models suggest a loss of exonic regions in the first exon of TNG67 and TN1 (see Figure S6e). But the protein sequence alignment shows that the starting amino acid of TNG67's OsTNG06t000423.1 and TN1's

OsTN6t000439.1 is not Methionine (see Figure S6e). This could be due to an error in the annotation, and we suspect start codon of the gene should be upstream and code for the same set of amino acids like Nipponbare's Os06t0157500-01. With the high protein sequence similarity, we hypothesize that the *RFT1* gene of the three cultivars are the same.

a) SE5

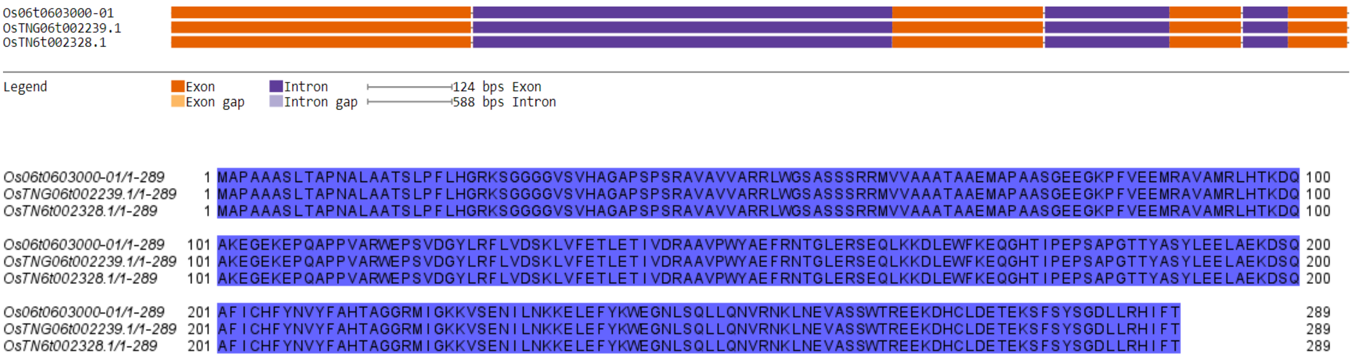


b) RFL

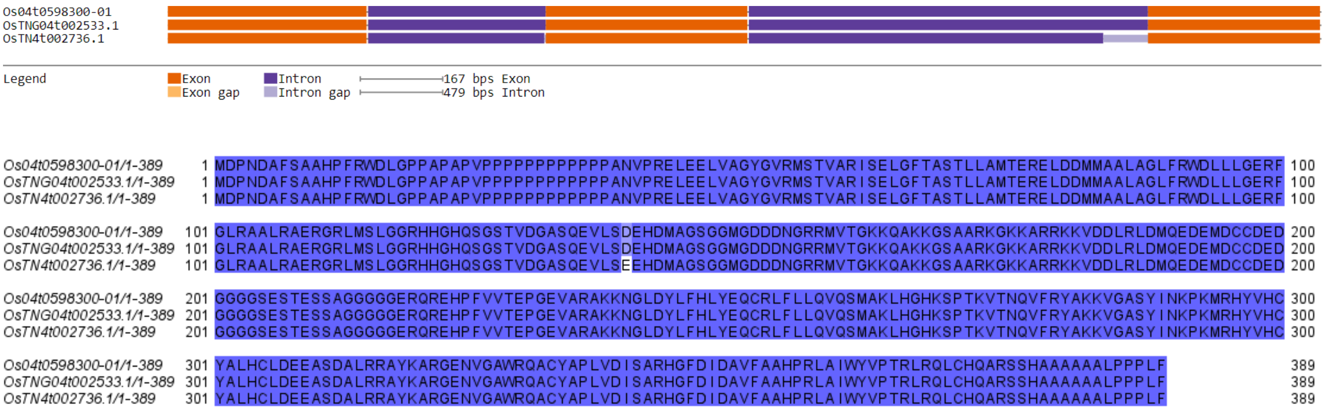


c) pps

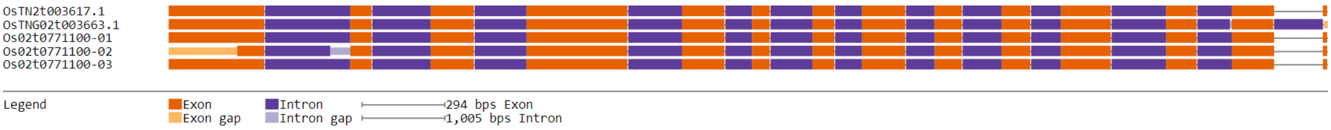

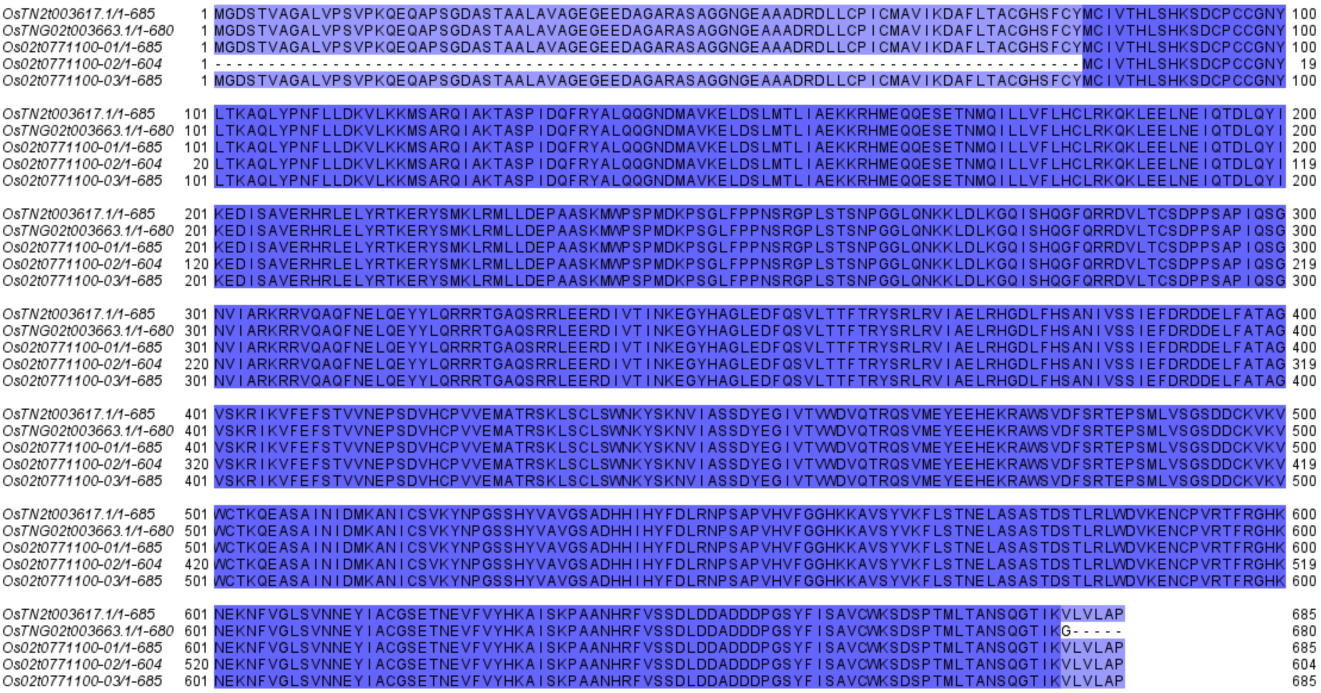
OsTN2t003617.1/1-685
OsTNG02t003663.1/1-680
Os02t0771100-01/1-685
Os02t0771100-02/1-604
Os02t0771100-03/1-685

d) ETR2

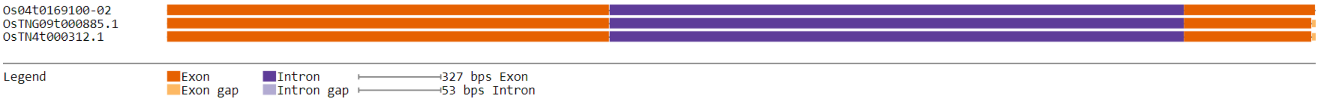
Os04t0169100-02
OsTNG09t000885.1
OsTN4t000312.1
Legend
Exon
Exon gap
Intron
Intron gap
327 bps Exon
53 bps Intron

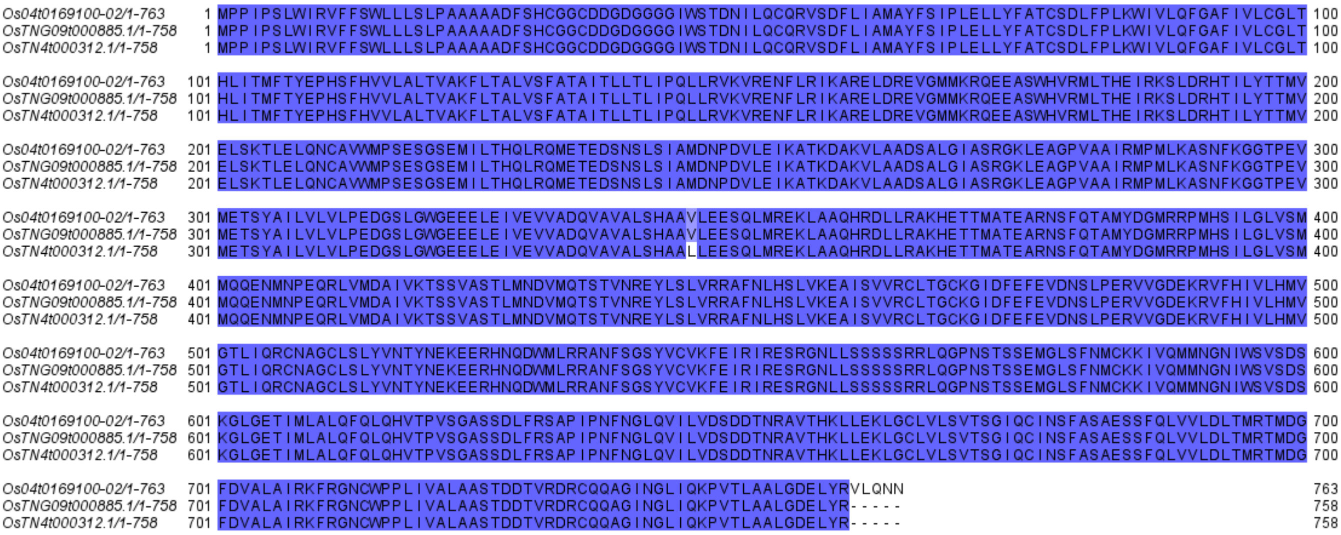
Os04t0169100-02/1-763
OsTNG09t000885.1/1-758
OsTN4t000312.1/1-758

e) RFT1

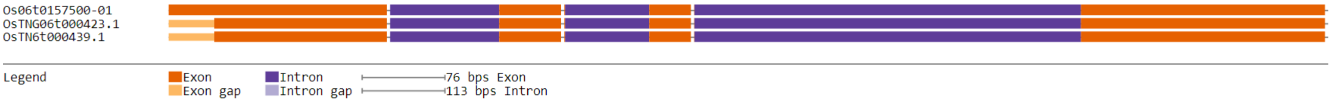
Os06t0157500-01
OsTNG06t000423.1
OsTN6t000439.1
Legend
Exon
Exon gap
Intron
Intron gap
76 bps Exon
113 bps Intron

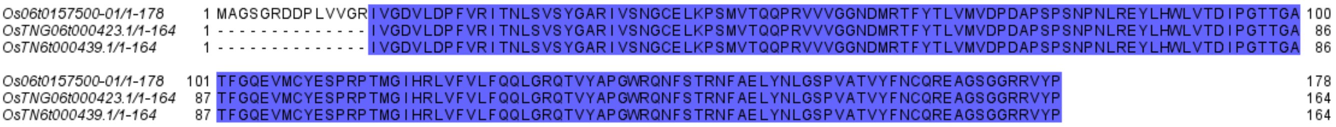


**Figure S6. Gene structure diagrams and protein sequence alignment of photoperiod genes of TNG67 that is similar to both Nipponbare and TN**1. a) SE5, b) RFL, c) pps, d) ETR2, and e) RFT1

***TNG67 is more similar to Nipponbare***

For the TNG67 gene that are more similar to Nipponbare, we found four genes and they are *Hd16* (heading date 16), *OsCOL4* (constans-like gene 4), and *OsDof12* (Dof transcription factor 12). Based on the gene structure models of *Hd16* (see Figure S7a), as well the protein sequence alignments of the gene. OsTNG03t003641.1, Os03t0793500-01, and Os03t0793500-02 agree well to each other, while there is a loss of exonic regions from the 13$^{th}$ to 16$^{th}$ exons for OsTN3t003776.1. With the high similarity of the TNG67 Hd16 protein sequence with its orthologues in Nipponbare, it is safe to say that TNG67's Hd16 functions like its Nipponbare counterpart.

For OsCOL4 (see Figure S7b), there are two amino acid changes, in which TN1's OsTN2t002590.1 is different with its orthologues in TNG67 and Nipponbare. These amino acid changes were a mutation changing Proline to Threonine and Serine to Alanine. The former is a change from a nonpolar to a polar amino acid, while the latter is the reverse. These could have an impact on the folding of the protein. Hence, we say that OsTN2t002590.1 is different, making the *OsCOL4* gene of TNG67 and Nipponbare to be similar as supported by their gene structure diagrams and protein sequence alignment. The same argument can be said to the *OsDof12 gene* (see Figure S7c). TN1's OsTN3t000628.1 has an amino change from Alanine to Threonine, and it's change from a nonpolar amino acid to a polar one, making it different to TNG67's OsTNG03t000566.1 and Nipponbare's Os03t0169600-01. In the case of *Hd3a*, there is loss of exonic regions in the middle portion of the gene structure of TNG67 and

TN1 (see Figure S7d). But between the two, TNG67 aligned better against Nipponbare's Os06t0157700-01. So the TNG67 *Hd3a* is more similar to its counterpart in Nipponbare. Meanwhile, looking at the gene structure diagrams for *OsEMF2b* (see Figure S7e), TNG67's OsTNG09t000632.1, Nipponbare's Os09t0306800-01, and TN1's OsTN9t000600.1 have fewer exon losses compared to TN1's OsTN10t002235.1 and OsTN9t000288.1. Checking the alignment would show that TNG67's OsTNG09t000632.1 and Nipponbare's Os09t0306800-01 have more agreement compared to the other orthologues. So we conclude that the OsEMF2b of TNG67 is similar to Nipponbare.

For GF14c, the protein sequence alignment shows an almost perfect agreement among the orthologues, except for the tail-end of the alignment that OsTN8t001598.1 has an amino acid change from Cytosine to Guanine (see Figure S7f). Disregarding the LVK amino acid sequence at the start of the alignment because it don't start with Methionine, we consider the Cysteine to Guanine in OsTN8t001598.1 to be a better identifier, whether what grouping does *GF14c* gene of TNG67 belongs to. Because Cyteine is a polar amino acid, while Glycine is a nonpolar, the change could have an effect in the folding of the protein in TN1. Cysteine could form a disulfide bond with another Cysteine, and the change would mean a loss of this interaction in the protein structure of TN1 GF14c. With the OsTNG08t001966.1 having a very good alignment with the three isoforms of Nipponbare, we say that the TNG67 GF14c is similar to Nipponbare.

*a) Hd16*

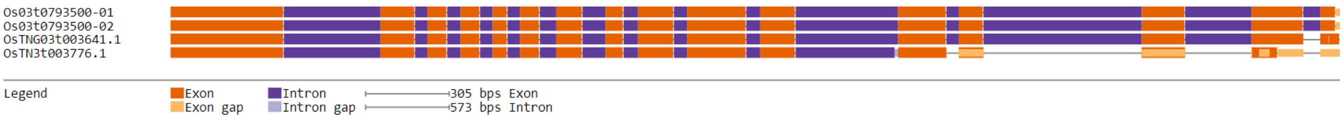

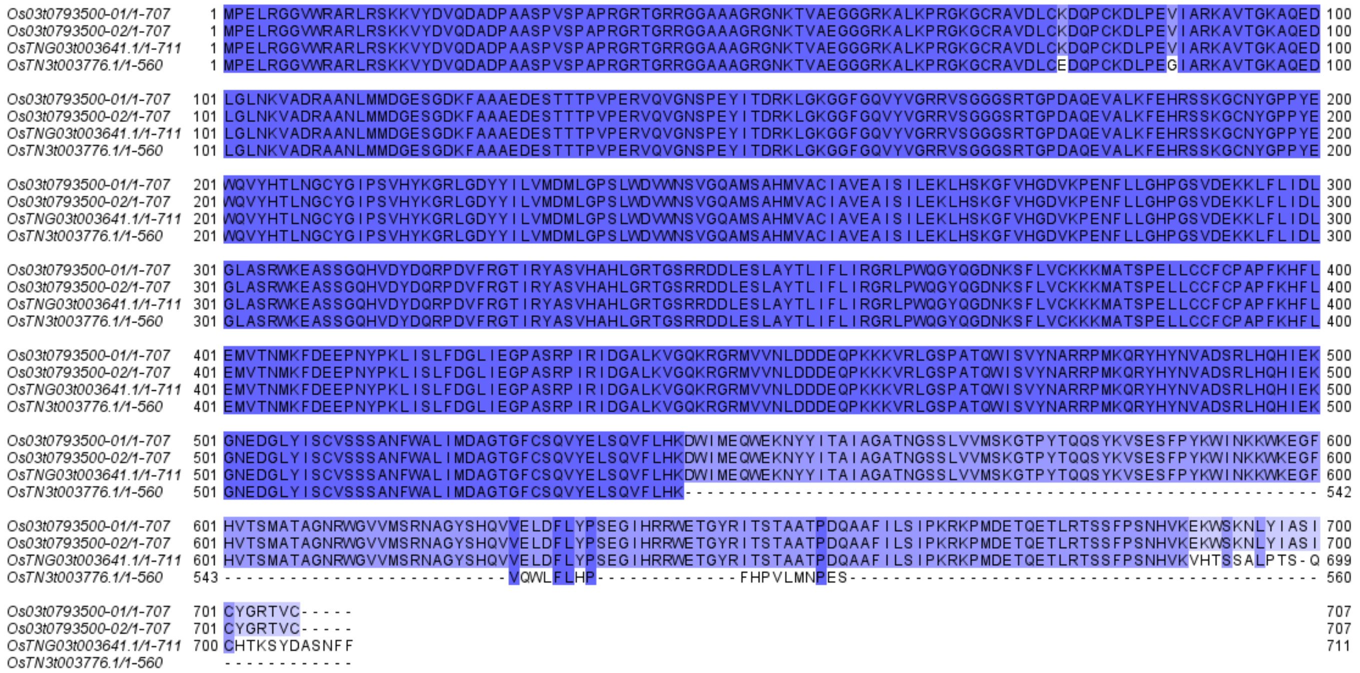


*b) OsCOL4*

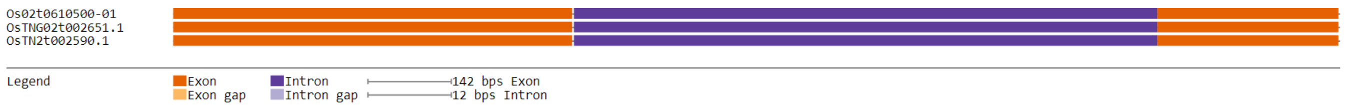


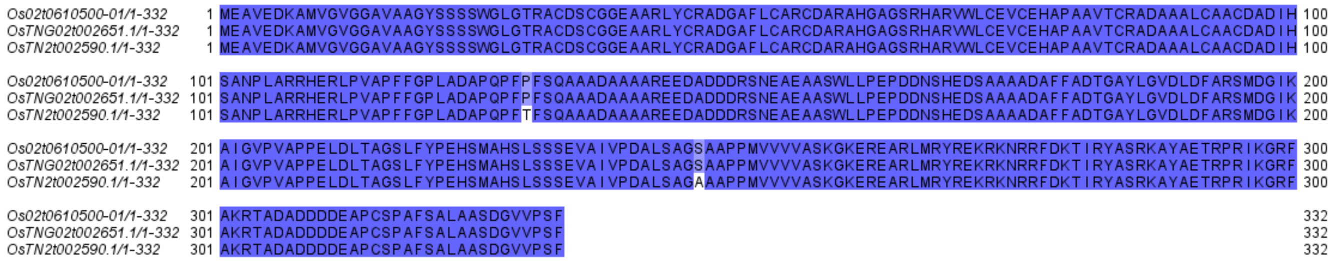


*c) OsDof12*

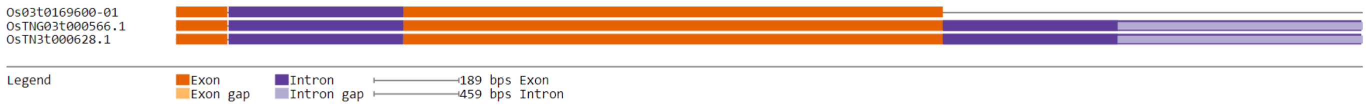

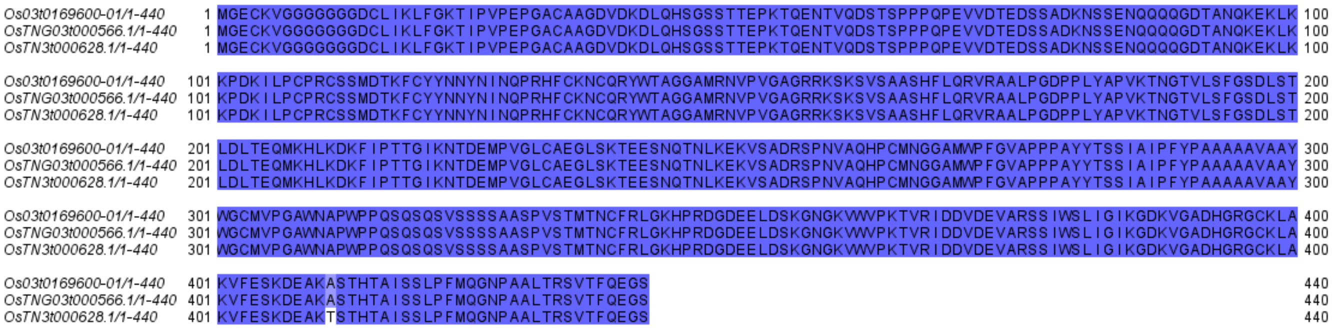


*d) Hd3a*

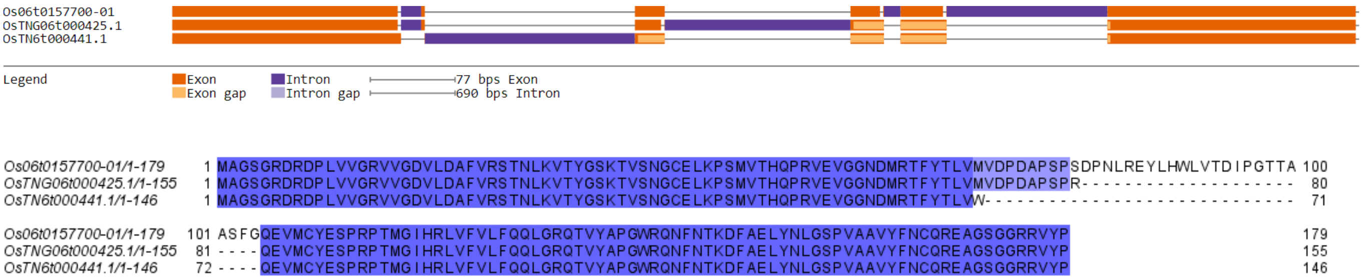


*e) OsEMF2b*

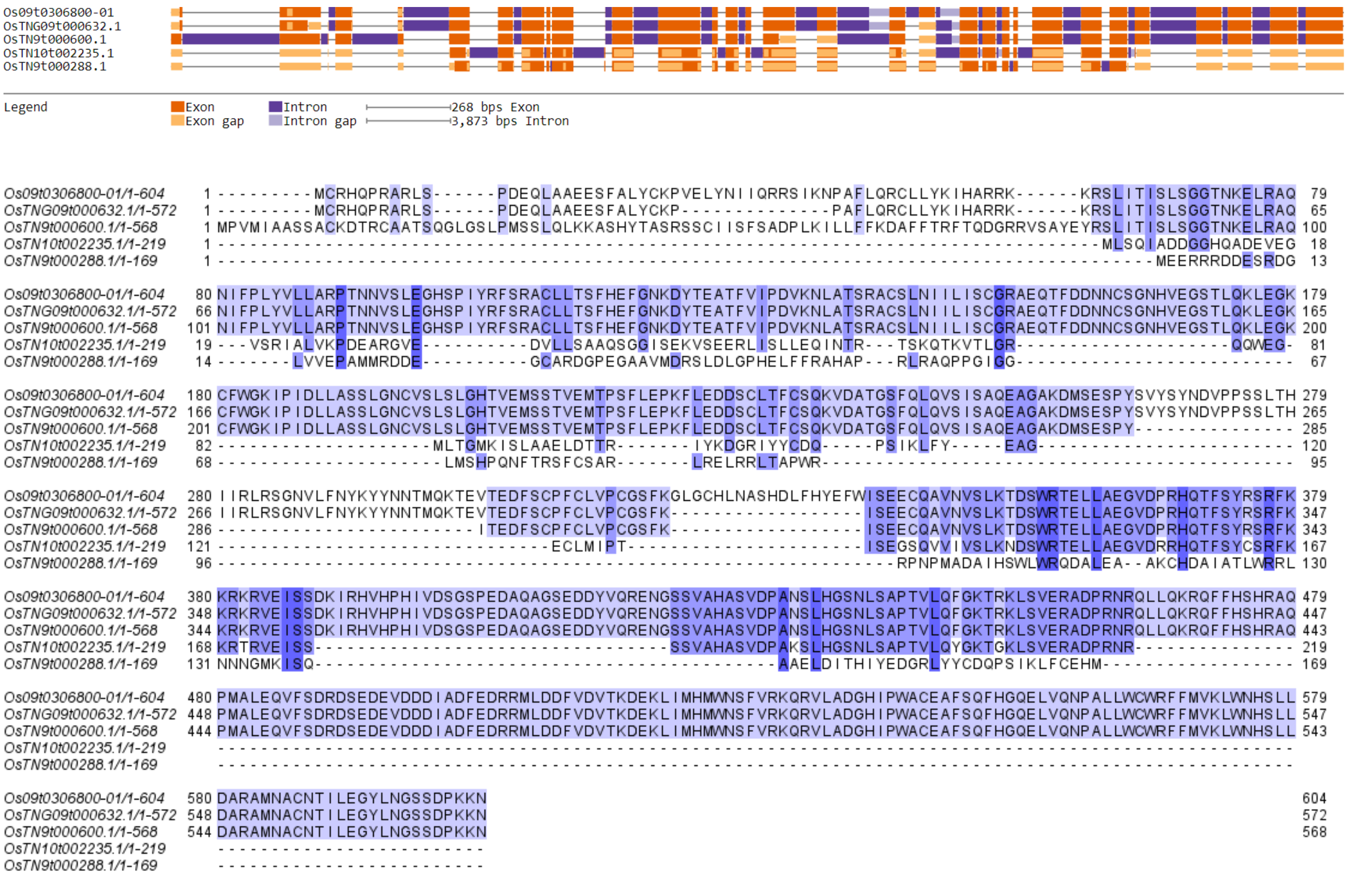


*f) GF14c*

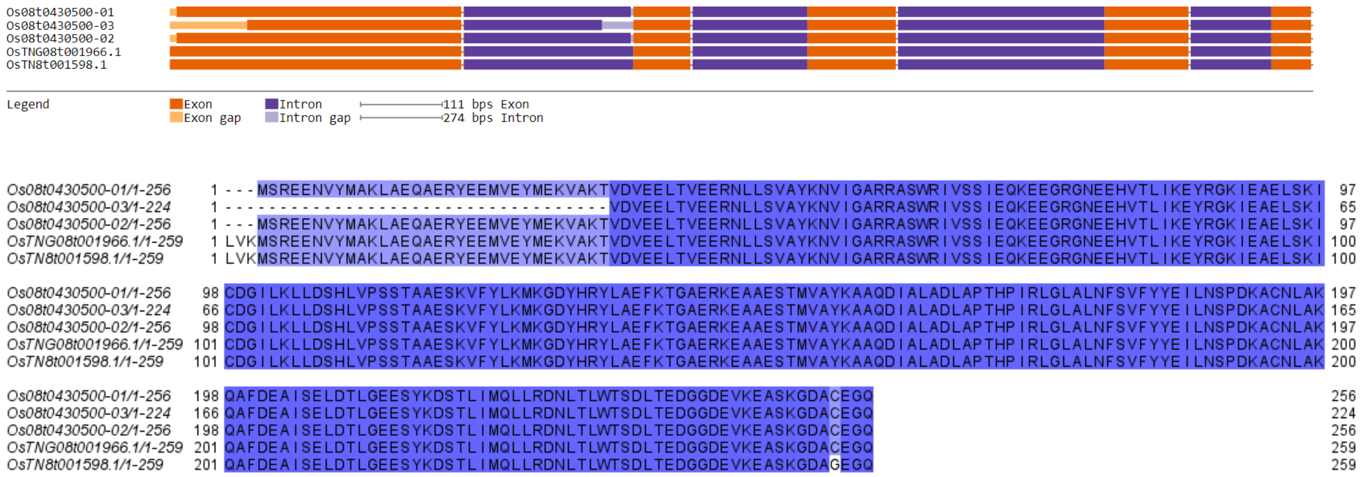


**Figure S7. Gene structure diagrams and protein sequence alignment of photoperiod genes of TNG67 that are similar to Nipponbare.** a) *Hd16*, b) *OsCOL4*, c) *OsDof12*, d) *Hd3a*, e) *OsEMF2b*, and f) *GF14c*

#### *TNG67 is unique*

We found three photoperiod genes of TNG67 that are unique when compared to its equivalents in Nipponbare and TN1. They are *PHYA*, *EHD3* and *OsCO3*. These unique genes contain an amino acid sequence or an exon/exonic region in the gene that is only found in TNG67. We analyzed the *PHYA* (see Figure S8a) and *EHD3* (see Figure S8b), and their proteins have longer lengths, 1,208 aa for OsTNG03t003134.1 and 670 aa for OsTNG08t000038.1. These PHYA and EHD3 proteins of TNG67 have a different sequence for their N-terminal, making them longer than their orthologues. We expect the structure of these PHYA and EHD3 proteins to be different compared to their orthologues in Nipponbare and TN1. For the case of the *OsCO3* gene of TNG67, its protein sequence is the same for the first 208 aa when compared to its orthologues in Nipponbare and TN1. After that, the sequences of the three cultivars differ with each other (see Figure S8c). Through the sequence alignment, the TNG67 OsCO3 is unique with respect to its equivalent proteins. For OsPIPK1, the we looked at the uncolored sequences in the alignment because these are the unaligned sequences. A large portion of these unmatched sequences belongs to TNG67's OsTNG03t003013.1, so we say that the OsPIPK1 of TNG67 is unique. For *Oscry2*, TNG67's unique sequence is found on the its 1$^{st}$ exon, 2$^{nd}$ exon, and a small portion of its 3rd exon. There is also another unique sequence found at the 7th exon of the gene (see Figure S8e). These unique sequences can be easily spotted as the unaligned amino acids in the protein sequence alignment (see Figure S8e).

*a) PHYA*

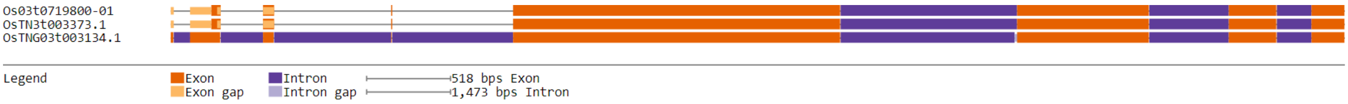


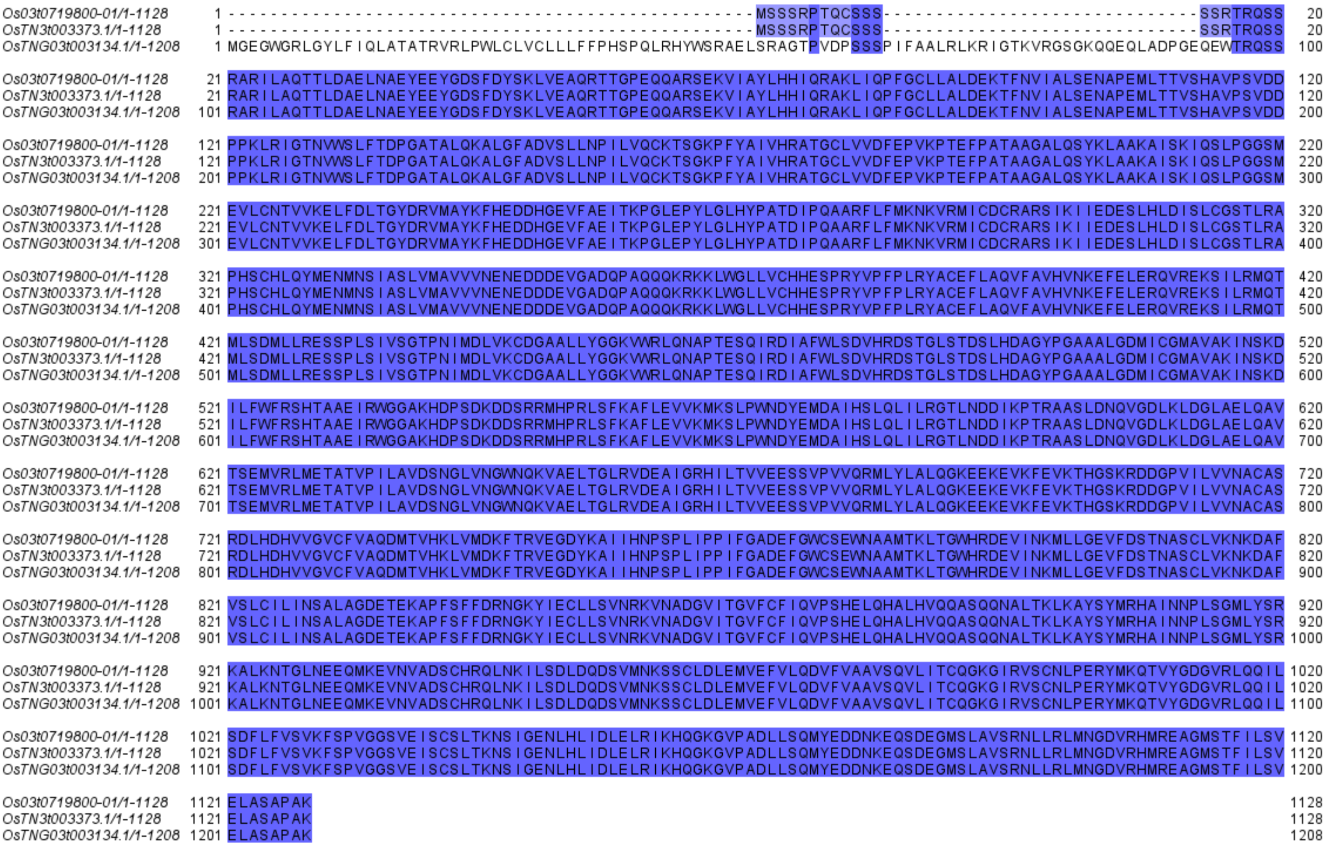


*b) EHD3*

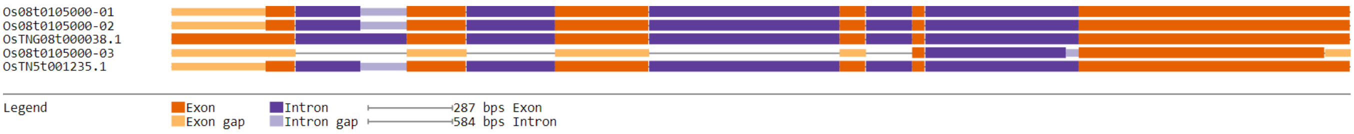

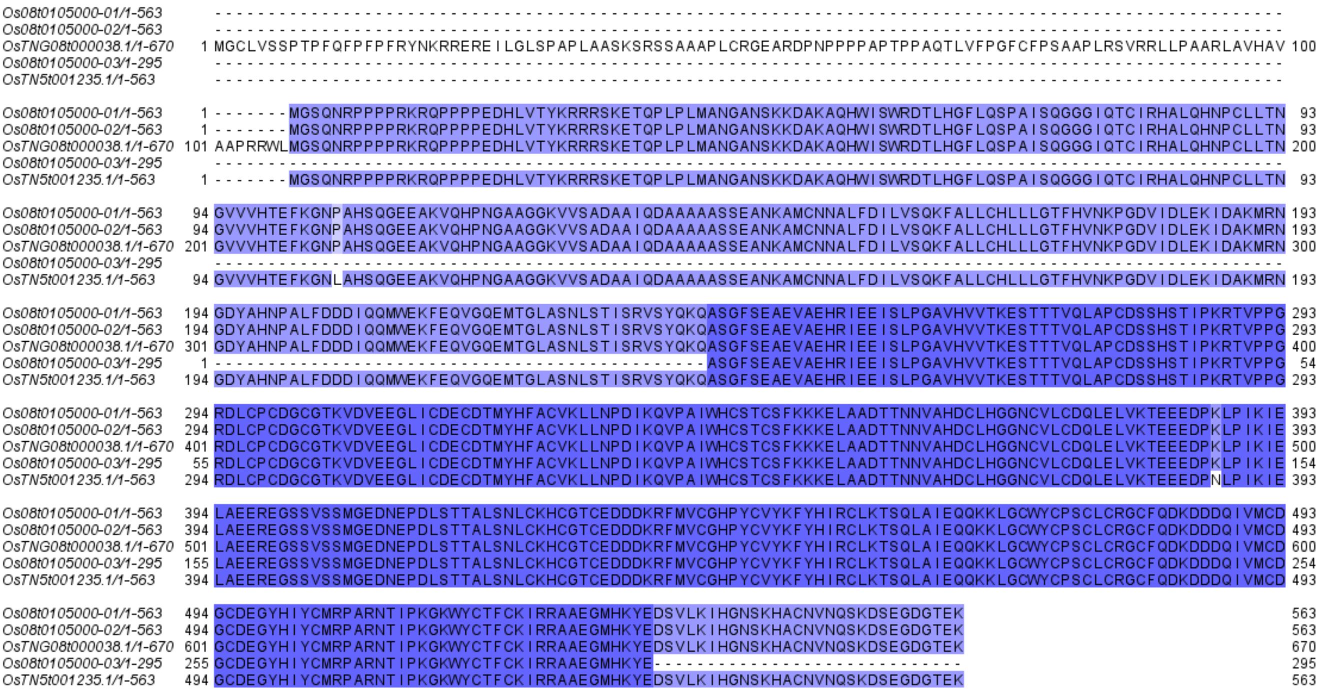


c) *OsCO3*

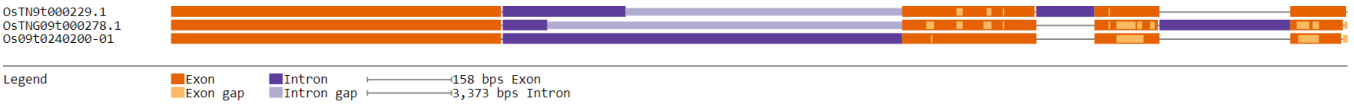


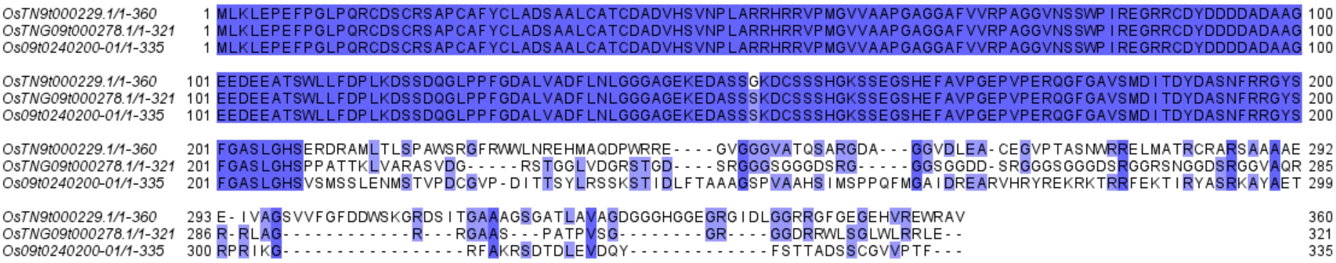


d) *OsPIPK1*

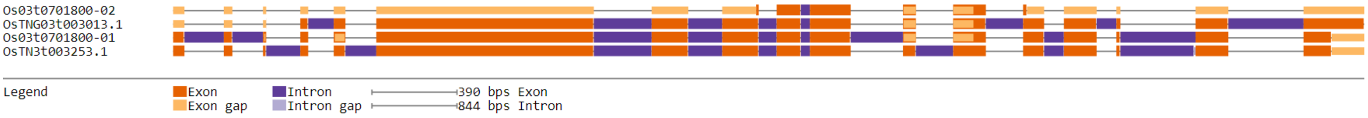

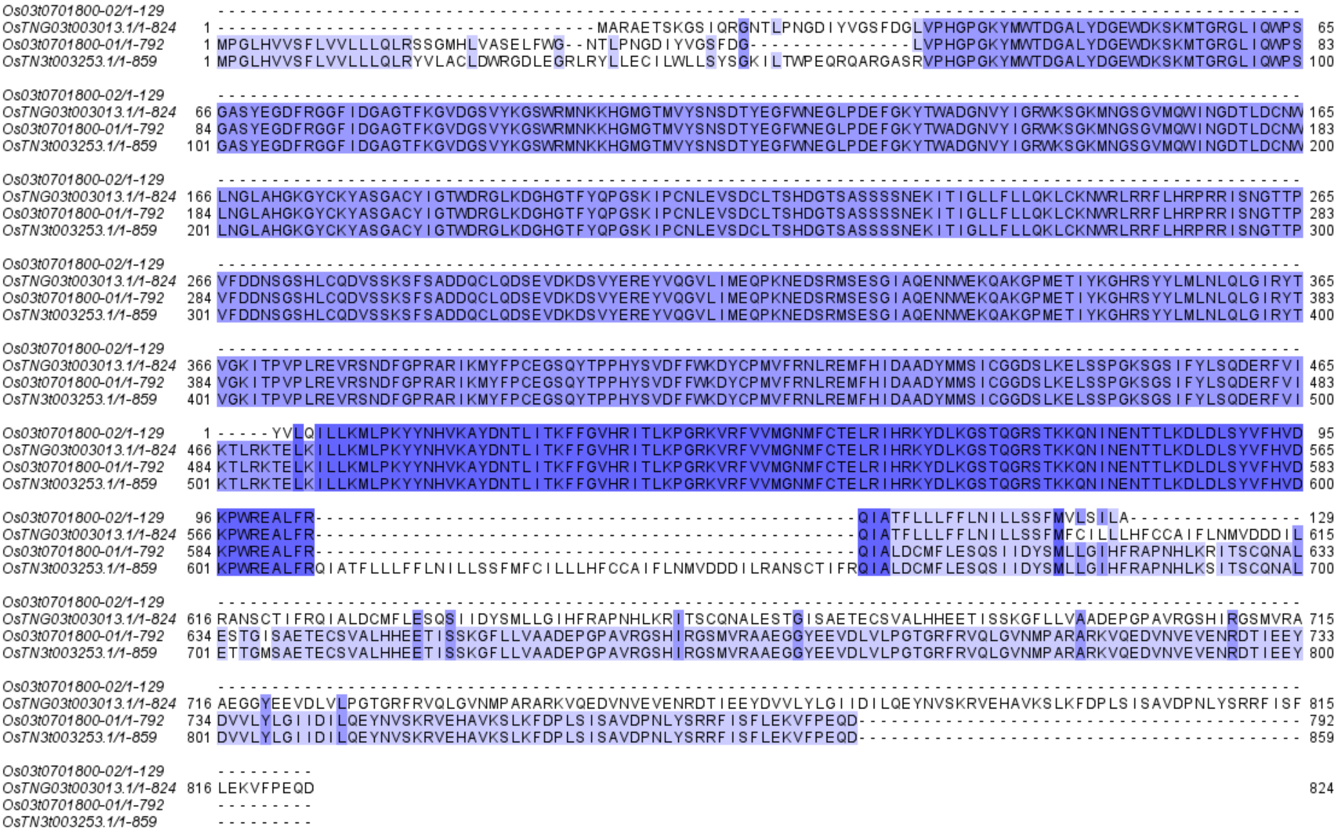

Os03t0701800-02/1-129 ----------------------------------------------------------------------------------------------------
OsTNG03t003013.1/1-824 1 -----------------------------------MARAETSKGSIQRGNTLPNGDIYVGSFDGLVPHGPGKYMWTDGALYDGEWDKSKMTGRGLIQWPS 65
Os03t0701800-01/1-792 1 MPGLHVVSFLVVLLLQLRSSGMHLVASELFWG--NTLPNGDIYVGSFDG---------------LVPHGPGKYMWTDGALYDGEWDKSKMTGRGLIQWPS 83
OsTN3t003253.1/1-859 1 MPGLHVVSFLVVLLLQLRYVLACLDWRGDLEGRLRYLLECILWLLSYSGKILTWPEQRQARGASRVPHGPGKYMWTDGALYDGEWDKSKMTGRGLIQWPS 100
Os03t0701800-02/1-129 ----------------------------------------------------------------------------------------------------
OsTNG03t003013.1/1-824 66 GASYEGDFRGGFIDGAGTFKGVDGSVYKGSWRMNKKHGMGTMVYSNSDTYEGFWNEGLPDEFGKYTWADGNVYIGRWKSGKMNGSGVMQWINGDTLDCNW 165
Os03t0701800-01/1-792 84 GASYEGDFRGGFIDGAGTFKGVDGSVYKGSWRMNKKHGMGTMVYSNSDTYEGFWNEGLPDEFGKYTWADGNVYIGRWKSGKMNGSGVMQWINGDTLDCNW 183
OsTN3t003253.1/1-859 101 GASYEGDFRGGFIDGAGTFKGVDGSVYKGSWRMNKKHGMGTMVYSNSDTYEGFWNEGLPDEFGKYTWADGNVYIGRWKSGKMNGSGVMQWINGDTLDCNW 200
Os03t0701800-02/1-129 ----------------------------------------------------------------------------------------------------
OsTNG03t003013.1/1-824 166 LNGLAHGKGYCKYASGACYIGTWDRGLKDGHGTFYQPGSKIPCNLEVSDCLTSHDGTSASSSSNEKITIGLLFLLQKLCKNWRLRRFLHRPRRISNGTTP 265
Os03t0701800-01/1-792 184 LNGLAHGKGYCKYASGACYIGTWDRGLKDGHGTFYQPGSKIPCNLEVSDCLTSHDGTSASSSSNEKITIGLLFLLQKLCKNWRLRRFLHRPRRISNGTTP 283
OsTN3t003253.1/1-859 201 LNGLAHGKGYCKYASGACYIGTWDRGLKDGHGTFYQPGSKIPCNLEVSDCLTSHDGTSASSSSNEKITIGLLFLLQKLCKNWRLRRFLHRPRRISNGTTP 300
Os03t0701800-02/1-129 ----------------------------------------------------------------------------------------------------
OsTNG03t003013.1/1-824 266 VFDDNSGSHLCQDVSSKSFSADDQCLQDSEVDKDSVYEREYVQGVLIMEQPKNEDSRMSESGIAQENNWEKQAKGPMETIYKGHRSYYLMLNLQLGIRYT 365
Os03t0701800-01/1-792 284 VFDDNSGSHLCQDVSSKSFSADDQCLQDSEVDKDSVYEREYVQGVLIMEQPKNEDSRMSESGIAQENNWEKQAKGPMETIYKGHRSYYLMLNLQLGIRYT 383
OsTN3t003253.1/1-859 301 VFDDNSGSHLCQDVSSKSFSADDQCLQDSEVDKDSVYEREYVQGVLIMEQPKNEDSRMSESGIAQENNWEKQAKGPMETIYKGHRSYYLMLNLQLGIRYT 400
Os03t0701800-02/1-129 ----------------------------------------------------------------------------------------------------
OsTNG03t003013.1/1-824 366 VGKITPVPLREVRSNDFGPRARIKMYFPCEGSQYTPPHYSVDFFWKDYCPMVFRNLREMFHIDAADYMMSICGGDSLKELSSPGKSGSIFYLSQDERFVI 465
Os03t0701800-01/1-792 384 VGKITPVPLREVRSNDFGPRARIKMYFPCEGSQYTPPHYSVDFFWKDYCPMVFRNLREMFHIDAADYMMSICGGDSLKELSSPGKSGSIFYLSQDERFVI 483
OsTN3t003253.1/1-859 401 VGKITPVPLREVRSNDFGPRARIKMYFPCEGSQYTPPHYSVDFFWKDYCPMVFRNLREMFHIDAADYMMSICGGDSLKELSSPGKSGSIFYLSQDERFVI 500
Os03t0701800-02/1-129 1 -----YVLQILLKMLPKYYNHVKAYDNTLITKFFGVHRITLKPGRKVRFVVMGNMFCTELRIHRKYDLKGSTQGRSTKKQNINENTTLKDLDLSYVFHVD 95
OsTNG03t003013.1/1-824 466 KTLRKTELKILLKMLPKYYNHVKAYDNTLITKFFGVHRITLKPGRKVRFVVMGNMFCTELRIHRKYDLKGSTQGRSTKKQNINENTTLKDLDLSYVFHVD 565
Os03t0701800-01/1-792 484 KTLRKTELKILLKMLPKYYNHVKAYDNTLITKFFGVHRITLKPGRKVRFVVMGNMFCTELRIHRKYDLKGSTQGRSTKKQNINENTTLKDLDLSYVFHVD 583
OsTN3t003253.1/1-859 501 KTLRKTELKILLKMLPKYYNHVKAYDNTLITKFFGVHRITLKPGRKVRFVVMGNMFCTELRIHRKYDLKGSTQGRSTKKQNINENTTLKDLDLSYVFHVD 600
Os03t0701800-02/1-129 96 KPWREALFR--------------------------------------------------QIATFLLLFFLNILLSSFMVLSILA---------------- 129
OsTNG03t003013.1/1-824 566 KPWREALFR--------------------------------------------------QIATFLLLFFLNILLSSFMFCILLLHFCCAIFLNMVDDDIL 615
Os03t0701800-01/1-792 584 KPWREALFR--------------------------------------------------QIALDCMFLESQSIIDYSMLLGIHFRAPNHLKRITSCQNAL 633
OsTN3t003253.1/1-859 601 KPWREALFRQIATFLLLFFLNILLSSFMFCILLLHFCCAIFLNMVDDDILRANSCTIFRQIALDCMFLESQSIIDYSMLLGIHFRAPNHLKSITSCQNAL 700
Os03t0701800-02/1-129 ----------------------------------------------------------------------------------------------------
OsTNG03t003013.1/1-824 616 RANSCTIFRQIALDCMFLESQSIIDYSMLLGIHFRAPNHLKRITSCQNALESTGISAETECSVALHHEETISSKGFLLVAADEPGPAVRGSHIRGSMVRA 715
Os03t0701800-01/1-792 634 ESTGISAETECSVALHHEETISSKGFLLVAADEPGPAVRGSHIRGSMVRAAEGGYEEVDLVLPGTGRFRVQLGVNMPARARKVQEDVNVEVENRDTIEEY 733
OsTN3t003253.1/1-859 701 ETTGMSAETECSVALHHEETISSKGFLLVAADEPGPAVRGSHIRGSMVRAAEGGYEEVDLVLPGTGRFRVQLGVNMPARARKVQEDVNVEVENRDTIEEY 800
Os03t0701800-02/1-129 ----------------------------------------------------------------------------------------------------
OsTNG03t003013.1/1-824 716 AEGGYEEVDLVLPGTGRFRVQLGVNMPARARKVQEDVNVEVENRDTIEEYDVVLYLGIIDILQEYNVSKRVEHAVKSLKFDPLSISAVDPNLYSRRFISF 815
Os03t0701800-01/1-792 734 DVVLYLGIIDILQEYNVSKRVEHAVKSLKFDPLSISAVDPNLYSRRFISFLEKVFPEQD----------------------------------------- 792
OsTN3t003253.1/1-859 801 DVVLYLGIIDILQEYNVSKRVEHAVKSLKFDPLSISAVDPNLYSRRFISFLEKVFPEQD----------------------------------------- 859
Os03t0701800-02/1-129 ---------
OsTNG03t003013.1/1-824 816 LEKVFPEQD 824
Os03t0701800-01/1-792 ---------
OsTN3t003253.1/1-859 ---------


*e) Oscry2*

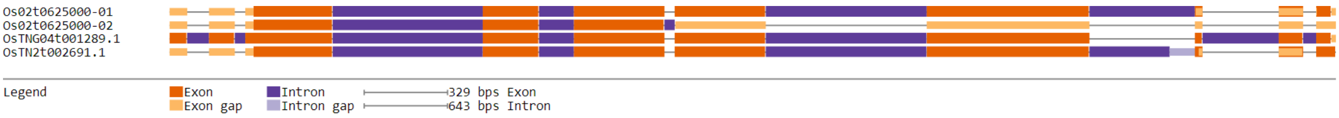

Os02t0625000-01
Os02t0625000-02
OsTNG04t001289.1
OsTN2t002691.1
Legend
Exon
Exon gap
Intron
Intron gap
329 bps Exon
643 bps Intron

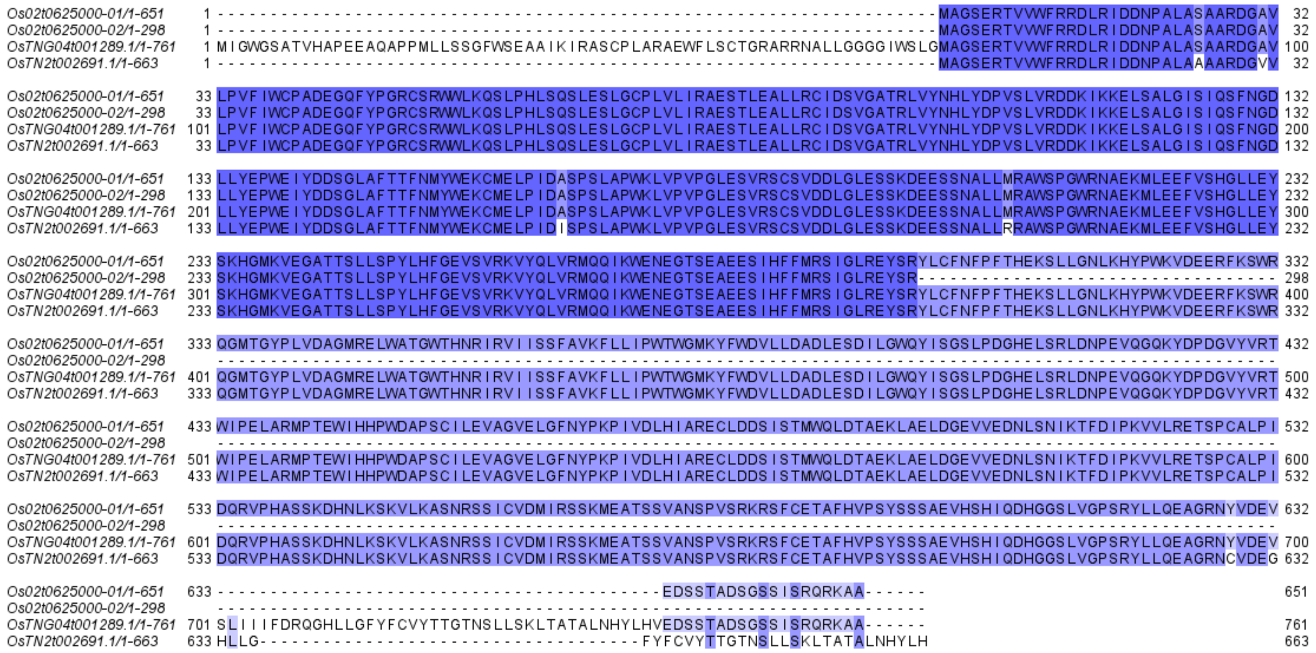

**Figure S8. Gene structure diagrams and protein sequence alignment of photoperiod genes of TNG67 that are unique**. a) *PHYA*, b) *EHD3*, c) *OsCO3*, d) *OsPIPK1*, e) *Oscry2*

***TNG67 is hybrid***

Photoperiod hybrid genes of TNG67 contain segments in its gene that are similar to both Nipponbare and TN1. To detect whether a photoperiod gene of TNG67 is a hybrid or not, we check first their gene structure diagrams for segments that are similar to Nipponbare, and others that are similar to TN1. When none is found, we checked the protein sequence alignment and looked for gaps or amino acid changes that would indicate the similarities or differences between the cultivars. We found eight photoperiod genes of TNG67 that are considered to be hybrid. These are *OsELF3-1/EF7*, *OsHDT1*, *OsATG7*, *DTH8*, *Hd1*, *EF1*, *Ghd8*, and *Ehd1*. For OsELF3-1, Nipponbare has lost an exonic region in exon 1 (see Figure S9a), though the sequence alignment shows that TNG7's OsTNG06t000314.1 and TN1's OsTN6t000324.1 aligned very well, their first amino acid is Leucine, which could be the result of an error in the annotation. For the amino acid substitutions in the alignment, they favor TNG67 being similar to Nipponbare (see Figure S9a). We also used amino acid substitutions in the alignment (see Figure S9b) in determining the TNG67 *OsATG7* as a hybrid gene. The same can be said about

EF1 with a CWFNF sequence only found between TNG67 and Nipponbare (see Figure S9c). OsHDT1 is a special case because it is a combination of a unique sequence, not found in either Nipponbare or TN1, plus segments that are similar with Nipponbare. The unique sequence of TNG67 OsHDT1 can be easily seen as the uncolored sequence in the alignment, while the shared sequence between TNG67 and Nipponbare is shaded in violet blue (see Figure S9d).

For DTH8, it has a high similarity with Nipponbare, but at the end of the gene it becomes similar to TN1. Because Nipponbare has two isoforms in Os08t0174500-01 and Os08t0174500-02, we only chose the former as the representative protein of Nipponbare DTH8. The sequence alignment shows that there is high similarity between TNG67's OsTNG08t000651.1 and two isoforms of Nipponbare. TN1's OsTN8t000434.1 had amino acid changes making it different with its orthologue in TNG67. But the DTH8 proteins of TNG67 (351 aa) and TN1 (343 aa) are longer than their counterparts in Nipponbare, so at the end of the alignment, TNG67 and TN1 becomes similar. For *Hd1*, the loss of exonic regions near the end and start of the gene structure (see Figure S9f) is where the state of being hybrid of the TNG67 gene was defined. In the protein sequence alignment, the gap in Nipponbare's Os06t0275000-01 (see Figure S9f), made TNG67's OsTNG06t001241.1 and TN1's OsTN6t001222.1 to be similar. There are also single amino acid changes wherein TNG67 and Nipponbare are the same, and TN1 is different, such as Asparagine to Lysine and Arginine to Glutamine (see Figure S9f). While near the end of alignment, the sequence of TN1's OsTN6t001222.1 has become different when compared to its orthologues, which shares their sequence with each other (see Figure S9f). Overall, *Hd1* is a hybrid photoperiod gene of TNG67. In *Ghd8*, the hybrid nature of can be easily identified through the exonic loss on its gene structure (see Figure S9g). The exons were divided into four, with the yellow bars at the tail-end representing the C-terminal portion alignment, wherein TNG67's OsTNG08t000651.1

and TN1's OsTN8t000434.1 are similar. In contrast, the Ghd8 TNG67 and Nipponbare isoforms agree to each other and these are colored violet-blue in the alignment (see Figure 8g). There are reasons why *OsTrx1* is a hybrid photoperiod gene. We first disregard the aligned sections of the TNG67's OsTNG09t000260.1 and TN1's OsTN9t000211.1 at the start of the protein sequence alignment. Those aligned sequences could be the result of an error in the annotation, as the alignment does not start with Methionine. For the missing 3$^{rd}$ exon in Nipponbare (see Figure S9h), they correspond to an alignment between TNG67 and TN1 only, which makes the OsTrx1 of TNG67 to be similar to TN1. For evidences supporting the TNG67 gene to be similar to Nipponbare are the amino acid change from Proline (nonpolar amino acid) to Glutamine (polar amino acid) and the VN subsequence in TNG67's OsTNG09t000260.1 and Nipponbare's Os09t0134500-02 vs RR subsequence in TN1's OsTN9t000211.1 (see Figure S9h). For PHYB, the protein sequence alignment shows why the gene is a hybrid photoperiod TNG67 gene. The amino acid changes that are violet-blue colored in Figure S9i, illustrates the similarity between TNG67 and Nipponbare. Meanwhile, the exonic loss in the Nipponbare PHYB corresponds to a gap in the protein sequence alignment (see Figure S9i), wherein TNG67's OsTNG02t001958.1 and TN1's OsTN3t001707.1 aligned perfectly. Thus, we say that the *PHYB* gene of TNG67 belongs to the hybrid group of photoperiod genes.

The *NRRa/NRRb* gene is another example of a hybrid photoperiod gene. It contains a unique sequence corresponding to the unaligned region for OsTNG05t003013.1 in the protein sequence alignment (see Figure S9j). In addition, the tail-end of the alignment shows an agreement of the sequences between TNG67's OsTNG05t003013.1 and two isoforms of TN1 (see Figure S9j), suggesting a similarity to the latter. With a unique sequence plus a TNG67/TN1 similarity, we conclude that the *NRRa/NRRb* gene of TNG67 is a hybrid photoperiod gene.

### *a) OsELF3-1/EF7*

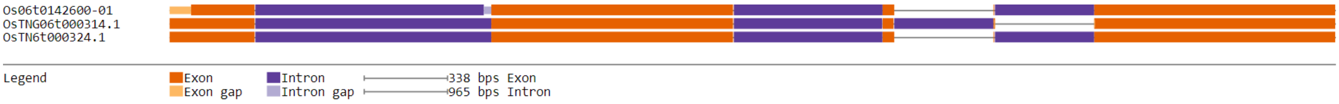


```
Os06t0142600-01/1-760      1 -----------------------------MATRGGGGGGGGKEAKGKVMGPLFPRLHVNDAAKGGGPRAPPRNKMALYEQFTVPSHRFSGGGGGGGVGGS  71
OsTNG06t000314.1/1-787     1 LPFFWPQKPATSSTPPWPDAPEKLLRREAMATRGGGGGGGGKEAKGKVMGPLFPRLHVNDAAKGGGPRAPPRNKMALYEQFTVPSHRFSGGGGGGGVGGS 100
OsTN6t000324.1/1-789       1 LPFFWPQKPATSSTPPWPDAPEKLLRREAMATRGGGGGGGGKEAKGKVMGPLFPRLHVNDAAKGGGPRAPPRNKMALYEQFTVPSHRFSGGGGGGGVGGS 100

Os06t0142600-01/1-760     72 PAHSTSAASQSQSQSQVYGRDSSLFQPFNVPSNRPGHSTEKINSDKINKKISGSRKELGMLSSQTKGMDIYASRSTAEAPQRRAENTIKSSSGKRLADDD 171
OsTNG06t000314.1/1-787   101 PAHSTSAASQSQSQSQVYGRDSSLFQPFNVPSNRPGHSTEKINSDKINKKISGSRKELGMLSSQTKGMDIYASRSTAEAPQRRAENTIKSSSGKRLADDD 200
OsTN6t000324.1/1-789     101 PAHSTSAASQSQSQSQVYGRDSSLFQPFNVPSNRPGHSTEKINSDKINKKISGSRKELGMLSSQTKGMDIYASRSTAEAPQRRAENTIKSSSGKRLADDD 200

Os06t0142600-01/1-760    172 EFMVPSVFNSRFPQYSTQENAGVQDQSTPLVAANPHKSPSTVSKSSTKCYNTVSKKLERIHVSDVKSRTPLKDKEMEAAQTSKNVEVEKSSSFHASKDMF 271
OsTNG06t000314.1/1-787   201 EFMVPSVFNSRFPQYSTQENAGVQDQSTPLVAANPHKSPSTVSKSSTKCYNTVSKKLERIHVSDVKSRTPLKDKEMEAAQTSKNVEVEKSSSFHASKDMF 300
OsTN6t000324.1/1-789     201 EFMVPSVFNSRFPQYSTQENAGVQDQSTPLVAANPHKSPSTVSKSSTKCYNTVSKKLERIHVSDVKSRTPLKDKEMEAAQTSKNVEVEKSSSFHASKDMF 300

Os06t0142600-01/1-760    272 ESRHAKVYPKMDKTGIINDSDEPHGGNSGHQATSRNGGSMKFQNPPMRRNEISSNPSSENTDRHYNLPQGGIEETGTKRKRLLEQHDAEKSDDVSRLLEQ 371
OsTNG06t000314.1/1-787   301 ESRHAKVYPKMDKTGIINDSDEPHGGNSGHQATSRNGGSMKFQNPPMRRNEISSNPSSENTDRHYNLPQGGIEETGTKRKRLLEQHDAEKSDDVSRLLEQ 400
OsTN6t000324.1/1-789     301 ESRHAKVYPKMDKTGIINDSDEPHGGNSGHQATSRNGGSMKFQNPPMRRNEISSNPSSENTDRHYNLLQGGIEETGTKRKRLLEQHDAEKSDDVSRLLEQ 400

Os06t0142600-01/1-760    372 HDAENIDDVSDSSVECITGWEISPDKIVGAIGTKHFWKARRAIMNQQRVFAVQVFELHKLVKVQKLIAASPHVLIESDPCLGNALLGSKNKLVEENLKAQ 471
OsTNG06t000314.1/1-787   401 HDAENIDDVSDSSVECITGWEISPDKIVGAIGTKHFWKARRAIMNQQRVFAVQVFELHKLV--QKLIAASPHVLIESDPCLGNALLGSKNKLVEENLKAQ 498
OsTN6t000324.1/1-789     401 HDAENIDDVSDSSVECITGWEISPDKIVGAIGTKHFWKARRAIMNQQRVFAVQVFELHKLVKVQKLIAASPHVLIEGDPCLGNALLGSKNKLVEENLKAQ 500

Os06t0142600-01/1-760    472 PLLVATIDDVEPSLQQPEVSKENTEDSPPSPHDTGLGSGQRDQAATNGVSKSNRRATPVASDNKQNNWGVQLQPPQNQWLVPVMSPLEGLVYKPYSGPCP 571
OsTNG06t000314.1/1-787   499 PLLVATIDDVEPSLQQPEVSKENTEDSPPSPHDTGLGSGQRDQAATNGVSKSNRRATPVASDNKQNNWGVQLQPPQNQWLVPVMSPLEGLVYKPYSGPCP 598
OsTN6t000324.1/1-789     501 PLLVATIDDVEPSLQQPEVSKENTEDSPPSPHDTGLGSGQRDQAATNGVSKSNRRATPVASDNKQNNWGVQLQPPQNQWLVPVMSPSEGLVYKPYSGPCP 600

Os06t0142600-01/1-760    572 PAGSILAPFYANCTPLSLPSTAGDFMNSAYGVPMPHQPQHMGAPGPPSMPMNYFPPFSIPVMNPTAPAPVVEQGRHPSMPQPYGNFEQQSWISCNMSHPS 671
OsTNG06t000314.1/1-787   599 PAGSILAPFYANCTPLSLPSTAGDFMNSAYGVPMPHQPQHMGAPGPPSMPMNYFPPFSIPVMNPTAPAPVVEQGRHPSMPQPYGNFEQQSWISCNMSHPS 698
OsTN6t000324.1/1-789     601 PAGSILAPFYANCTPLSLPSTAGDFMNSAYGVPMPHQPQHMGAPGPPSMPMNYFPPFSIPVMNPTAPAPVVEQGRHPSMPQPYGNFEQQSWISCNMSHPS 700

Os06t0142600-01/1-760    672 GIWRFHASRDSEAQASSASSPFDRFQCSGSGPVSAFPTVSAQNNQPQPSYSSRDNQTNVIKVVPHNSRTASESAARIFRSIQMERQRDD            760
OsTNG06t000314.1/1-787   699 GIWRFHASRDSEAQASSASSPFDRFQCSGSGPVSAFPTVSAQNNQPQPSYSSRDNQTNVIKVVPHNSRTASESAARIFRSIQMERQRDD            787
OsTN6t000324.1/1-789     701 GIWRFHASRDSEAQASSASSPFDRFQCSGSGPVSAFPTASAQNNQPQPSYGSRDNQTNVIKVVPHNSRTASESAARIFRSIQMERQRDD            789
```

### *b) OsATG7*

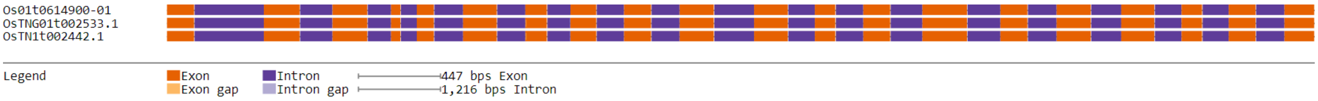


```
Os01t0614900-01/1-1042      1 MAARAEAAAAAPRPLQAAAIGVCAETGFWDALRRLKLDVLGTDDSPIPITGYYTPRQYEKIASLFRICPESILPPSANSFGDRNNCPVPGTLLNTNNMRG  100
OsTNG01t002533.1/1-1041     1 MAARAEAAAAAPRPLQAAAIGVCAETGFWDALRRLKLDVLGTDDSPIPITGYYTPRQYEKIASLFRICPESILPPSANSFGDRNNCPVPGTLLNTNNMRG  100
OsTN1t002442.1/1-1041       1 MAARAEAAAAAPRPLQAAAIGVCAETGFWDALRRLKLDVLGTDDSPIPITGYYTPRQYEKIASLFRICPESILPPSANSFGDRNNCPVPGTLLNTNNMRG  100

Os01t0614900-01/1-1042    101 FQNLDRALLLKAEAKKILHDIKSGKVEENPALLLRFLVISFADLKNWKVYYNVAFPSLIFDSKITLLSLKLASQVLKQEEATSLSNAFTEWRKSSETTVV  200
OsTNG01t002533.1/1-1041   101 FQNLDRALLLKAEAKKILHDIKSGKVEENPALLLRFLVISFADLKNWKVYYNVAFPSLIFDSKITLLSLKLASQVLKQEEATSLSNAFTEWRKSSETTVV  200
OsTN1t002442.1/1-1041     101 FQNLDRALLLKAEAKKILHDIKSGKVEENPALLLRFLVISFADLKNWKVYYNVAFPSLIFDSKITLLSLKLASQVLKQEEATSLSNAFTEWRKSSETTVV  200

Os01t0614900-01/1-1042    201 PFFLINISPDSSATIRQLKDWKACQGNGQKLLFGFYDHGNRGFPGWALRNYIAFVSLRWKIEKVHFFCYREKRGRPDIQQSLVGEASFPAPHAGWDEPDY  300
OsTNG01t002533.1/1-1041   201 PFFLINISPDSSATIRQLKDWKACQGNGQKLLFGFYDHGNRGFPGWALRNYIAFVSLRWKIEKVHFFCYREKRGRPDIQQSLVGEASFPAPHG-WDEPDY  299
OsTN1t002442.1/1-1041     201 PFFLINISPDSSATIRQLKDWKACQGNGQKLLFGFYDHGNRGFPGWALRNYIAFVSLRWKIEKVHFFCYREKRGRPDIQQSLVGEASFPAPHG-WDEPDY  299

Os01t0614900-01/1-1042    301 VPEAIGWEGETAGKESKEMKPKEIDLSSINPASQDEEKQLMHLKLMGWRHFPVNLDKLAGVRCLLLGAGTLGCEVARLLMTWGVRKLTVVDDGCVSMSDL  400
OsTNG01t002533.1/1-1041   300 VPEAIGWEGETAGKESKEMKPKEIDLSSINPASQDEEKQLMHLKLMGWRHFPVNLDKLAGVRCLLLGAGTLGCEVARLLMTWGVRKLTVVDDGCVSMSDL  399
OsTN1t002442.1/1-1041     300 VPEAIGWEGETAGKESKEMKPKEIDLSSINPASQDEEKQLMHLKLMGWRHFPVNLDKLAGVRCLLLGAGTLGCEVARLLMTWGVRKLTVVDDGCVSMSDL  399

Os01t0614900-01/1-1042    401 VKQSLYTDKDCGVPRVTAIVPHLKERCSAVEVEGIQMGIPKLEYNISASKISSITDDCKRLQTLVDSNDVVFLLNETWEGMWLPTLLCADKNKIAITVLL  500
OsTNG01t002533.1/1-1041   400 VKQSLYTDKDCGVPRVTAIVPHLKERCSAVEVEGIQMGIPKLEYNISASKISSITDDCKRLQTLVDSNDVVFLLNETWEGMWLPTLLCADKNKIAITVLL  499
OsTN1t002442.1/1-1041     400 VKQSLYTDKDCGVPRVTAIVPHLKERCSAVEVEGIQMGIPKLEYNISASKISSITDDCKRLQTLVDSNDVVFLLNETWEGMWLPTLLCADKNKIAITVLL  499

Os01t0614900-01/1-1042    501 GYDNYLVMRHGAGPGTKSGGMDEGIAQIENLSTQDALGRQRLGCCFCSDTTSLVNSDHNGALDQQSAVILPGLTSVASGKAVELFARMLHHPDEIHAPGD  600
OsTNG01t002533.1/1-1041   500 GYDNYLVMRHGAGPGTKSGGMDEGIAQIENLSTQDALGRQRLGCCFCSDTTSLVNSDHNGALDQQSAVILPGLTSVASGKAVELFARMLHHPDEIHAPGD  599
OsTN1t002442.1/1-1041     500 GYDNYLVMRHGAGPGTKSGGMDEGIAQIENLSTQDALGRQRLGCCFCSDTTSLVNSDHNGALDQQSAVILPGLTSVASGKAVELFARMLHHPDEIHAPGD  599

Os01t0614900-01/1-1042    601 IAGTDTEHQLGLLPHQMQGSLSKCVLSTVLCNSSSNCIACSNAVLSEYRRRGFDFVTQAITCPTYLKDLTGISDLKKPFASKISASIPVSKTSASIPVNL  700
OsTNG01t002533.1/1-1041   600 IAGTDTEHQLGLLPHQMQGSLSKCVLSTVLCNSSSNCIACSNAVLSEYRRRGFDFVTQAITCPTYLKDLTGISDLKKPFASKISASIPVSKTSASIPVNL  699
OsTN1t002442.1/1-1041     600 IAGTDTEHQLGLLPHQMQGSLSKCVLSTVLCNSSSNCIACSNAVLSEYRRRGFDFVTQAITCPTYLKDLTGISDLKKPFASKISASIPVSKTSASIPVNL  699

Os01t0614900-01/1-1042    701 EKLSSARCLLLGAGTLGCDVARILMDCGVRKLTVVDSGRVVVSNLARQSLYTSDDRDSPKASAILGRLKERCPSVDAKGIKMEIPMPGHPVSPNEAVSVL  800
OsTNG01t002533.1/1-1041   700 EKLSSARCLLLGAGTLGCDVARILMDCGVRKLTVVDSGRVVVSNLARQSLYTSDDRDSPKASAILGRLKERCPSVDAKGIKMEIPMPGHPVSPNEAVSVL  799
OsTN1t002442.1/1-1041     700 EKLSSARCLLLGAGTLGCDVARILMDCGVRKLTVVDSGRVVVSNLARQSLYTSDDRDSPKASAILGRLRERCPSVDAKGIKMEIPMPGHPVSPNEAVSVL  799

Os01t0614900-01/1-1042    801 EDCKRLQELVSSHDAVFLLTDTRESRWLPTLLCANENKIAITAALGYDSYLVMRHGAGPGTNCGSPDVVAAADTLSAEDVLGRQRLGCYFCNDVVAPVDS  900
OsTNG01t002533.1/1-1041   800 EDCKRLQELVSSHDAVFLLTDTRESRWLPTLLCANENKIAITAALGYDSYLVMRHGAGPGTNCGSPDVVAAADTLSAEDVLGRQRLGCYFCNDVVAPVDS  899
OsTN1t002442.1/1-1041     800 EDCKRLQELVSSHDAVFLLTDTRESRWLPTLLCANENKIAITAALGYDSYLVMRHGAGPGTNCGSPDVVAAADTLSAEDVLGRQRLGCYFCNDVVAPVDS  899

Os01t0614900-01/1-1042    901 VSNRTLDQQCTVTRPGLSSITSGCAADLFTRMLHHPDGIHAPGEIAGTSSEGPLGLLPHQIRGSLSQYNLLTLLGYSSSNCTACSNAVLSEYHRRGMDFV 1000
OsTNG01t002533.1/1-1041   900 VSNRTLDQQCTVTRPGLSSITSGCAADLFTRMLHHPDGIHAPGEIAGTSSEGPLGLLPHQIRGSLSQYNLLTLLGYSSSNCTACSNAVLSEYHRRGMDFV  999
OsTN1t002442.1/1-1041     900 VSNRTLDQQCTVTRPGLASITSGCAADLFTRMLHHPDGIHAPGEIAGTSSEGPLGLLPHQIRGSLSQYNLLTLLGYSSSNCTACSNAVLSEYRRRGMDFV  999

Os01t0614900-01/1-1042   1001 MQVINEPTYLEDLTGLTDLMKSAAYSQVEWIDEVDDDDEMDI                                                           1042
OsTNG01t002533.1/1-1041  1000 MQVINEPTYLEDLTGLTDLMKSAAYSQVEWIDEVDDDDEMDI                                                           1041
OsTN1t002442.1/1-1041    1000 MQVINEPTYLEDLTGLTDLMKSAAYSQVEWIDEVDDDDEMDI                                                           1041
```

c) *EF1*

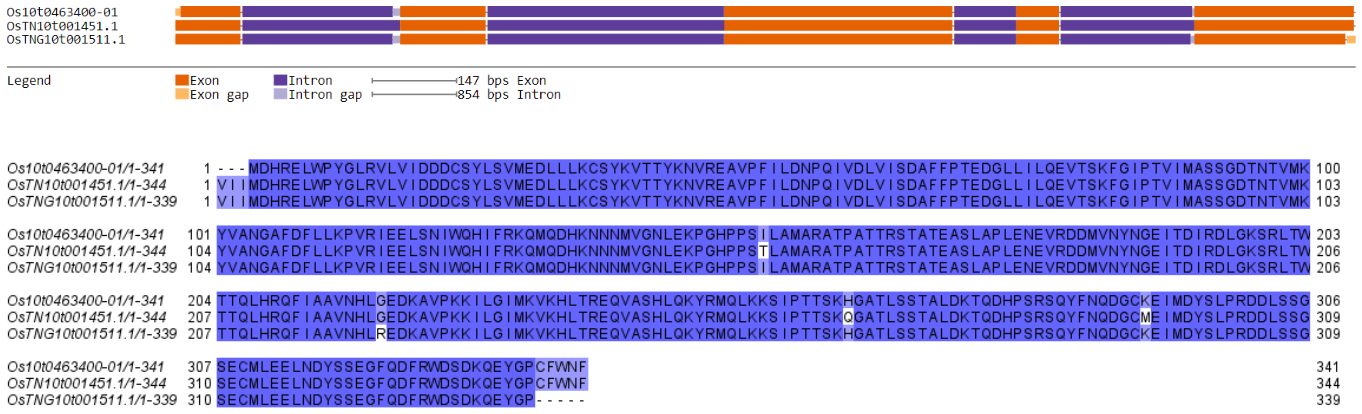


d) *OsHDT1*

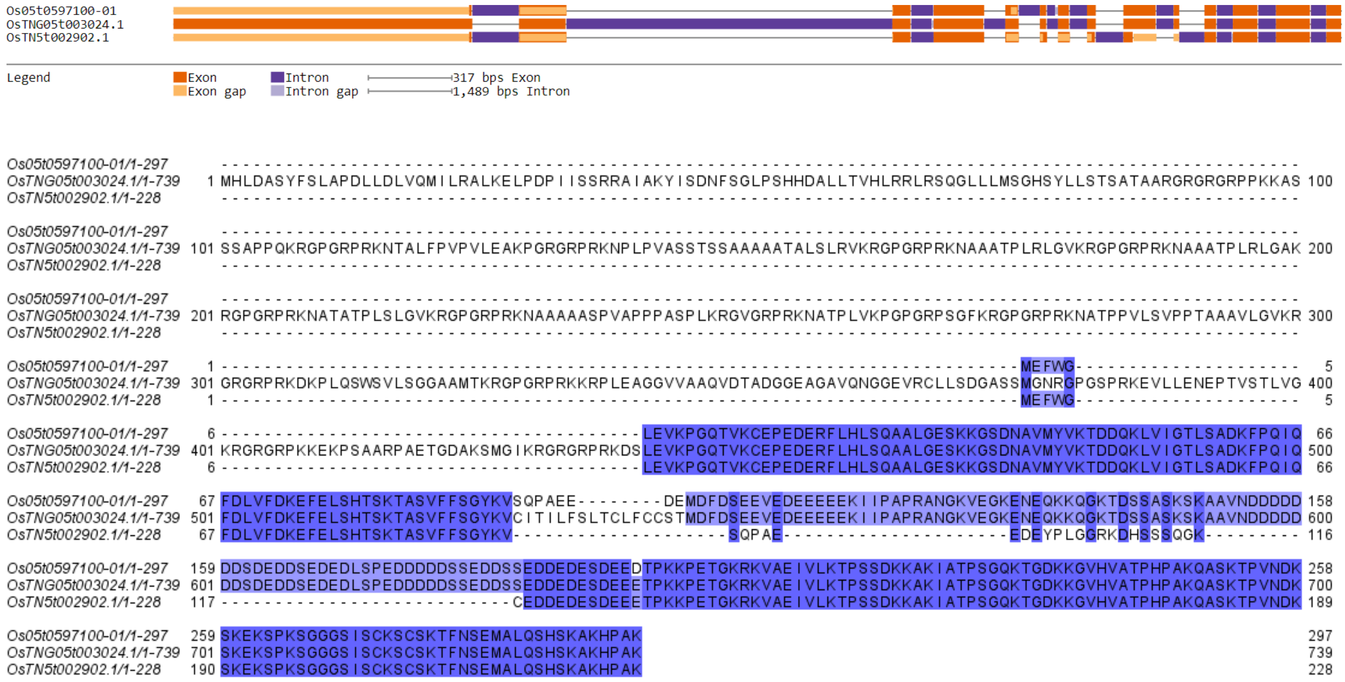


e) *DTH8*

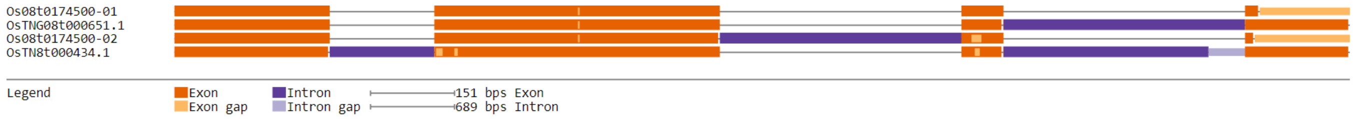

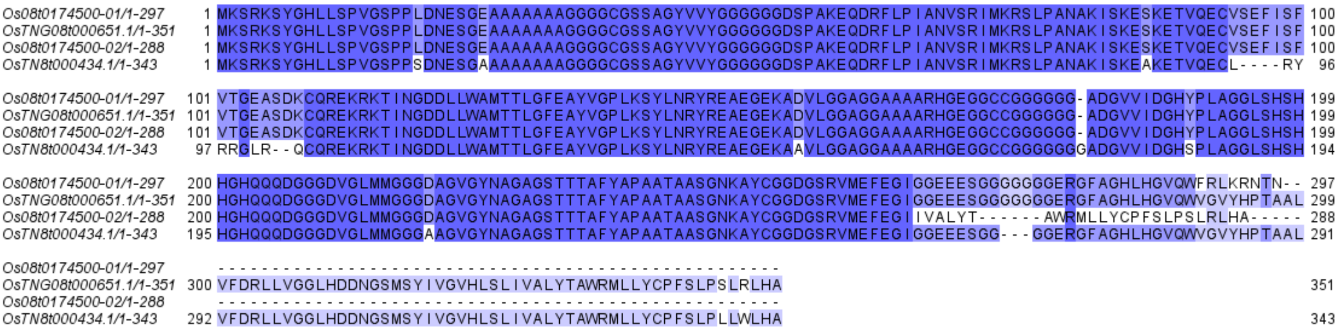

Os08t0174500-01/1-297 1 MKSRKSYGHLLSPVGSPPLDNESGEAAAAAAAGGGGCGSSAGYVVYGGGGGGDSPAKEQDRFLPIANVSRIMKRSLPANAKISKESKETVQECVSEFISF 100
OsTNG08t000651.1/1-351 1 MKSRKSYGHLLSPVGSPPLDNESGEAAAAAAAGGGGCGSSAGYVVYGGGGGGDSPAKEQDRFLPIANVSRIMKRSLPANAKISKESKETVQECVSEFISF 100
Os08t0174500-02/1-288 1 MKSRKSYGHLLSPVGSPPLDNESGEAAAAAAAGGGGCGSSAGYVVYGGGGGGDSPAKEQDRFLPIANVSRIMKRSLPANAKISKESKETVQECVSEFISF 100
OsTN8t000434.1/1-343 1 MKSRKSYGHLLSPVGSPPSDNESGAAAAAAAAGGGGCGSSAGYVVYGGGGGGDSPAKEQDRFLPIANVSRIMKRSLPANAKISKEAKETVQECL----RY 96
Os08t0174500-01/1-297 101 VTGEASDKCQREKRKTINGDDLLWAMTTLGFEAYVGPLKSYLNRYREAEGEKADVLGGAGGAAAARHGEGGCCGGGGGG-ADGVVIDGHYPLAGGLSHSH 199
OsTNG08t000651.1/1-351 101 VTGEASDKCQREKRKTINGDDLLWAMTTLGFEAYVGPLKSYLNRYREAEGEKADVLGGAGGAAAARHGEGGCCGGGGGG-ADGVVIDGHYPLAGGLSHSH 199
Os08t0174500-02/1-288 101 VTGEASDKCQREKRKTINGDDLLWAMTTLGFEAYVGPLKSYLNRYREAEGEKADVLGGAGGAAAARHGEGGCCGGGGGG-ADGVVIDGHYPLAGGLSHSH 199
OsTN8t000434.1/1-343 97 RRGLR--QCQREKRKTINGDDLLWAMTTLGFEAYVGPLKSYLNRYREAEGEKAAVLGGAGGAAAARHGEGGCCGGGGGGGADGVVIDGHSPLAGGLSHSH 194
Os08t0174500-01/1-297 200 HGHQQQDGGGDVGLMMGGGDAGVGYNAGAGSTTTAFYAPAATAASGNKAYCGGDGSRVMEFEGIGGEEESGGGGGGGERGFAGHLHGVQWFRLKRNTN-- 297
OsTNG08t000651.1/1-351 200 HGHQQQDGGGDVGLMMGGGDAGVGYNAGAGSTTTAFYAPAATAASGNKAYCGGDGSRVMEFEGIGGEEESGGGGGGGERGFAGHLHGVQWVGVYHPTAAL 299
Os08t0174500-02/1-288 200 HGHQQQDGGGDVGLMMGGGDAGVGYNAGAGSTTTAFYAPAATAASGNKAYCGGDGSRVMEFEGIIVALYT------AWRMLLYCPFSLPSLRLHA----- 288
OsTN8t000434.1/1-343 195 HGHQQQDGGGDVGLMMGGGAAGVGYNAGAGSTTTAFYAPAATAASGNKAYCGGDGSRVMEFEGIGGEEESGG---GGERGFAGHLHGVQWVGVYHPTAAL 291
Os08t0174500-01/1-297
OsTNG08t000651.1/1-351 300 VFDRLLVGGLHDDNGSMSYIVGVHLSLIVALYTAWRMLLYCPFSLPSLRLHA 351
Os08t0174500-02/1-288
OsTN8t000434.1/1-343 292 VFDRLLVGGLHDDNGSMSYIVGVHLSLIVALYTAWRMLLYCPFSLPLLWLHA 343


## f) *Hd1*

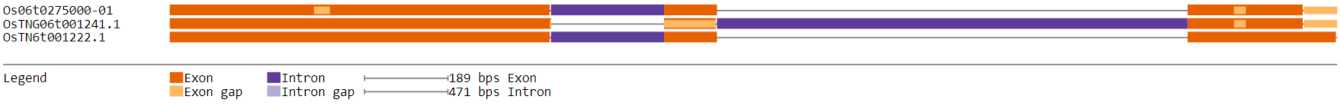

Os06t0275000-01
OsTNG06t001241.1
OsTN6t001222.1
Legend
Exon
Exon gap
Intron
Intron gap
189 bps Exon
471 bps Intron


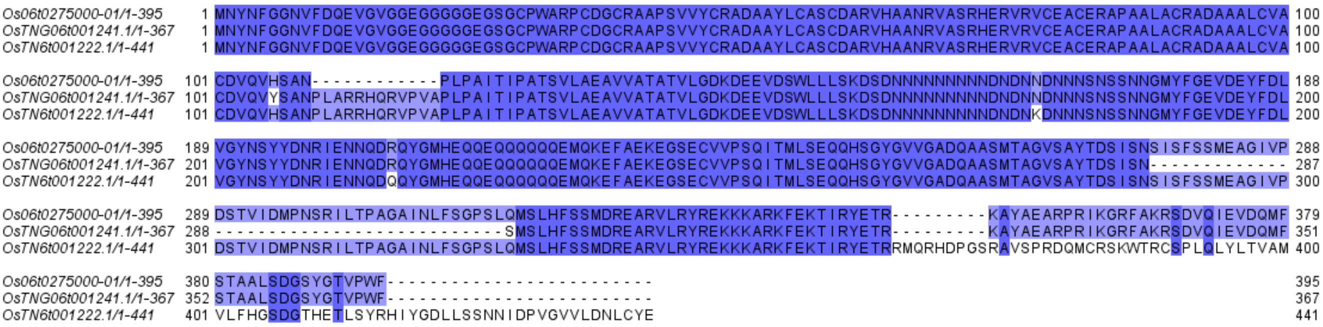

Os06t0275000-01/1-395 1 MNYNFGGNVFDQEVGVGGEGGGGGEGSGCPWARPCDGCRAAPSVVYCRADAAYLCASCDARVHAANRVASRHERVRVCEACERAPAALACRADAAALCVA 100
OsTNG06t001241.1/1-367 1 MNYNFGGNVFDQEVGVGGEGGGGGEGSGCPWARPCDGCRAAPSVVYCRADAAYLCASCDARVHAANRVASRHERVRVCEACERAPAALACRADAAALCVA 100
OsTN6t001222.1/1-441 1 MNYNFGGNVFDQEVGVGGEGGGGGEGSGCPWARPCDGCRAAPSVVYCRADAAYLCASCDARVHAANRVASRHERVRVCEACERAPAALACRADAAALCVA 100
Os06t0275000-01/1-395 101 CDVQVHSAN------------PLPAITIPATSVLAEAVVATATVLGDKDEEVDSWLLLSKDSDNNNNNNNNNDNDNNDNNNSNSSNNGMYFGEVDEYFDL 188
OsTNG06t001241.1/1-367 101 CDVQVYSANPLARRHQRVPVAPLPAITIPATSVLAEAVVATATVLGDKDEEVDSWLLLSKDSDNNNNNNNNNDNDNNDNNNSNSSNNGMYFGEVDEYFDL 200
OsTN6t001222.1/1-441 101 CDVQVHSANPLARRHQRVPVAPLPAITIPATSVLAEAVVATATVLGDKDEEVDSWLLLSKDSDNNNNNNNNNDNDNKDNNNSNSSNNGMYFGEVDEYFDL 200
Os06t0275000-01/1-395 189 VGYNSYYDNRIENNQDRQYGMHEQQEQQQQQEMQKEFAEKEGSECVVPSQITMLSEQQHSGYGVVGADQAASMTAGVSAYTDSISNSISFSSMEAGIVP 288
OsTNG06t001241.1/1-367 201 VGYNSYYDNRIENNQDRQYGMHEQQEQQQQQEMQKEFAEKEGSECVVPSQITMLSEQQHSGYGVVGADQAASMTAGVSAYTDSISN------------- 287
OsTN6t001222.1/1-441 201 VGYNSYYDNRIENNQDQQYGMHEQQEQQQQQEMQKEFAEKEGSECVVPSQITMLSEQQHSGYGVVGADQAASMTAGVSAYTDSISNSISFSSMEAGIVP 300
Os06t0275000-01/1-395 289 DSTVIDMPNSRILTPAGAINLFSGPSLQMSLHFSSMDREARVLRYREKKKARKFEKTIRYETR---------KAYAEARPRIKGRFAKRSDVQIEVDQMF 379
OsTNG06t001241.1/1-367 288 ---------------------------SMSLHFSSMDREARVLRYREKKKARKFEKTIRYETR---------KAYAEARPRIKGRFAKRSDVQIEVDQMF 351
OsTN6t001222.1/1-441 301 DSTVIDMPNSRILTPAGAINLFSGPSLQMSLHFSSMDREARVLRYREKKKARKFEKTIRYETRRMQRHDPGSRAVSPRDQMCRSKWTRCSPLQLYLTVAM 400
Os06t0275000-01/1-395 380 STAALSDGSYGTVPWF------------------------ 395
OsTNG06t001241.1/1-367 352 STAALSDGSYGTVPWF------------------------ 367
OsTN6t001222.1/1-441 401 VLFHGSDGTHETLSYRHIYGDLLSSNNIDPVGVVLDNLCYE 441


### g) *Ghd8*

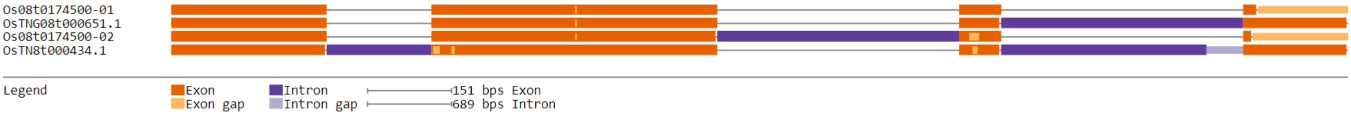

Os08t0174500-01
OsTNG08t000651.1
Os08t0174500-02
OsTN8t000434.1
Legend
Exon
Exon gap
Intron
Intron gap
151 bps Exon
689 bps Intron


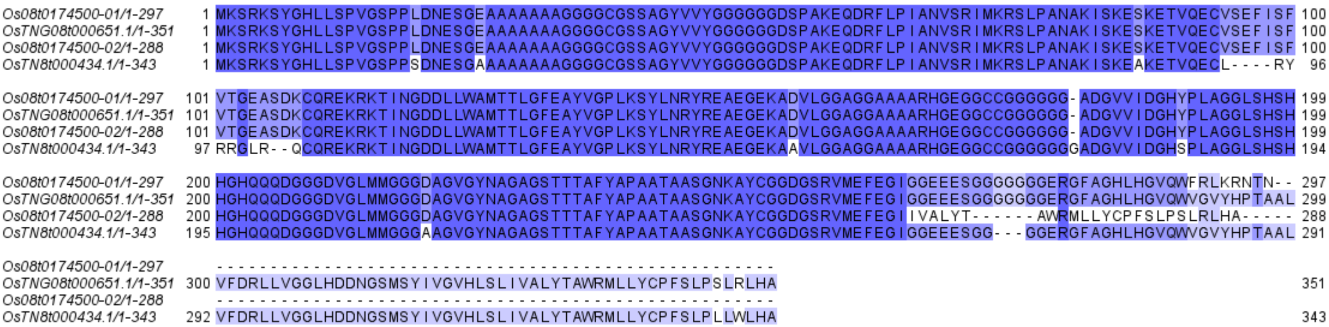

Os08t0174500-01/1-297 1 MKSRKSYGHLLSPVGSPPLDNESGEAAAAAAAGGGGCGSSAGYVVYGGGGGGDSPAKEQDRFLPIANVSRIMKRSLPANAKISKESKETVQECVSEFISF 100
OsTNG08t000651.1/1-351 1 MKSRKSYGHLLSPVGSPPLDNESGEAAAAAAAGGGGCGSSAGYVVYGGGGGGDSPAKEQDRFLPIANVSRIMKRSLPANAKISKESKETVQECVSEFISF 100
Os08t0174500-02/1-288 1 MKSRKSYGHLLSPVGSPPLDNESGEAAAAAAAGGGGCGSSAGYVVYGGGGGGDSPAKEQDRFLPIANVSRIMKRSLPANAKISKESKETVQECVSEFISF 100
OsTN8t000434.1/1-343 1 MKSRKSYGHLLSPVGSPPSDNESGAAAAAAAAGGGGCGSSAGYVVYGGGGGGDSPAKEQDRFLPIANVSRIMKRSLPANAKISKEAKETVQECL----RY 96
Os08t0174500-01/1-297 101 VTGEASDKCQREKRKTINGDDLLWAMTTLGFEAYVGPLKSYLNRYREAEGEKADVLGGAGGAAAARHGEGGCCGGGGGG-ADGVVIDGHYPLAGGLSHSH 199
OsTNG08t000651.1/1-351 101 VTGEASDKCQREKRKTINGDDLLWAMTTLGFEAYVGPLKSYLNRYREAEGEKADVLGGAGGAAAARHGEGGCCGGGGGG-ADGVVIDGHYPLAGGLSHSH 199
Os08t0174500-02/1-288 101 VTGEASDKCQREKRKTINGDDLLWAMTTLGFEAYVGPLKSYLNRYREAEGEKADVLGGAGGAAAARHGEGGCCGGGGGG-ADGVVIDGHYPLAGGLSHSH 199
OsTN8t000434.1/1-343 97 RRGLR--QCQREKRKTINGDDLLWAMTTLGFEAYVGPLKSYLNRYREAEGEKAAVLGGAGGAAAARHGEGGCCGGGGGGGADGVVIDGHSPLAGGLSHSH 194
Os08t0174500-01/1-297 200 HGHQQQDGGGDVGLMMGGGDAGVGYNAGAGSTTTAFYAPAATAASGNKAYCGGDGSRVMEFEGIGGEEESGGGGGGGERGFAGHLHGVQWFRLKRNTN-- 297
OsTNG08t000651.1/1-351 200 HGHQQQDGGGDVGLMMGGGDAGVGYNAGAGSTTTAFYAPAATAASGNKAYCGGDGSRVMEFEGIGGEEESGGGGGGGERGFAGHLHGVQWVGVYHPTAAL 299
Os08t0174500-02/1-288 200 HGHQQQDGGGDVGLMMGGGDAGVGYNAGAGSTTTAFYAPAATAASGNKAYCGGDGSRVMEFEGIIVALYT------AWRMLLYCPFSLPSLRLHA----- 288
OsTN8t000434.1/1-343 195 HGHQQQDGGGDVGLMMGGGAAGVGYNAGAGSTTTAFYAPAATAASGNKAYCGGDGSRVMEFEGIGGEEESGG---GGERGFAGHLHGVQWVGVYHPTAAL 291
Os08t0174500-01/1-297
OsTNG08t000651.1/1-351 300 VFDRLLVGGLHDDNGSMSYIVGVHLSLIVALYTAWRMLLYCPFSLPSLRLHA 351
Os08t0174500-02/1-288
OsTN8t000434.1/1-343 292 VFDRLLVGGLHDDNGSMSYIVGVHLSLIVALYTAWRMLLYCPFSLPLLWLHA 343

h) *OsTrx1*

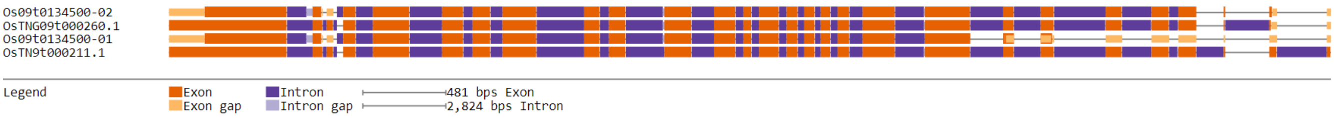


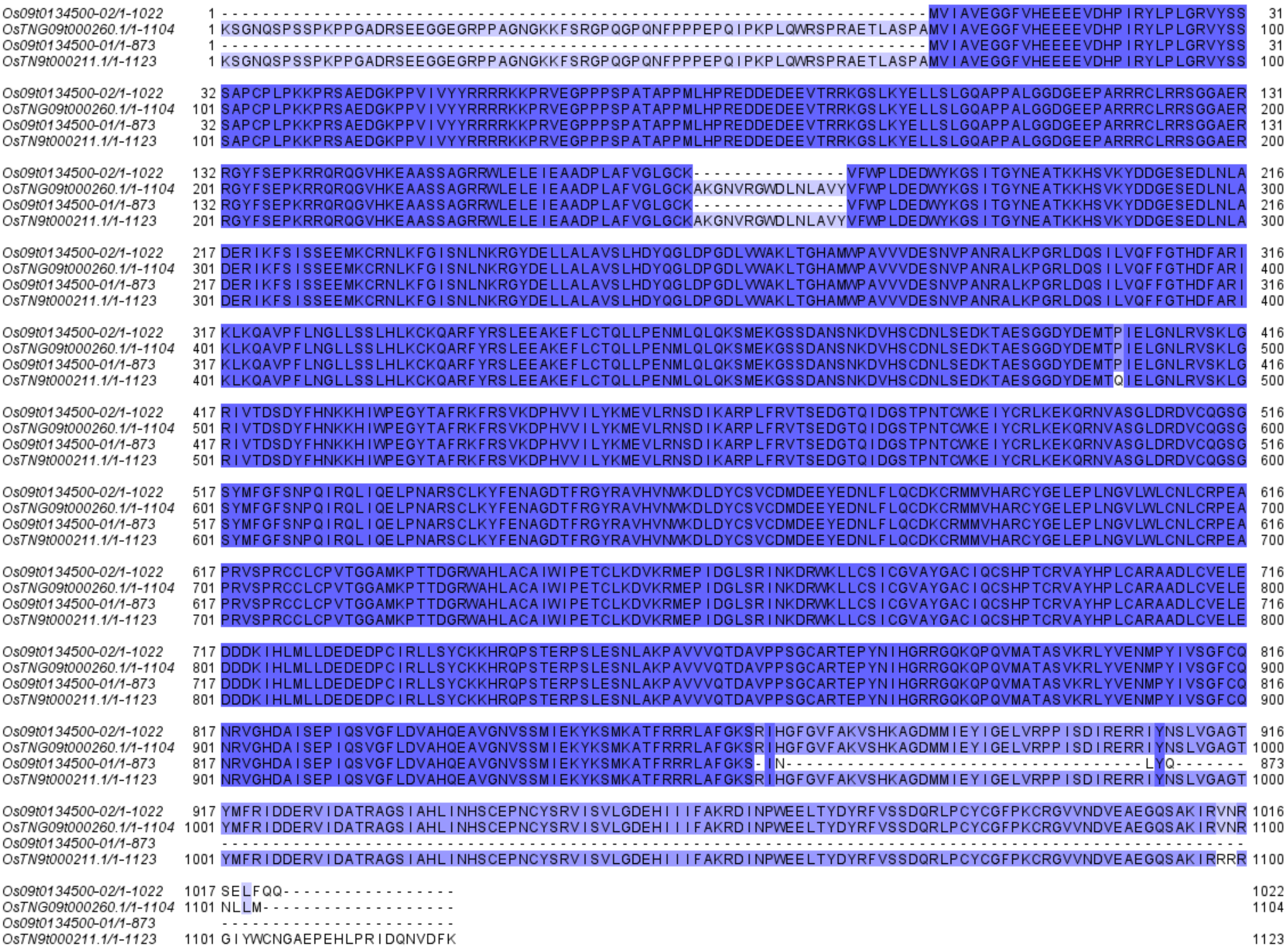


i) *PHYB*

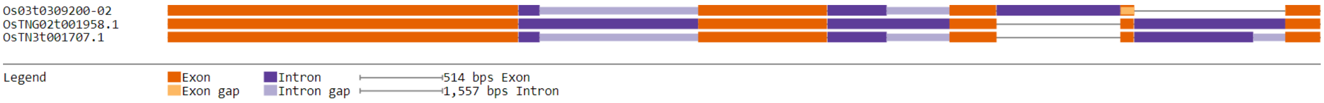

```
Os03t0309200-02/1-1171      1 MASGSRATPTRSPSSARPAAPRHQHHHSQSSGGSTSRAGGGGGGGGGGGGGGAAAAESVSKAVAQYTLDARLHAVFEQSGASGRSFDYTQSLRASPTPSSE  100
OsTNG02t001958.1/1-1200     1 MASGSRATPTRSPSSARPAAPRHQHHHSQSSGGSTSRAGGGGGGGGGGGGGGAAAAESVSKAVAQYTLDARLHAVFEQSGASGRSFDYTQSLRASPTPSSE  100
OsTN3t001707.1/1-1200       1 MASGSRATPTRSPSSARPAAPRHQHHHSQSSGGSTSRAGGGGGGGGGGGGGGAAAAESVSKAVAQYTLDARLHAVFEQSGASGRSFDYTQSLRASPTPSSE  100

Os03t0309200-02/1-1171    101 QQIAAYLSRIQRGGHIQPFGCTLAVADDSSFRLLAYSENTADLLDLSPHHSVPSLDSSAVPPPVSLGADARLLFAPSSAVLLERAFAAREISLLNPLWIH  200
OsTNG02t001958.1/1-1200   101 QQIAAYLSRIQRGGHIQPFGCTLAVADDSSFRLLAYSENTADLLDLSPHHSVPSLDSSAVPPPVSLGADARLLFAPSSAVLLERAFAAREISLLNPLWIH  200
OsTN3t001707.1/1-1200     101 QQIAAYLSRIQRGGHIQPFGCTLAVADDSSFRLLAYSENTADLLDLSPHHSVPSLDSSAVPPPVSLGADARLLFAPSSAVLLERAFAAREISLLNPLWIH  200

Os03t0309200-02/1-1171    201 SRVSSKPFYAILHRIDVGVVIDLEPARTEDPALSIAGAVQSQKLAVRAISRLQALPGGDVKLLCDTVVEYVRELTGYDRVMVYRFHEDEHGEVVAESRRN  300
OsTNG02t001958.1/1-1200   201 SRVSSKPFYAILHRIDVGVVIDLEPARTEDPALSIAGAVQSQKLAVRAISRLQALPGGDVKLLCDTVVEYVRELTGYDRVMVYRFHEDEHGEVVAESRRN  300
OsTN3t001707.1/1-1200     201 SRVSSKPFYAILHRIDVGVVIDLEPARTEDPALSIAGAVQSQKLAVRAISRLQALPGGDVKLLCDTVVEHVRELTGYDRVMVYRFHEDEHGEVVAESRRS  300

Os03t0309200-02/1-1171    301 NLEPYIGLHYPATDIPQASRFLFRQNRVRMIADCHAAPVRVIQDPALTQPLCLVGSTLRSPHGCHAQYMANMGSIASLVMAVIISSGGDDDHNISRGSIP  400
OsTNG02t001958.1/1-1200   301 NLEPYIGLHYPATDIPQASRFLFRQNRVRMIADCHAAPVRVIQDPALTQPLCLVGSTLRSPHGCHAQYMANMGSIASLVMAVIISSGGDDDHNISRGSIP  400
OsTN3t001707.1/1-1200     301 NLEPYIGLHYPATDIPQASRFLFRQNRVRMIADCHAAPVRVIQDPALTQPLCLVGSTLRSPHGCHAQYMANMGSIASLVMAVIISSGGDDDHNIARGSIP  400

Os03t0309200-02/1-1171    401 SAMKLWGLVVCHHTSPRCIPFPLRYACEFLMQAFGLQLNMELQLAHQLSEKHILRTQTLLCDMLLRDSPTGIVTQSPSIMDLVKCDGAALYYHGKYYPLG  500
OsTNG02t001958.1/1-1200   401 SAMKLWGLVVCHHTSPRCIPFPLRYACEFLMQAFGLQLNMELQLAHQLSEKHILRTQTLLCDMLLRDSPTGIVTQSPSIMDLVKCDGAALYYHGKYYPLG  500
OsTN3t001707.1/1-1200     401 SAMKLWGLVVCHHTSPRCIPFPLRYACEFLMQAFGLQLNMELQLAHQLSEKHILRTQTLLCDMLLRDSPTGIVTQSPSIMDLVKCDGAALYYHGKYYPLG  500

Os03t0309200-02/1-1171    501 VTPTEVQIKDIIEWLTMCHGDSTGLSTDSLADAGYPGAAALGDAVSGMAVAYITPSDYLFWFRSHTAKEIKWGGAKHHPEDKDDGQRMHPRSSFKAFLEV  600
OsTNG02t001958.1/1-1200   501 VTPTEVQIKDIIEWLTMCHGDSTGLSTDSLADAGYPGAAALGDAVSGMAVAYITPSDYLFWFRSHTAKEIKWGGAKHHPEDKDDGQRMHPRSSFKAFLEV  600
OsTN3t001707.1/1-1200     501 VTPTEVQIKDIIEWLTMCHGDSTGLSTDSLADAGYSGAAALGDAVSGMAVAYITPSDYLFWFRSHTAKEIKWGGAKHHPEDKDDGQRMHPRSSFKAFLEV  600

Os03t0309200-02/1-1171    601 VKSRSLPWENAEMDAIHSLQLILRDSFRDSAEGTSNSKAIVNGQVQLGELELRGIDELSSVAREMVRLIETATVPIFAVDTDGCINGWNAKVAELTGLSV  700
OsTNG02t001958.1/1-1200   601 VKSRSLPWENAEMDAIHSLQLILRDSFRDSAEGTSNSKAIVNGQVQLGELELRGIDELSSVAREMVRLIETATVPIFAVDTDGCINGWNAKVAELTGLSV  700
OsTN3t001707.1/1-1200     601 VKSRSLPWENAEMDAIHSLQLILRDSFRDSAEGTSNSKAIVNGQVQLGELELRGIDELSSVAREMVRLIETATVPIFAVDTDGCINGWNAKVAELTGLSV  700

Os03t0309200-02/1-1171    701 EEAMGKSLVNDLIFKESEETVNKLLSRALRGDEDKNVEIKLKTFGPEQSKGPIFVIVNACSSRDYTKNIVGVCFVGQDVTGQKVVMDKFINIQGDYKAIV  800
OsTNG02t001958.1/1-1200   701 EEAMGKSLVNDLIFKESEETVNKLLSRALRGDEDKNVEIKLKTFGPEQSKGPIFVIVNACSSRDYTKNIVGVCFVGQDVTGQKVVMDKFINIQGDYKAIV  800
OsTN3t001707.1/1-1200     701 EEAMGKSLVNDLIFKESEETVNKLLSRALRGDEDKNVEIKLKTFGPEQSKGPIFVIVNACSSRDYTKNIVGVCFVGQDVTGQKVVMDKFINIQGDYKAIV  800

Os03t0309200-02/1-1171    801 HNPNPLIPPIFASDENTCCSEWNTAMEKLTGWSRGEVVGKLLVGEVFGNCCRLKGPDALTKFMIVLHNAIGGQDCEKFPFSFFDKNGKYVQALLTANTRS  900
OsTNG02t001958.1/1-1200   801 HNPNPLIPPIFASDENTCCSEWNTAMEKLTGWSRGEVVGKLLVGEVFGNCCRLKGPDALTKFMIVLHNAIGGQDCEKFPFSFFDKNGKYVQALLTANTRS  900
OsTN3t001707.1/1-1200     801 HNPNPLIPPIFASDENTCCSEWNTAMEKLTGWSRGEVVGKLLVGEVFGNCCRLKGPDALTKFMIVLHNAIGGQDCEKFPFSFFDKNGKYVQALLTANTRS  900

Os03t0309200-02/1-1171    901 RMDGEAIGAFCFLQIASPELQQAFEIQRHHEKKCYARMKELAYIYQEIKNPLNGIRFTNSLLEMTDLKDDQRQFLETSTACEKQMSKIVKDASLQSIEDG 1000
OsTNG02t001958.1/1-1200   901 RMDGEAIGAFCFLQIASPELQQAFEIQRHHEKKCYARMKELAYIYQEIKNPLNGIRFTNSLLEMTDLKDDQRQFLETSTACEKQMSKIVKDASLQSIEDG 1000
OsTN3t001707.1/1-1200     901 RMDGEAIGAFCFLQIASPELQQAFEIQRHHEKKCYARMKELAYIYQEIKNPLNGIRFTNSLLEMTDLKDDQRQFLETSTACEKQMSKIVKDASLQSIEDG 1000

Os03t0309200-02/1-1171   1001 SLVLEKGEFSLGSVMNAVVSQVMIQLRERDLQLIRDIPDEIKEASAYGDQYRIQQVLCDFLLSMVRFAPAENGWVEIQVRPNIKQNSDGTDTMLFLFR-- 1098
OsTNG02t001958.1/1-1200  1001 SLVLEKGEFSLGSVMNAVVSQVMIQLRERDLQLIRDIPDEIKEASAYGDQYRIQQVLCDFLLSMVRFAPAENGWVEIQVRPNIKQNSDGTDTMLFLFRLA 1100
OsTN3t001707.1/1-1200    1001 SLVLEKGEFSLGSVMNAVVSQVMIQLRERDLQLIRDIPDEIKEASAYGDQYRIQQVLCDFLLSMVRFAPAENGWVEIQVRPNIKQNSDGTDTMLFLFRLA 1100

Os03t0309200-02/1-1171   1099 ---------------------------FACPGEGLPPEIVQDMFSNSRWTTQEGIGLSICRKILKLMGGEVQYIRESERSFFHIVLELPQPQQAASRGTS 1171
OsTNG02t001958.1/1-1200  1101 IYLHFQYQKAIHILTQEFLVLNLVEDKFACPGEGLPPEIVQDMFSNSRWTTQEGIGLSICRKILKLMGGEVQYIRESERSFFHIVLELPQPQQAASRGTS 1200
OsTN3t001707.1/1-1200    1101 IYLHFQYQKAIHILTQEFLVLNLVEDKFACPGEGLPPEIVQDMFSNSRWTTQEGIGLSICRKILKLMGGEVQYIRESERSFFHIVLELPQPQQAASRGTS 1200
```

j) *NRRa/NRRb*

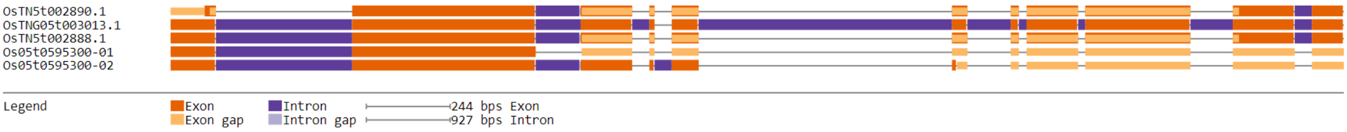


```
OsTN5t002890.1/1-269       1 ---------------------------------LSILQ------GNMAQPSFLSEYDLGGEGDLFKAPEPIIEEPVLSLDPVAAAISMMSGSENVMDETI  61
OsTNG05t003013.1/1-569     1 MYAAMYPETFGFSAYPQQQQPPPDAASCIYTTALPLIADPPDILGNMAQPSFLSEYDLGGEGDLFKAPEPIIEEPVLSLDPVAAAISMMSGSENVMDETI 100
OsTN5t002888.1/1-308       1 MYAAMYPETFGFSAYPQQQQPPPDAASCIYTTALPLIADPPDILGNMAQPSFLSEYDLGGEGDLFKAPEPIIEEPVLSLDPVAAAISMMSGSENVMDETI 100
Os05t0595300-01/1-223      1 MYAAMYPETFGFSAYPQQQQPPPDAASCIYTTALPLIADPPDILGNMAQPSFLSEYDLGGEGDLFKAPEPIIEEPVLSLDPVAAAISMMSGSENVMDETI 100
Os05t0595300-02/1-308      1 MYAAMYPETFGFSAYPQQQQPPPDAASCIYTTALPLIADPPDILGNMAQPSFLSEYDLGGEGDLFKAPEPIIEEPVLSLDPVAAAISMMSGSENVMDETI 100

OsTN5t002890.1/1-269      62 EVADISDIQNDSLLSEVLYECEKELMEKSAIEETISELLDVKIPMLQVEEFPRETQVQLPAMEKEKPSVPECCSLQKSVSSGCLNSADWINGPARPNFLD 161
OsTNG05t003013.1/1-569   101 EVADISDIQNDSLLSEVLYECEKELMEKSAIEETISELLDVKIPMLQVEEFPRETQVQLPAMEKEKPSVPECCSLQKSVSSGCLNSADWINGPARPNFLD 200
OsTN5t002888.1/1-308     101 EVADISDIQNDSLLSEVLYECEKELMEKSAIEETISELLDVKIPMLQVEEFPRETQVQLPAMEKEKPSVPECCSLQKSVSSGCLNSADWINGPARPNFLD 200
Os05t0595300-01/1-223    101 EVADISDIQNDSLLSEVLYECEKELMEKSAIEETISELLDVKIPMLQVEEFPRETQVQLPAMEKEKPSVPECCSLQKSVSSGCLNSADWINGPARPNFLD 200
Os05t0595300-02/1-308    101 EVADISDIQNDSLLSEVLYECEKELMEKSAIEETISELLDVKIPMLQVEEFPRETQVQLPAMEKEKPSVPECCSLQKSVSSGCLNSADWINGPARPNFLD 200

OsTN5t002890.1/1-269     162 FQGLDFETAFGLRRAYSEGDIQN----------------------------------------------------------------------------- 184
OsTNG05t003013.1/1-569   201 FQGLDFETAFGLRRAYSEGDIQHPRPGNSGNAQLASCERLVTISDLKSEERKQKLSRYRKKKVKRNFGRKIKYACRKALADSQPRVRGRFAKIEEGDLLK 300
OsTN5t002888.1/1-308     201 FQGLDFETAFGLRRAYSEGDIQN----------------------------------------------------------------------------- 223
Os05t0595300-01/1-223    201 FQGLDFETAFGLRRAYSEGDIQV----------------------------------------------------------------------------- 223
Os05t0595300-02/1-308    201 FQGLDFETAFGLRRAYSEGDIQNLGASTPRPGNSGNAQLASCERLVTISDLKSEERKQKLSRYRKKKVKRNFGRKIKYACRKALADSQPRVRGRFAKIEE 300

OsTN5t002890.1/1-269         ----------------------------------------------------------------------------------------------------
OsTNG05t003013.1/1-569   301 PRKYYQAPLDAVRLDDRDLNMFAIKNGNMAQPSFLSEYDLGGEGDLFKAPEPIIEEPVLSLDPVAAAISMMSGSENCEKELMEKSAIEETISELLDVKIP 400
OsTN5t002888.1/1-308         ----------------------------------------------------------------------------------------------------
Os05t0595300-01/1-223        ----------------------------------------------------------------------------------------------------
Os05t0595300-02/1-308    301 GDLLKPRK-------------------------------------------------------------------------------------------- 308

OsTN5t002890.1/1-269     185 ------------------------------------------------------------------------------------LGASTPRPGNSGNAQL 200
OsTNG05t003013.1/1-569   401 MLQVEEFPRETQVQLPAMEKEKPSVPECCSLQKSVSSGCLNSADWINGPARPNFLDFQGLDFETAFGLRRAYSEGDIQLSFQQNLGASTPRPGNSGNAQL 500
OsTN5t002888.1/1-308     224 ------------------------------------------------------------------------------------LGASTPRPGNSGNAQL 239
Os05t0595300-01/1-223        ----------------------------------------------------------------------------------------------------
Os05t0595300-02/1-308        ----------------------------------------------------------------------------------------------------

OsTN5t002890.1/1-269     201 ASCERLVTISDLKSEERKQKLSRYRKKKVKRNFGRKIKYACRKALADSQPRVRGRFAKIEEGDLLKPRK                                269
OsTNG05t003013.1/1-569   501 ASCERLVTISDLKSEERKQKLSRYRKKKVKRNFGRKIKYACRKALADSQPRVRGRFAKIEEGDLLKPRK                                569
OsTN5t002888.1/1-308     240 ASCERLVTISDLKSEERKQKLSRYRKKKVKRNFGRKIKYACRKALADSQPRVRGRFAKIEEGDLLKPRK                                308
Os05t0595300-01/1-223        ---------------------------------------------------------------------
Os05t0595300-02/1-308        ---------------------------------------------------------------------
```

**Figure S9. Gene structure diagrams and protein sequence alignment of photoperiod genes of TNG67 that are hybrid.** a) *OsELF3-1/EF7*, b) *OsATG7*, c) *EF1*, d) *OsHDT1*, e) *DTH8*, f) *Hd1*, g) *Ghd8*, h) *OsTrx1*, i) *PHYB* and j) *NRRa/NRRb*
